# Does social antagonism facilitate supergene expansion? A novel region of suppressed recombination in a 4-haplotype supergene system

**DOI:** 10.1101/2023.03.29.534793

**Authors:** Giulia Scarparo, Marie Palanchon, Alan Brelsford, Jessica Purcell

## Abstract

Models of both sex chromosome evolution and the genetic basis of local adaptation suggest that selection acts to lock beneficial combinations of alleles together in regions of reduced or suppressed recombination. Drawing inspiration from such models, we apply similar logic to investigate whether an autosomal supergene underlying colony social organization in ants expanded to include “socially antagonistic” alleles. We tested this premise in a *Formica* ant species wherein we identified four supergene haplotypes on chromosome 3 underlying colony social organization and sex ratio. Remarkably, we discovered a novel rearranged supergene variant (9r) on chromosome 9 underlying queen miniaturization. The 9r is tightly linked to one of the haplotypes (P_2_) on chromosome 3, found predominantly in multi-queen (polygyne) colonies. We suggest that queen miniaturization is strongly disfavored in the single queen (monogyne) background, and thus socially antagonistic. As such, divergent selection experienced by ants living in alternative social ‘environments’ (monogyne and polygyne) may have contributed to the emergence of a genetic polymorphism on chromosome 9 and associated queen size dimorphism. Consequently, an ancestral polygyne-associated haplotype may have expanded to include the polymorphism on chromosome 9, resulting in a larger region of suppressed recombination spanning two chromosomes. This process is analogous to the formation of neo-sex chromosomes and consistent with models of expanding regions of suppressed recombination. We also propose that miniaturized queens, 16-20% smaller than queens without 9r, could be incipient intraspecific social parasites.

**Significance statement:** When sets of gene variants work well together, selection may lead to a reduction in recombination between them. Here, we discover a novel supergene region on chromosome 9 that controls a previously undescribed queen size polymorphism in *Formica cinerea* ants. The haplotype that is found in small queens, 9r, is tightly linked to a supergene haplotype on chromosome 3 that is found in multi-queen colonies. We propose that the region of suppressed recombination expanded to include both chromosome 3 and chromosome 9 because small queens could be successful in the multi-queen but not in the single-queen environment.

## Introduction

The field of sex chromosome evolution is facing a paradigm shift (Berdan et al. 2022). A long-standing model suggests that sex chromosomes initially form and expand through linkage between sex-determining genes and alleles at other genes that are favorable in one sex but detrimental in the other (Charlesworth and Charlesworth 1978; Charlesworth et al. 2005). However, the proposed role of linkage between a sex determining gene and sexually antagonistic loci lacks support in numerous empirical systems (Furman et al. 2020; Perrin 2021; Renner and Müller 2021). New models propose alternative combinations of selective forces not involving linked antagonistic loci that could trigger the formation or expansion of incipient sex chromosomes (Lenormand and Roze 2022; Jay et al. 2022). Research on autosomal supergenes, regions of the genome that control other complex traits and exhibit suppressed recombination between multiple genes, has drawn substantial inspiration from the sex chromosome evolution literature (e.g. Wang et al. 2013, Tuttle et al. 2016, Branco et al. 2018). Taking these comparisons a step further, autosomal supergenes should present an alternative avenue to test the predictions of classical and emerging models of sex-chromosome evolution (e.g. Branco et al. 2017; Branco et al. 2018; Duhamel et al. 2022).

The importance of reduced or suppressed recombination has been widely recognized in contexts beyond sex chromosome evolution, such as in the emergence of local adaptation (e.g. Charlesworth and Charlesworth 1979; Kirkpatrick and Barton 2006). Yeaman (2013) applies similar logic in a model predicting that alleles co-adapted to different environments in divergent populations eventually rearrange in physically close and tightly linked clusters of loci. Recent empirical evidence supports the idea that regions of the genome that are highly differentiated between ecotypes also exhibit elevated rates of rearrangement. For example, in a comparison of marine and freshwater sticklebacks, Li et al. (2022) showed that genomic regions of high differentiation were also hotspots of rearrangement, suggesting that ecological selection can shape genetic architecture. Autosomal supergenes that shape additional ‘environments’, such as social context, mating type, or life history strategy, offer an opportunity to seek independent support for models of genetic linkage. Specifically, if supergenes form or expand to link favorable combinations of alleles, this would be consistent with both the Charlesworth and Yeaman models, extending them to novel contexts.

During the past decade, the discovery of autosomal supergenes underlying complex mating systems (Branco et al. 2018), migratory behavior (Kess et al. 2019), mimetic coloration (Joron et al. 2011), and social organization (Wang et al. 2013; Purcell et al. 2014), among other complex traits, has rapidly accelerated. As in sex chromosomes, alternative phenotypes controlled by supergene variants lead to different trait optima. Here, we propose that alternative social contexts shaped by a supergene could favor the expansion of regions of suppressed recombination to include “socially antagonistic” loci. We define socially antagonistic loci as alleles that have beneficial fitness outcomes in one social environment, but detrimental outcomes in the other social environment (see also Martinez-Ruiz et al. 2020; Chapuisat 2023). We examine this idea in a supergene system in ants with (at least) three known alternative supergene haplotypes (Brelsford et al. 2020), in a relatively ancient autosomal supergene that likely emerged at least 23 M years ago (Purcell et al. 2021).

The *Formica* social supergene was initially described in the Alpine silver ant *Formica selysi* (Purcell et al. 2014). Alternative haplotypes of the supergene are associated with colony queen number, thus determining whether a colony is monogyne (with only one queen) or polygyne (with two or more queens). In *Formica* ants (and other socially polymorphic ants), a suite of other traits is frequently associated with variation in colony queen number, including body size of queens and workers (Sundström 1995a; Schwander et al. 2005; Rosset and Chapuisat 2007), colony size (Rosset and Chapuisat 2007), dispersal probability (Sundström 1995a; Fontcuberta et al. 2022), and investment in sexual offspring (Sundström 1995b; Rosset and Chapuisat 2006). In *F. selysi*, monogyne colonies exclusively harbor individuals carrying the monogyne-associated haplotype, M, whereas polygyne colonies always contain individuals bearing at least one copy of the alternative polygyne-associated haplotype, P (Purcell et al. 2014; Avril et al. 2019). The P haplotype acts as a maternal-effect killer, causing the early death of any offspring of heterozygous mothers that do not bear the P haplotype (Avril et al. 2020). Recently, Tafreshi et al. (2022) proposed that this polymorphism is only stable in the presence of both assortative mating and large fitness differences between supergene genotypes, both of which have recent empirical support (Avril et al. 2019; Blacher et al. 2023).

Although most of the ant species with a known supergene have only two alternative haplotypes (reviewed by Kay et al. 2022; Chapuisat 2023), a third haplotype associated with split sex ratio phenotypes (i.e., whether colonies specialize in producing males or gynes) was recently identified in *Formica glacialis* and *Formica podzolica* (Lagunas-Robles et al. 2021). In 2020, Brelsford and coauthors analyzing a limited number of individuals found two other species (*F. cinerea* and *F. lemani*) having three haplotypes, although they did not characterize their functions. Here, we investigate the supergene variation, both in terms of genetic architecture and phenotype, in *F. cinerea*. We ask whether autosomal supergene variants expand to include genes or chromosomal regions that encode socially antagonistic alleles, consistent with the classical sex chromosome evolution model.

## Results

### Variants of the supergene

As expected based on a preliminary assessment of genetic variation on the *F. cinerea* supergene (Brelsford et al. 2020), we detected more than two haplotypes of the social supergene in *F. cinerea*. A PCA of chromosome 3 clearly distinguished six clusters of individuals along PC axes 1 and 2 (Fig. 1A). Four additional clusters were distinguished by PC axis 3 (Fig. 1B). We examined F_IS_ in each of the clusters to determine whether individuals were homozygous or heterozygous for the supergene. We examined the frequency of reference and alternative alleles relative to the *F. selysi* reference genome, which was constructed from a pool of M males (Brelsford et al. 2020), to make a preliminary determination about whether haplotypes were M-like or P-like. Overall, there were four distinct haplotypes in our focal populations: two were M (M_A_ and M_D_) and two were P (P_1_ and P_2_). We found all possible combinations in heterozygotes. The M_D_ was mostly found in a heterozygous state, with the exception of one newly-mated queen (M_D_M_D_) and seven males (M_D_) out of 239 total individuals carrying M_D_.

**Figure 1.**
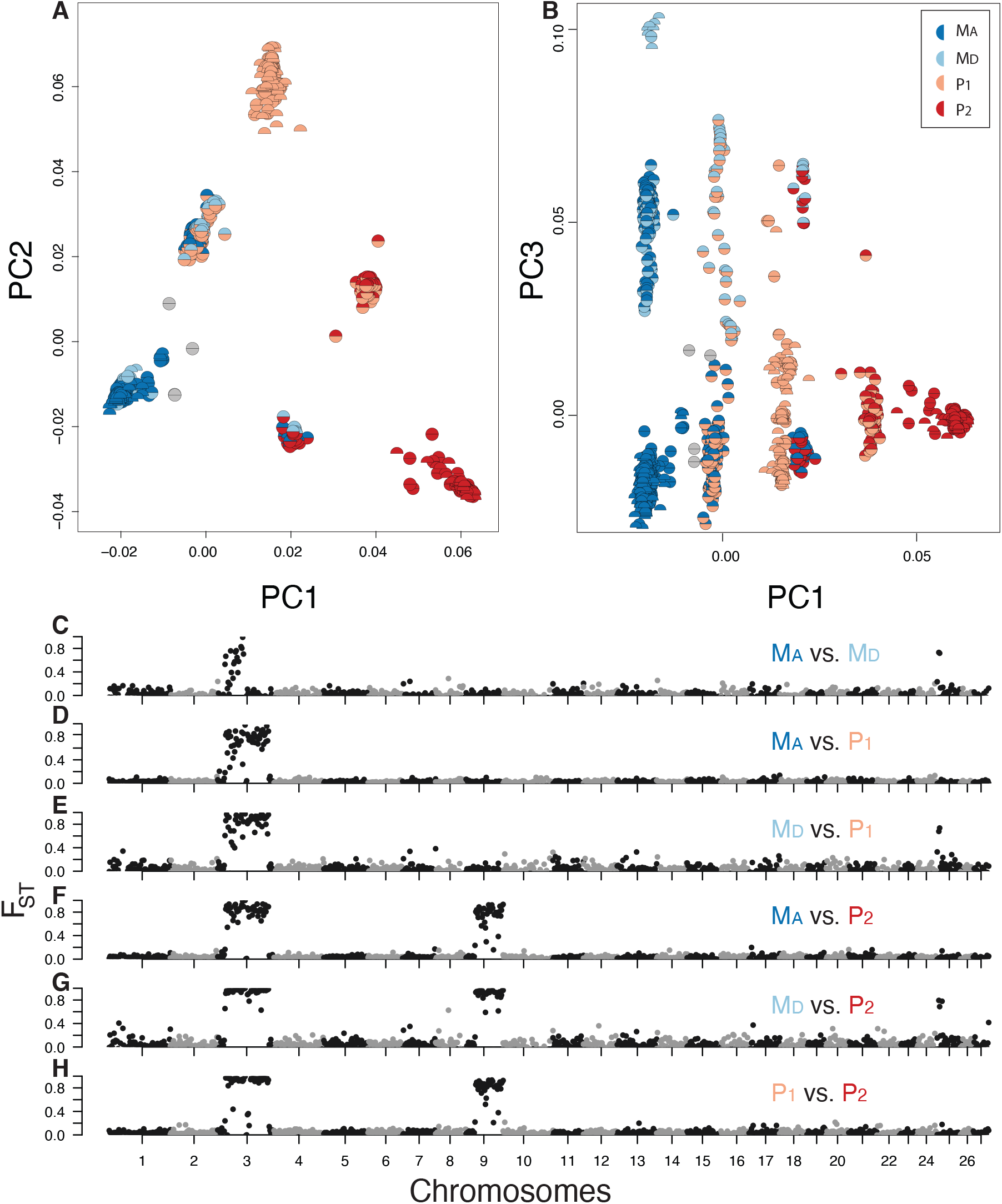
Principal component analysis and genetic differentiation identify four supergene haplotypes, including one encompassing variation on both chromosome 3 and chromosome 9. Principal component analysis axes 1 and 2 (**A**) distinguish six groups of individuals. The solid colored circles show homozygous individuals (according to positive F_IS_ values). Dual-color circles in clusters between homozygous groups show heterozygous individuals (according to negative F_IS_ values). Each half of the circle refers to a haplotype on chromosome 3, and haploid males are represented by half circles. Four additional clusters were distinguished by plotting PC3 (**B**), revealing the presence of a fourth haplotype (M_D_). With the exception of the M_D_ haplotype which spans only the first half of chromosome 3 (**C**), high differentiation (F_ST_) occurred between each haplotype across most of chromosome 3 when comparing haploid males (**D, E**). High differentiation is also evident on chromosome 9 (**F–H**), when comparing the P_2_ haplotype to the other three haplotypes. PC1 explains 51% of the total variance, while PC2 and PC3 explain respectively 29% and 3.8%.

To investigate genetic differences between alternative haplotypes, we looked at the F_ST_ between haplotype pairs, revealing that the M_D_ spans only the first half of chromosome 3 (Fig. 1C), while the genetic differentiation between M haplotypes and P_1_ spanned the same supergene region discovered in *F. selysi* (Figs. 1D-E; Purcell et al. 2014). The F_ST_ plots show high differentiation that spans almost all of chromosomes 3 and 9 when comparing P_2_ with the other haplotypes (Figs. 1F–H).

Given the second region of high differentiation between the P_2_ haplotype and all other supergene haplotypes, we investigated variation on chromosome 9. Here, we detected two alternative haplotypes. The PCA (Fig. 2A) displayed three distinct clusters of individuals along PC 1. Individuals in the left and right clusters appear to be homozygous based on positive F_IS_ values, while individuals in the central cluster are heterozygous. We named the two alternative haplotypes as follows: “9a” referring to the ancestral chromosome structure, and “9r” referring to the rearranged chromosome structure, as revealed by analysis of linkage disequilibrium within each homozygous genotype (Fig. S2A-B).

**Figure 2.**
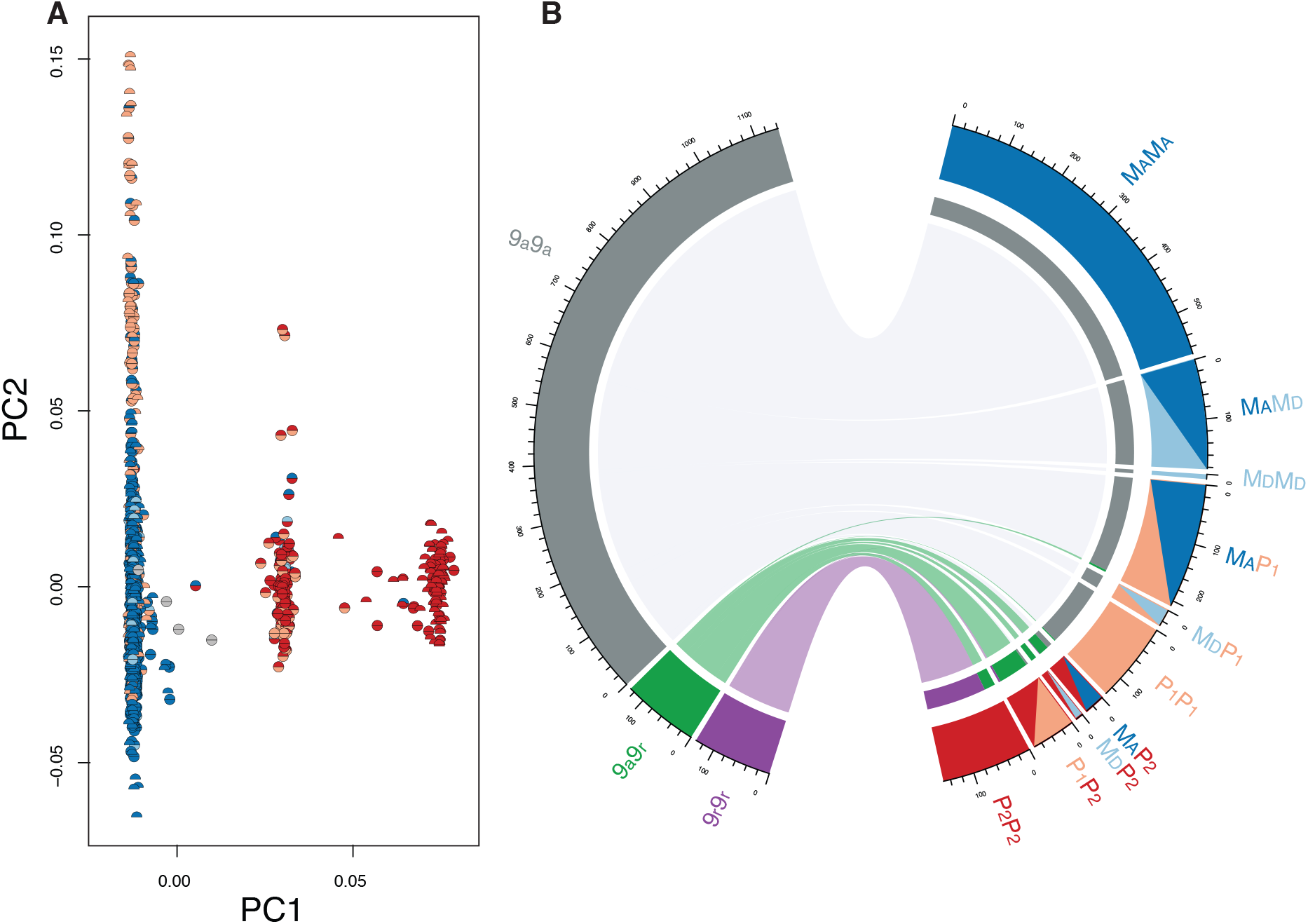
PCA of variants on chromosome 9 identified three clusters corresponding to three supergene genotypes, 9a9a (left cluster), 9a9r (middle cluster), and 9r9r (right cluster) (**A**). The colors of half circles in the PCA indicate chromosome 3 haplotypes to reveal mismatches between chromosomes 3 and 9. PC1 explains 75% of the total variance, and PC2 2.1%. Individuals with the 9r haplotype on chromosome 9 almost always have the P_2_ haplotype on chromosome 3, although we found some mismatches (14 out of 1151 9a9a individuals, all workers, harbor at least one copy of the P_2_; 7 out of 130 9a9r individuals, one gyne and six workers, do not carry the P_2_ haplotype). The chord diagram (**B**) shows associations between genotypes on chromosome 9 (left segments) and genotypes on chromosome 3 (right segments). Note that the lines connect chromosome 9 with chromosome 3 genotypes in the same individuals, and do not indicate regions of synteny.

The P_2_ haplotype on chromosome 3 and the 9r on chromosome 9 are almost always transmitted together. No P_2_P_2_ individual has been found to be 9a9a homozygous (Fig. S2C). In contrast, individuals without the P_2_ almost always bear only the 9a haplotype. However, we noticed some mismatches in this pattern showing an imperfect association between the P_2_ and 9r (Fig. 2B).

### Assessing colony social forms and their association with variants on chromosome 3

In other *Formica* species, colony social form is controlled by the social supergene on chromosome 3 (Brelsford et al. 2020). To verify that queen number is associated with variation on chromosome 3 in *F. cinerea*, we assessed colony social form and looked at supergene genotype distribution within colony. Of the 120 analyzed colonies, half were monogyne (39 monogyne monandrous, 21 monogyne polyandrous), and half polygyne (Fig. 3). We found a nearly perfect association of the M haplotypes with the monogyne form, where 59 out of 60 colonies contain exclusively M_A_M_A_ and M_D_M_A_ individuals (Fig. 3; Fig. S3A). Similarly, a strong association is observed between the P haplotypes and the polygyne form, with 56 out of 60 polygyne colonies having members with at least one P haplotype (Fig. 3; Fig. S3).

**Figure 3.**
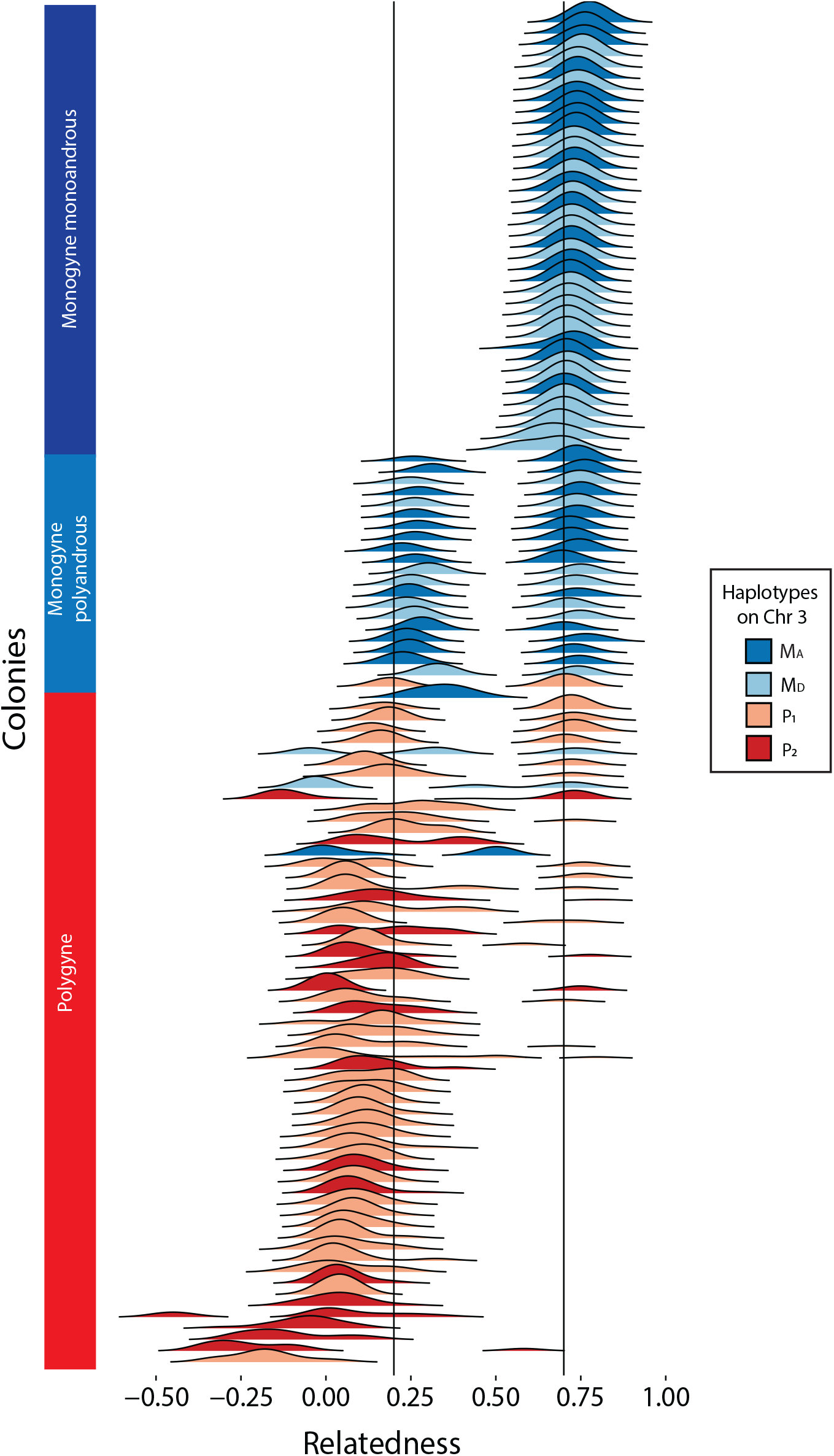
Ridgeline plots of the distribution of pairwise relatedness among nestmates reveal variation in colony social structure. Of the 120 colonies analyzed, half were monogyne (39 monandrous monogynous, 21 polyandrous monogynous) and half were polygyne. In 59 out of 60 monogyne colonies, we observed either exclusively M_A_M_A_ workers or a mix of M_A_M_A_ and M_A_M_D_ workers. In contrast, 56 out of 60 polygyne colonies harbored at least one copy of one of the P haplotypes (P_1_ and P_2_). The plot is colored to highlight the most frequent P haplotype within colony: P_2_>P_1_ (see also Suppl. fig. 2); in colonies lacking both P haplotypes, we colored each colony to highlight the presence of the M_D_ haplotype: M_D_>M_A_. Vertical lines at 0.7 and 0.2 show approximately where we expect peaks of full sibs and half-siblings respectively considering the downward bias typical of relatedness estimates based on RADseq markers (Attard et al., 2018).

Looking at supergene genotype distributions among colonies, we observe a hierarchical relationship between the haplotypes as follows:

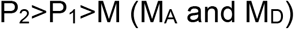

where P_2_, when present, is present in all individuals in at least one copy; in colonies with the P_1_, but not the P_2_, all the nestmates have at least one copy of the P_1_; in colonies lacking both P haplotypes, the M_A_ is always present while M_D_ is only present in a subset of colonies. Thirteen out of 120 colonies exhibited exceptions to this pattern (Fig. S3).

### Chromosome nine harbors a miniaturizing haplotype

Based on field observations that *F. cinerea* alates varied substantially in size, we measured the head width of gynes, queens and males. Our results reveal that alates with at least one copy of the P_2_ haplotype have significantly smaller heads than alates without the P_2_ (p < 0.0001) (Fig. 4A,C). However, this size reduction is caused by the 9r haplotype on chromosome 9 rather than the P_2_, as demonstrated by a GWAS analysis that identified numerous loci associated with alate size, all on chromosome 9 (Fig. 4E). The presence of a small gyne without the P_2_ but with the 9r is consistent with this pattern (Fig. 4A). On average, gynes with one copy of the 9r haplotype are 15.7% smaller than 9a9a gynes (p < 0.0001). This size reduction is 20.26% when gynes are 9r homozygous (p < 0.0001). 9r homozygous gynes are 5.42% smaller than 9r heterozygous gynes (p < 0.0001). Males exhibit a similar pattern, although this miniaturization appears to be less drastic, with 9r males being only 8.6% smaller than 9a males (p < 0.0001)(Fig. 4D).

**Figure 4.**
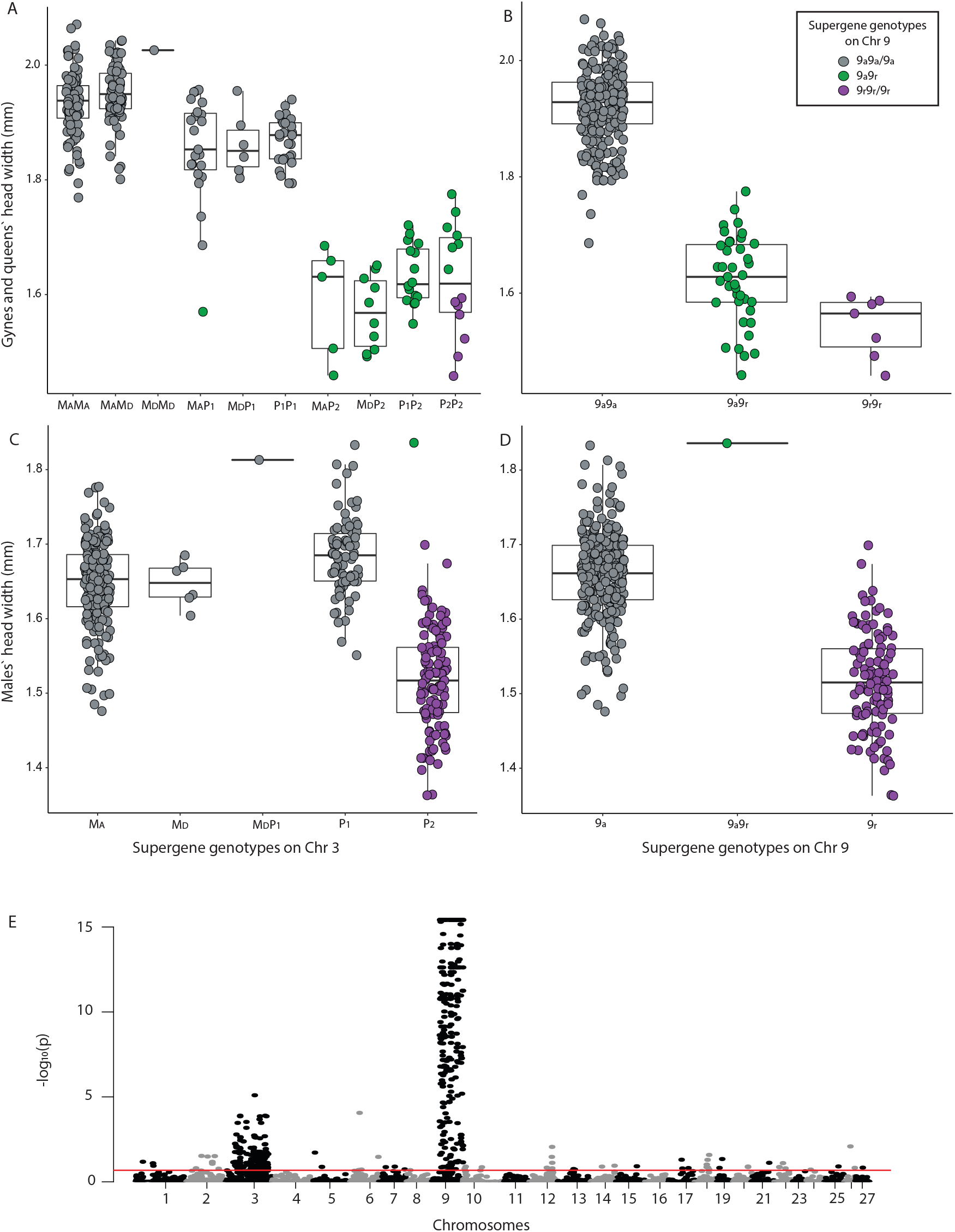
Head width (a good proxy for body size) is significantly reduced when at least one copy of the chromosome 3 P_2_ (gynes and queens: A, males: C) or chromosome 9 9r (gynes and queens: B, males: D) haplotype is present (p < 0.0001). GWAS analysis (**E**) confirms that a large region on chromosome 9 is most strongly associated with body size miniaturization in *F. cinerea*. The red line shows the significance threshold adjusted for false discovery rate.

### The M_D_ influences colony sex ratio

Some species of social insects during the reproductive period show a pattern of split sex ratio at the population level, in which some colonies specialize in the production of future queens and others in the production of males (Lagunas-Robles et al. 2021; Trivers and Hare 1976; Pamilo and Rosengren 1984; Boomsma and Grafen 1990). This also occurs in *F. cinerea*, especially in monogyne colonies (Fig. 5). In contrast, polygyne colonies more often produce a mix of males and gynes or exclusively males. Our data show that the M_D_ haplotype is associated with the production of females (gynes and workers) (X-squared = 344.71, df = 6, p-value < 0.0001). This haplotype is rarely present in males although we found some exceptions (7 M_D_ males).

**Figure 5.**
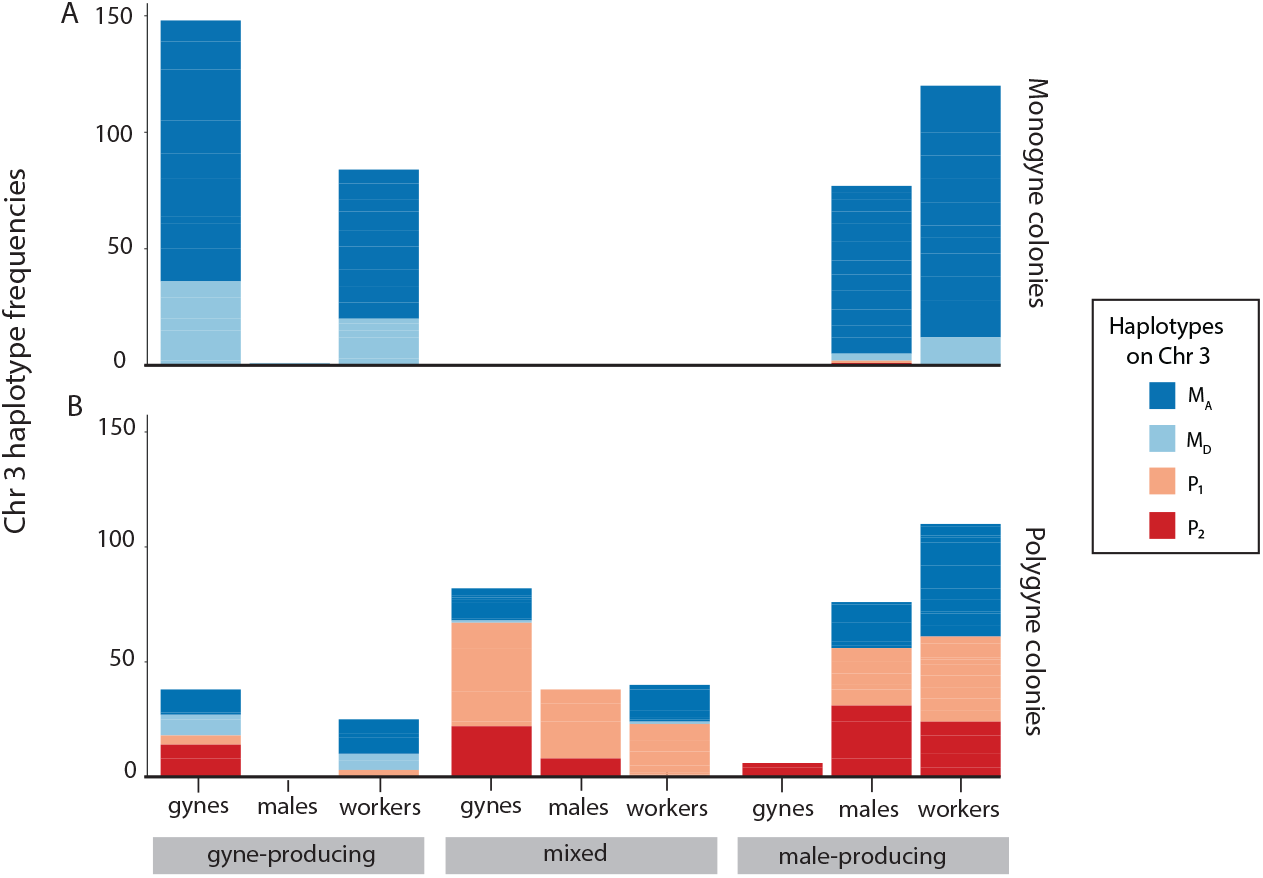
In monogyne colonies (A), the M_D_ haplotype is more common in gyne-producing than male-producing colonies and no mixed colonies were observed (X-squared = 344.71, df = 6, p-value < 0.0001). In polygyne colonies (B), the M_D_ haplotype was primarily found in the two gyne-producing colonies. In both social forms, male-producing colonies rarely bear the M_D_. In polygyne colonies, that P1 haplotype was also underrepresented in gyne-producing colonies.

## Discussion

Most ant species harboring a social supergene have only two alternative haplotypes, one associated with monogyny and the other associated with polygyny (Kay et al. 2022). Here we describe for the first time a species, *Formica cinerea*, that remarkably bears four supergene haplotypes on chromosome 3, all co-occurring in a single population. As found in several congeneric species so far (Purcell et al. 2014; Lagunas-Robles et al. 2021; McGuire et al. 2022; Pierce et al. 2022), the social form in *F. cinerea* is genetically controlled. Two M haplotypes (M_A_ and M_D_) are strongly associated with single-queen colonies, while two P haplotypes (P_1_ and P_2_) are almost exclusively present in multi-queen colonies. We discovered a novel rearranged supergene variant (9r) on chromosome 9 underlying queen miniaturization, highly linked to the P_2_ polygyne-associated haplotype.

### Linkage of socially antagonistic alleles and supergene expansion

Alternative social forms in ants generally conform to the “polygyny syndrome” in which gynes of polygyne colonies are about 10% smaller and have lower relative fat content than those produced by monogyne colonies (Keller 1993). In *F. cinerea* polygyne colonies, we observe two distinct gyne sizes: 9a9a gynes are relatively large (though still smaller on average than monogyne-produced gynes); in contrast, gynes with a 9r haplotype are 16-20% smaller than 9a9a gynes (Fig. 4A-B). This aligns with other cases of extreme queen-size dimorphism (microgynes and macrogynes) (Wolf and Seppä 2016). We observed a similar pattern in males (Fig. 4C-D). No 9r *F. cinerea* gynes or queens have been observed in monogyne colonies. Polygyny, therefore, appears to be a precondition for microgyny in this species.

We suggest that fitness epistasis initially emerged between an ancestral P haplotype and an incipient mutation on chromosome 9 that caused reduced body size in queens (Fig. 6C). In the process of establishing a new colony, macrogynes rely solely on their body reserves (wing muscles and fat bodies) to raise their first brood (Wheeler and Buck 1996; Peeters and Ito 2001). In order to be successful it is essential that they produce a worker caste in a short time, before depleting all their body reserves. The independent colony founding strategy is highly risky, and founding queens often suffer high mortality (Peeters and Ito 2001). Microgynes lack large fat reserves necessary to establish new nests (Keller 1993; Lachaud et al. 1999; Wolf and Seppä 2016), and thus, would be severely disadvantaged in an independent founding monogyne context. Conversely, in the polygyne background, colony foundation risks are reduced because queens can join preexisting colonies. Based on studies of microgynes in other species (Lachaud et al. 1999; Wolf and Seppä 2016), we hypothesize that *F. cinerea* microgynes are less costly to produce. Although they are expected to lay fewer eggs than macrogynes (Lachaud et al.1999; Wolf and Seppä 2016), their lower fecundity could be buffered by coexistence with other reproductive queens. Genetic mismatches between body size and colony social form would have a high cost, leading to strong selection for genetic linkage (Suppl. Fig. 1C). This expanded region of suppressed recombination spanning two chromosomes would include socially antagonistic alleles, beneficial in polygyne colonies but detrimental in monogyne colonies.

**Figure 6.**
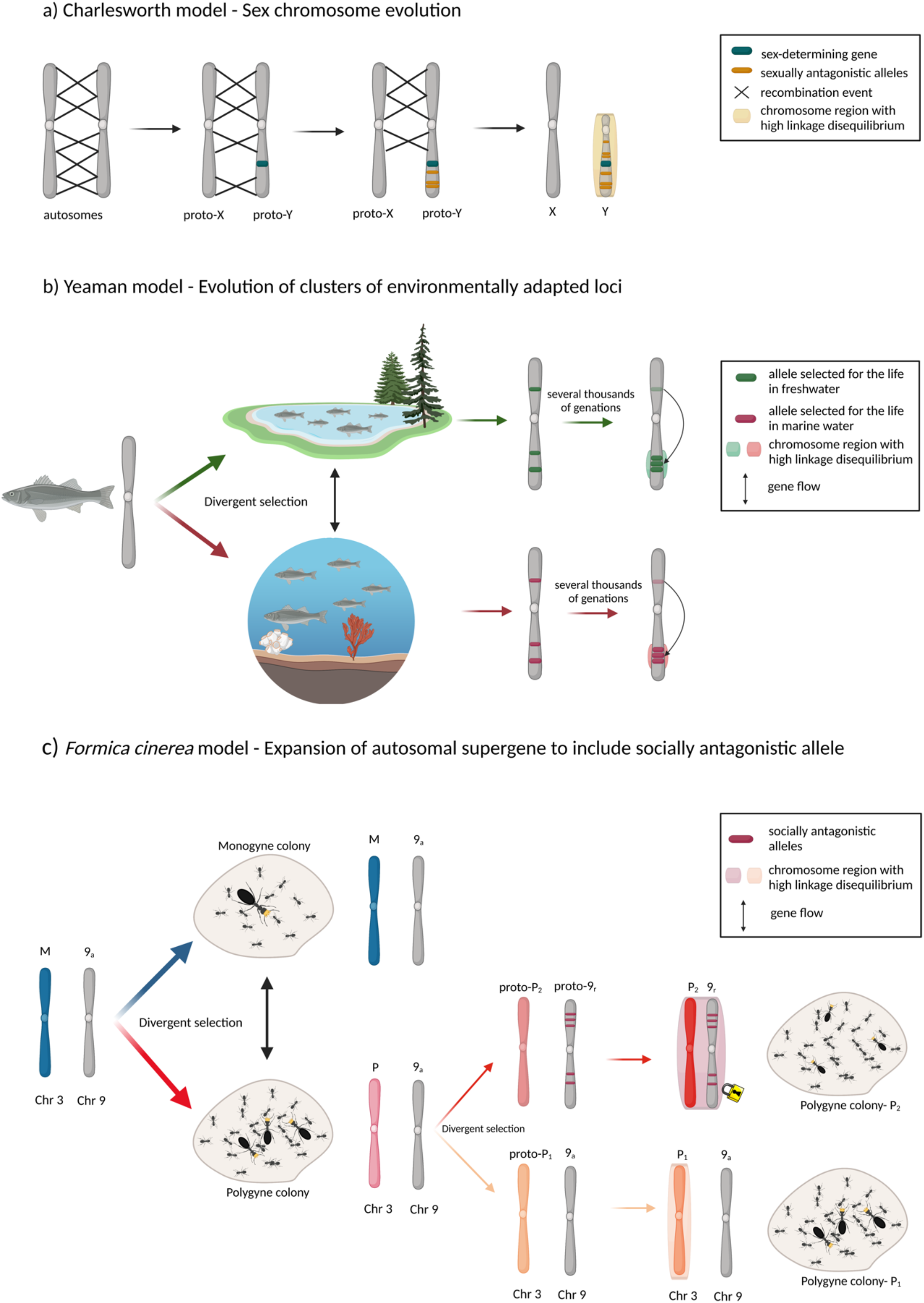
Overview of the dynamics of supergene evolution according to the Charlesworth model (A), the Yeaman model (B), and the case of *Formica cinerea* (C). The Charlesworth model (Charlesworth et al. 2005) suggested that in a hermaphrodite organism, a mutation on an autosomal chromosome caused the appearance of a sex-determining gene. Linked sexually antagonistic loci were then selected for reduced recombination (A). In the Yeaman (2013) model, there are loci under divergent selection between two environments connected by gene flow. Rearrangements that bring together clusters of these loci are favored by selection and allow weakly selected loci to diverge in opposition to gene flow. Populations inhabiting different environments diverge through the fixation of locally adapted alleles, and gene flow between the two populations selects for rearrangements that bring together clusters of loci involved in adaptation to the two environments (B). We hypothesize that in *Formica cinerea*, alternative social “environments” (experienced in monogyne and polygyne colonies, respectively) generate diverging selective pressures. If a miniaturizing allele arose on a chromosome unlinked to the social supergene, selection against that allele in monogyne colonies would oppose selection for it in polygyne colonies (social antagonism), and the allele would be unlikely to increase in frequency. A genomic rearrangement causing the miniaturizing allele to become linked to a chromosome that influences social form would itself be favored by selection, and would allow the miniaturizing allele to increase in frequency.

Ultimately, our results are consistent with predictions of the Charlesworth model in which antagonistic selection leads to the expansion of regions of suppressed recombination between advantageous combinations of alleles. Two novel features are present in our system. First, the expansion of LD is occurring in an autosomal supergene instead of in a sex chromosome (Charlesworth and Charlesworth 1978; Charlesworth et al. 2005). Second, the environment that shapes alternative traits is the social context determined by colony queen number as opposed to sex (Charlesworth & Charlesworth 1978) or the extrinsic environment (Yeaman 2013) (Fig. 6). We note that we cannot rule out the possibility that the initial mutation linking chromosome 9 and the P_2_ haplotype was selectively neutral, and that this linkage enabled the invasion of a queen miniaturizing mutation on chromosome 9 (reviewed by Ponnikas et al. 2018).

Although tightly linked, P_2_ and 9r are not perfectly co-transmitted (Fig. 2B). This occasional decoupling of linked alleles suggests that recombination occasionally happens between chromosomes 3 and 9, raising questions about how these two supergenes are linked. Several alternative mechanisms could mediate the incomplete linkage between P_2_ and 9r. We speculate that P_2_ and 9r may be linked by the fusion of chromosomes 3 and 9 or, alternatively, that they are linked through a reciprocal translocation. Further research is needed to identify the mechanism that locked these two regions of suppressed recombination together.

### Microgyny as an incipient form of intraspecific social parasitism?

Queen-size dimorphism associated with polygyny may lead to intraspecific parasitism, where the microgynes take advantage of the macrogynes by specializing in sexual offspring production (Schär and Nash 2014). In *Formica*, queen miniaturization was previously described only in species that parasitize other *Formica* species (*difficilis, dakotensis*, and *exsecta* clades) (Borowiec et al. 2021), although it has not been linked with colony social organization. Here we describe microgyny in a non-parasitic *Formica* species for the first time and speculate that 9r microgynes could be incipient intraspecific social parasites. The best-known case of intraspecific parasitism occurs in *Myrmica rubra: w*hen microgynes and macrogynes coexist in the same nest, microgynes produce very few worker offspring, focusing their reproductive effort mostly on sexuals (Schär and Nash 2014; Leppänen et al. 2015). During our field collections, we tried to minimize damage to nests, so we did not observe mature microgynes and macrogynes occurring together in the same nest. However, we found four colonies where all the workers were 9a9a homozygotes while alates were 9a9r and 9r9r microgynes. We also noticed that virgin microgynes and macrogynes were never produced by the same colony. Although preliminary, these findings could represent the first hint that *F. cinerea* microgynes are intraspecific social parasites.

### Supergene variation associated with three complex traits

Our results reveal that four haplotypes on chromosome 3 detected in *F. cinerea* are associated with at least three complex traits: social structure, alate size, and sex ratio. For the first time, we show evidence that microgyny is genetically controlled, as described above. As already studied in other *Formica* species, we confirmed that M haplotypes are associated with monogyne colonies, while P haplotypes are associated with polygyny. However, we found a few exceptions to this pattern: several apparently monogyne colonies include individuals with a P haplotype, and several apparently polygyne colonies lack P haplotypes (Figure S2).

A third complex phenotypic trait, colony sex ratio, is associated with the M_D_ haplotype, aligning with recent discoveries in *F. glacialis* and *F. podzolica* (Lagunas-Robles et al. 2021). We found that *F. cinerea* monogyne colonies, regardless of the number of matings, can specialize in the production of gynes or males. We show that split sex ratio is mediated by M_D_ and M_A_ haplotypes. Queens heterozygous for M_D_ tend to produce gynes, while queens homozygous for M_A_ tend to produce males. In contrast, polygyne colonies are mostly male-producing or mixed. Structurally, the M_D_ haplotype in *F. cinerea* spans the first half of chromosome 3 as in *F. glacialis* and *F. podzolica*. In a further parallel, we mainly found the M_D_ haplotype in heterozygous females, and observed a very low frequency of M_D_ homozygotes and haploids. We do not yet have enough information to determine whether these MD haplotypes share a common origin or originated independently.

## Conclusions

Unlike other ant species harboring a social supergene with only two alternative haplotypes (Kay et al. 2022; Chapuisat 2023), we found that colony queen number in *F. cinerea* is genetically controlled by a four-haplotype supergene on chromosome 3. Single-queen colonies contain almost exclusively M haplotypes (M_A_ and M_D_), while P haplotypes (P_1_ and P_2_) are strongly associated with multi-queen colonies. A novel supergene variant (9r) on chromosome 9 underlying a 16-20% reduction of queen body size (microgyny) is tightly linked to the polygyne-associated P_2_ haplotype on chromosome 3. Microgynes are absent from *F. cinerea* monogyne colonies, consistent with previous hypotheses that polygyny is a precondition for microgyny (Peeters and Ito 2001; Wolf & Seppä 2016). Here we propose that socially antagonistic selection favored the suppression of recombination between P haplotypes and a miniaturizing allele on chromosome 9, consistent with the Charlesworth model (Charlesworth et al. 2005). This novel association between regions of two chromosomes is analogous to the formation of neo-sex chromosomes (Charlesworth and Charlesworth 1980), as proposed in sticklebacks by Kitano et al. (2009) among other examples. Sex chromosome-autosome fusion may facilitate the process of local adaptation via geographic variation in the strength of sex-specific selection (Matsumoto et al. 2017). The social environment shaped by social supergenes is intermediate between the intrinsic physiological environment shaped by sex chromosomes and the extrinsic environment considered by the Yeaman model (Yeaman 2013). In other words, colony social organization is shaped by intrinsic characteristics of colony members and, in turn, it shapes the environmental conditions experienced by colony members. Thus, we propose that combining insights from models of sex chromosome evolution (Charlesworth et al. 2005) and local adaptation (Yeaman 2013) is an appropriate way to bridge the roles of intrinsic and extrinsic selection on recombination and genome structure.

## Material and Methods

*Formica cinerea* is a socially polymorphic species with a wide distribution across Europe (Seifert 2018). This species nests preferentially along sand and gravel banks of rivers and open sand dunes. We collected *F. cinerea* workers and alates (gynes and males) from colonies in Northern Italy (Aosta Valley and Piedmont) in June-July across several years, 2014, 2018-2021 (Table S1). Whenever possible, we sampled up to 10 gynes and males, and about 15 workers from each colony, and noted the observed sex-ratio. When multiple mature queens were found within colonies, we also sampled a subset of them. During 2019-2021, we collected newly mated wingless queens that were either looking for suitable locations to start new colonies or were under stones in self-dug chambers with none to few eggs. We stored samples in 96-100% ethanol.

### Library preparation

We extracted DNA from the head and thorax of workers, and only the head of gynes and males. For the 2014 and 2018-2020 samples, we used the QIAGEN DNeasy Blood & Tissue Kit with modifications described in McGuire et al. (2022). Briefly, we manually ground the tissue with sterile pestles in a 1.7 ml tube while immersed in liquid nitrogen, and left the pulverized samples overnight in buffer ATL and proteinase K at 56°C. We then used alternatively sourced spin columns (BPI-tech.com), 70% ethanol for DNA wash, and eluted the DNA in 30 µL of buffer EB. We extracted individuals collected in 2021 using the QiaAmp 96 DNA QiaCube HT kit and protocol. We manually ground the ant tissues as described above, and, following the overnight digestion in buffer ATL and proteinase K, we transferred the supernatant to the QIAcube HT/QIAxtractor robot to complete the extraction. We eluted the DNA in 100 µL of buffer EB.

We sequenced all samples using a double-digest restriction site-associated DNA sequencing (RADseq) approach (protocol from Brelsford et al. 2016). We digested the DNA using restriction enzymes MseI and PstI and ligated a universal MseI adapter and uniquely barcoded PstI adapter to each sample. We then removed small DNA fragments using Serapure magnetic beads (Rohland & Reich 2012) or Omega magnetic beads (Omega Biotek, 2021) in a 0.8:1 ratio (beads: sample solution). We amplified each sample in four separate PCR reactions with indexed Illumina primers and then pooled the replicate PCR products for each sample for a final PCR cycle. Finally, we pooled all PCR products in a tube and did a final round of small fragment removal using the magnetic beads. Sample sizes in each batch are provided (Table S2).

### Bioinformatics

We used *Stacks* 2.60 to demultiplex our data with default parameters (Catchen et al. 2013), *PEAR* v0.9.10 (Zhang et al. 2014) to merge paired-end reads and remove adaptor sequences, and *BWA-mem2* (Vasimuddin et al. 2019) to align reads to the *Formica selysi* genome (Brelsford et al. 2020). We called SNPs using *BCFtools mpileup* (Li and Durbin 2009) and filtered the genotypes for a minimum read depth of 7, a minor allele frequency of 5%, and excluded indels and sites with over 80% missing data using *VCFtools* 0.1.16-18 (Danecek et al. 2011).

#### Excluding duplicated regions

Ant males are haploid, and this feature provides an opportunity to identify and omit duplicated genomic regions. Males are treated as diploid in our initial pipeline, and loci that are heterozygous in at least 5% of males are flagged for removal from the complete dataset, because these reflect variable sequences in duplicated regions instead of alternative alleles in a single region of the genome.

#### Mitigating the batch effect

In order to have an adequate sample size for all supergene genotypes in all castes (particularly gynes and males, which are sampled opportunistically), we added data incrementally across years. Differences in extraction protocols and variation among sequencing lanes caused a batch effect (Fig. S1). To mitigate this issue, we calculated the Weir and Cockerham’s F_ST_ between batch pairs at each locus. We then removed all SNPs showing F_ST_ values ≥ 0.3 in the comparison of at least one pair of batches (because the geographic scope of sampling was similar across years, we would not expect to find true changes in allele frequency of this magnitude). Our final dataset resulted in 15129 SNPs and 1415 individuals (Table S3). Workers, gynes, males and mature queens were collected from 172 colonies, and 95 newly mated queens were collected from the environment.

### Determination of colony social form

We used COANCESTRY 1.0.1.10 (Wang 2011) to determine pairwise relatedness using workers and gynes (using Wang [2002] estimator) and infer colony social form. To ensure that these analyses were independent of our assessments of supergene variation, we created a dataset that excluded chromosome 3 and chromosome 9. To have a robust assignment, we kept only colonies with at least 5 diploid individuals and excluded haploid males. The final dataset resulted in 761 individuals and 120 colonies. A recent literature review and simulation study confirmed that relatedness estimates tend to be downward biased, yet more precise, in SNP-based datasets with hundreds or thousands of loci compared to microsatellite-based datasets with fewer loci (Attard et al. 2018). Given the known biases in datasets like ours, we called colonies with all pairwise relatedness estimates ≥ 0.6 as monogyne monandrous, colonies with bimodal distribution of pairwise relationships with at least 40% ≥ 0.6, but none <0.2 as monogyne polyandrous, and colonies with at least one pairwise relationship ≤ 0.1 as polygyne. We visualized the distribution of within-colony relatedness estimates with a ridgeline plot produced in R (R Core Team 2016) using the function *ggplot* (package ggplot2, Wickham 2009).

To investigate the association of the colony social organization with the supergene, we first performed a principal component analysis (PCA) for all individuals (workers, males, gynes and queens) using only the 1235 SNPs on chromosome 3, which contains the known *Formica* social supergene (Brelsford et al. 2020; Purcell et al. 2021). We then assigned the genotypes to each individual based on clusters in PCA and F_IS_ value (heterozygous individuals have negative F_IS_ values across the supergene, while homozygotes have positive values). To further investigate the genetic differentiation between each haplotype, we selected haploid males and calculated Weir and Cockerham’s F_ST_ for all pairwise combinations of supergene haplotypes. The PCA was calculated in PLINK (Purcell et al. 2007) with the --pca flag, while the F statistics were calculated in VCFtools (Danecek et al. 2011), using the --het flag (F_IS_) and the --weir-fst-pop flag. From the F_ST_ plot we noticed a second supergene on chromosome 9 visible when comparing the P_2_ haplotype (see Results section) on chromosome 3 with the other haplotypes. For chromosome 9, we performed a PCA and analysis of F_IS_ using only the 983 loci on that chromosome to assign genotypes to each individual. To identify which of the variants is rearranged relative to the *F. selysi* reference genome, we built two within-haplotype heatmaps of linkage disequilibrium using only homozygous individuals at each haplotype on chromosome 9. For this analysis, we used the LDheatmap package in R (Shin et al. 2016).

### Morphometrics

To assess whether polygyne *Formica cinerea* alates (gynes, queens and males) exhibit the reduction in size typical of polygyny syndrome (Keller 1993), we measured the maximum width across the eyes in 177 gynes and queens and 374 males using a Leica DMC2900 camera mounted on a Leica S8APO at 25× magnification. We used head width because it is known to have a strong positive correlation with several body segment dimensions in *Formica* species (Schwander et al. 2005; Tawdros et al. 2020), and thus serves as a good proxy for body size within caste. We fit two independent linear mixed models for chromosome 3 and chromosome 9 using colony as a random effect and genotype as a fixed effect to test whether gynes and queens with different supergene genotypes have significantly different sizes. We repeated the same analyses for males. For this analysis, we used the R package lme4 (Bates et al. 2014). Pairwise p-values were obtained after performing Tukey post hoc tests using the *emmeans* function in R (Lenth et al. 2019).

To identify genomic regions associated with body size, we performed a Genome Wide Association Study (GWAS) using a univariate linear mixed model implemented in *Gemma* v0.94 (Zhou and Stephens 2012). Males were excluded from this analysis. Since *Gemma* requires that no missing genotypes are present in the data, we imputed missing genotypes with *Beagle* v4.1 (Browning and Browning 2016) using the full dataset of SNPs that passed previously mentioned filters. *Gemma* uses a relatedness matrix generated from the sample genetic data to correct for non-independence of the samples due to population structure. We applied a false discovery rate adjustment (Benjamini and Hochberg 1995) to the *p*-values obtained from *Gemma* using the R function *p*.*adjust* (package stats, R Core Team et al. 2018).

### Sex ratio

While inspecting *F. cinerea* colonies during sampling, we took note of whether they exhibited a strongly skewed sex ratio, i.e. whether the colony preferentially produced gynes or males, or both sexes. We attributed the sex ratio to colonies observed with at least seven alates. Gyne producing colonies had at least seven gynes and no more than two males, male producing colonies had at least seven males and no more than two gynes, and mixed colonies were intermediate between the two. In total, 23 *F. cinerea* colonies were male-producing, 13 gyne-producing, and 6 were mixed. For each of these colonies, we looked at the haplotype frequencies on chromosome 3. We tested the significance of the association by performing a chi-square test on the haplotype frequencies.

## Data Availability

Genetic sequences and head width measurements of the alates reported herein will be deposited respectively at the NCBI Sequence Read Archive and Dryad Data Repository after the official acceptance of the paper.

## Acknowledgments

This material is based upon work supported by the US National Science Foundation DEB Grant No. 1733437 to A.B. and J.P. and DEB Grant No. 1631776 to J.P. We thank D. Zarate, G. Lagunas-Robles, Z. Alam, D. Pierce, N. Najar, and the FoG discussion group (Campbell, Samuk, and Ostevik labs) for providing helpful feedback on an earlier version of the manuscript and M. Molfini for assistance in the field.

