## Supplemental figures and tables for "Does social antagonism facilitate supergene expansion? A novel region of suppressed recombination in a 4-haplotype supergene system"

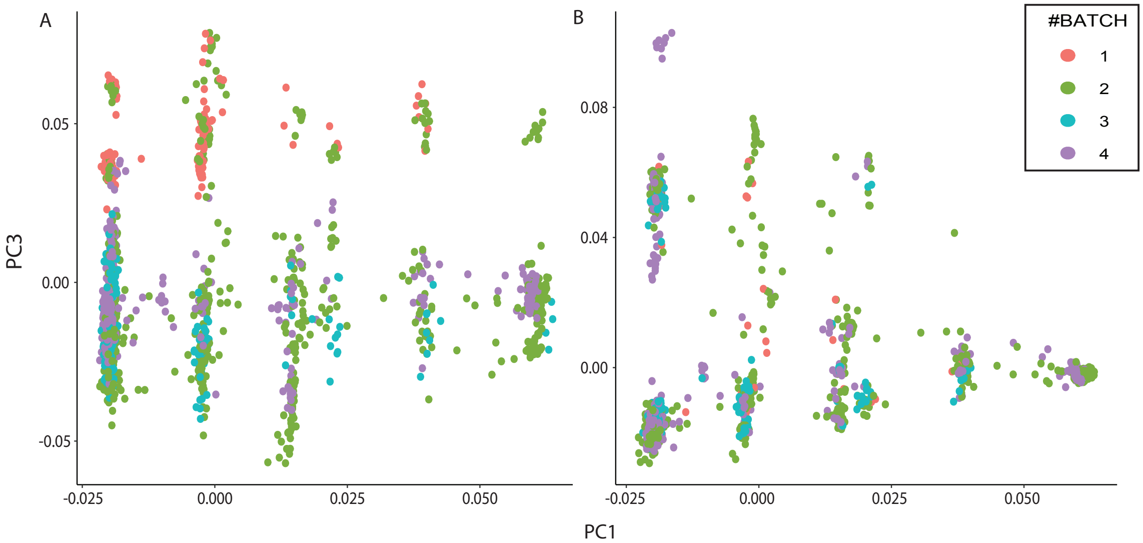


**Suppl. figure 1.** The PCA shows the presence of a batch effect caused by variation among sequencing lanes across the years, where samples group according to the batch origin rather than following a biological pattern (A). The batch effect was only visible when plotting PC1 versus PC3 thus hiding the presence of the $M_D$ haplotype, whereas there is no apparent problem when plotting PC1 versus PC2. Removing all SNPs showing $F_{ST}$ values ≥ 0.3 in comparison to at least one pair of batches, mitigated the batch effect problem (B).

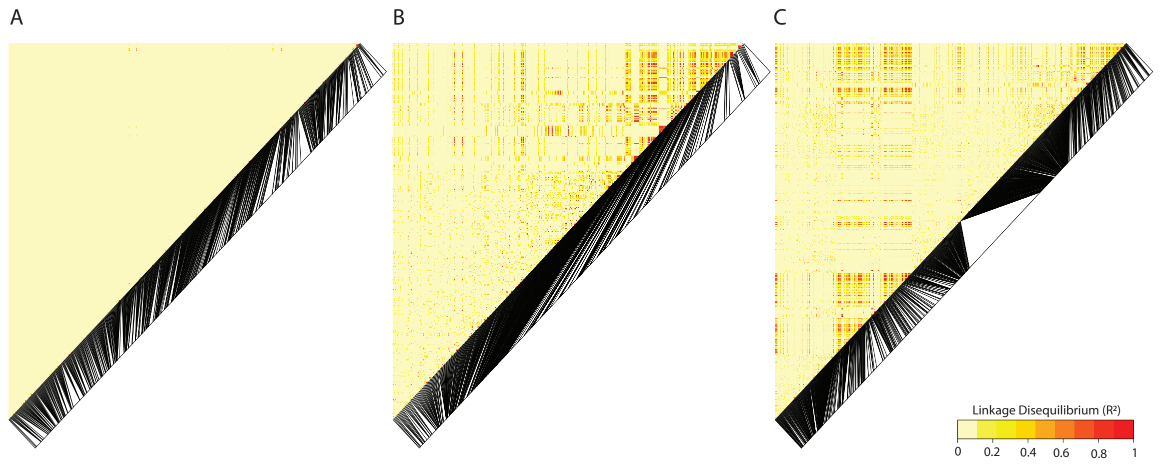


**Suppl. figure 2**. Heatmaps display pairwise linkage disequilibria between loci within 9a (A), within 9r (B), and between 9r on chromosome 9 and $P_2$ on chromosome 3 (C). No evidence of suppressed recombination is visible in 9a (A), while several markers in 9r haplotypes show high genetic linkage (B) confirming 9r as the recombinant variant. Among chromosomes, 9r and $P_2$ show many loci at high linkage disequilibrium.

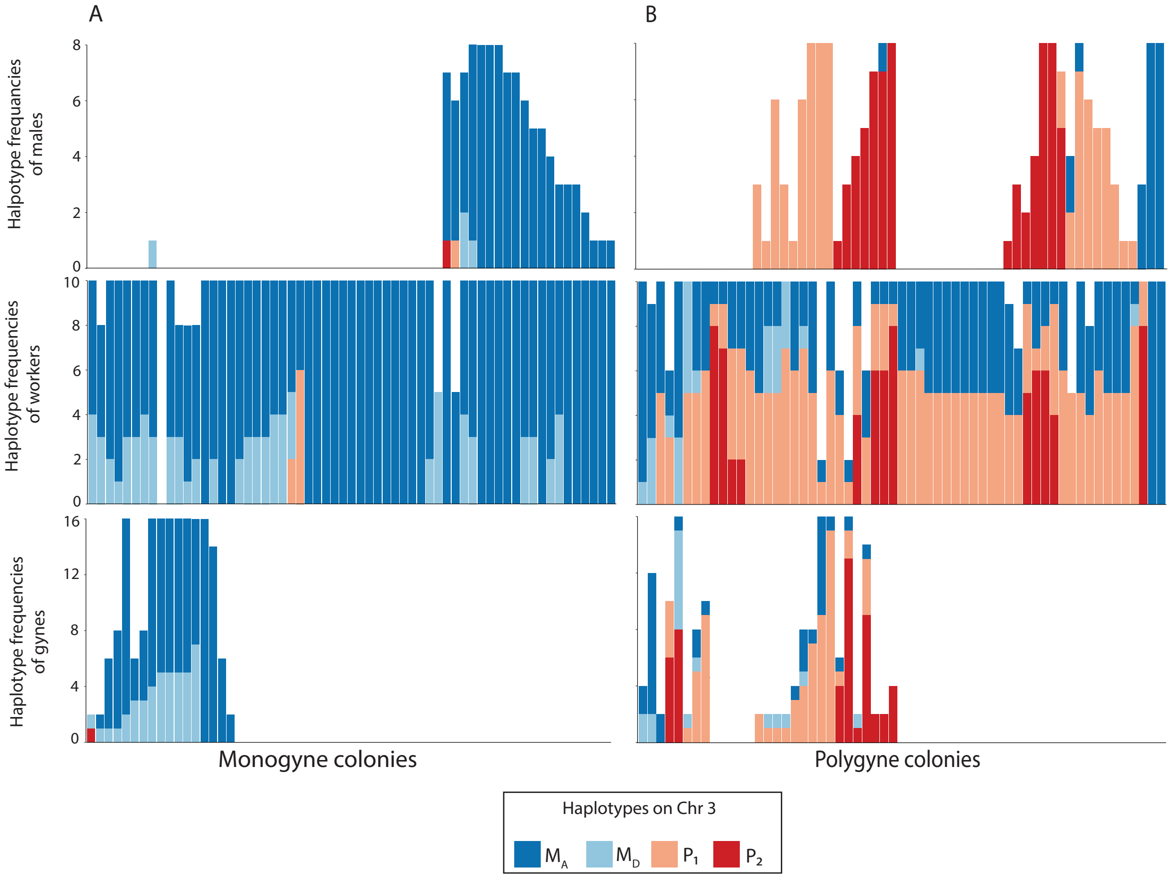


**Suppl. figure 3**. Stacked bar plots display within-colony haplotype frequencies on chromosome 3 in males (on top), workers (in the middle), and gynes (on the bottom) in monogyne (A) and polygyne (B) colonies. Individuals from monogyne colonies carry almost exclusively $M_A$ and $M_D$ haplotypes, although occasionally we detected some individuals bearing the $P_1$ or $P_2$. Conversely, individuals from polygyne colonies carry at least one P haplotype ($P_1$ or $P_2$), with occasional exceptions. Within colonies, there is an overrepresentation of one haplotype, present in at least one copy in each individual, and males usually harbor the same haplotype. Taken together these observations may indicate the presence of a transmission distortion mechanism.

**Table S1:** Sampling Locality information

| Locality Code | Locality Name | Latitude | Longitude | Elevation (m) |
|---|---|---|---|---|
| ANO | Anzola d'Ossola | 45.992697 | 8.3516450 | 209 |
| ARN | Arnard | 45.636286 | 7.7173270 | 358 |
| DOM | Domodossola | 46.122783 | 8.2763440 | 321 |
| DON | Donnas | 45.599903 | 7.7726580 | 322 |
| MAS | Masera | 46.133152 | 8.3208560 | 297 |
| MJO | Montjovet | 45.705561 | 7.6733610 | 406 |
| QUA | Quassolo | 45.537809 | 7.8341610 | 268 |
| QUIN | Quincinetto | 45.564840 | 7.8149820 | 291 |
| QUINI | Quincinetto isola | 45.575720 | 7.8093360 | 289 |
| RUM | Rumanica | 45.999188 | 8.2945050 | 217 |
| SV | Settimo Vittone | 45.548934 | 7.8250290 | 274 |
| VAR | Varzo | 46.206321 | 8.2377130 | 532 |

**Table S2.** Overview of batches

| # Batch | # *F. cinerea* samples | Sequencing facility | Sequencer |
|---|---|---|---|
| 1 | 138 | Novogene | HiSeq X Ten |
| 2 | 645 | UCSD Institute for Genomic Medicine | Novaseq 6000 |
| 3 | 167 | UCSD Institute for Genomic Medicine | Novaseq 6000 |
| 4 | 465 | UCSD Institute for Genomic Medicine | Novaseq 6000 |

**Table S3:** Overview of samples

| SAMPLE ID | CASTE | COLONY | YEAR | BATCH | GENOTYPES CHR3 | GENOTYPES CHR9 |
|---|---|---|---|---|---|---|
| 2020-MQ1 | MQ | SV2002 | 2020 | 4 | MAP1 | 9a9a |
| 2020-MQ2 | MQ | SV2003 | 2020 | 4 | P1P1 | 9a9a |
| 2020-MQ3 | MQ | QUINI20 | 2020 | 4 | MAP1 | 9a9a |
| 2020-NQ1 | NQ | - | 2020 | 4 | MAMA | 9a9a |
| 2020-NQ2 | NQ | - | 2020 | 4 | MAMA | 9a9a |
| 2020-NQ3 | NQ | - | 2020 | 4 | MAMA | 9a9a |
| 2020-NQ4 | NQ | - | 2020 | 4 | MAMD | 9a9a |
| 2020-NQ5 | NQ | - | 2020 | 4 | MAMA | 9a9a |
| 2020-NQ6 | NQ | - | 2020 | 4 | MAMD | 9a9a |
| 2021-MQ1 | MQ | ARN21-02 | 2021 | 4 | P1P1 | 9a9a |
| 2021-MQ2 | MQ | ARN21-02 | 2021 | 4 | P1P2 | 9a9r |
| 2021-MQ3 | MQ | QUINI21-02 | 2021 | 4 | P1P2 | 9a9r |
| 2021-MQ4 | MQ | QUINI21-02 | 2021 | 4 | P1P2 | 9a9r |
| 2021-MQ5 | MQ | ARN21-04 | 2021 | 4 | MAP2 | 9a9r |
| 2021-MQ6 | MQ | - | 2021 | 4 | MAP1 | 9a9a |
| 2021-NQ1 | NQ | - | 2021 | 4 | MAMA | 9a9a |
| 2021-NQ10 | NQ | - | 2021 | 4 | MAMD | 9a9a |
| 2021-NQ11 | NQ | - | 2021 | 4 | MAMD | 9a9a |
| 2021-NQ12 | NQ | - | 2021 | 4 | MAMD | 9a9a |
| 2021-NQ13 | NQ | - | 2021 | 4 | MAMD | 9a9a |
| 2021-NQ14 | NQ | - | 2021 | 4 | MAMA | 9a9a |
| 2021-NQ15 | NQ | - | 2021 | 4 | MAMD | 9a9a |
| 2021-NQ16 | NQ | - | 2021 | 4 | MAMD | 9a9a |
| 2021-NQ17 | NQ | - | 2021 | 4 | MAMA | 9a9a |
| 2021-NQ18 | NQ | - | 2021 | 4 | MAMA | 9a9a |
| 2021-NQ19 | NQ | - | 2021 | 4 | MDMD | 9a9a |
| 2021-NQ2 | NQ | - | 2021 | 4 | MAMD | 9a9a |
| 2021-NQ20 | NQ | - | 2021 | 4 | MAMA | 9a9a |
| 2021-NQ21 | NQ | - | 2021 | 4 | MAMD | 9a9a |
| 2021-NQ22 | NQ | - | 2021 | 4 | MAMA | 9a9a |
| 2021-NQ23 | NQ | - | 2021 | 4 | MAMD | 9a9a |
| 2021-NQ24 | NQ | - | 2021 | 4 | MAMD | 9a9a |
| 2021-NQ25 | NQ | - | 2021 | 4 | MAMA | 9a9a |
| 2021-NQ26 | NQ | - | 2021 | 4 | MAMA | 9a9a |
| 2021-NQ27 | NQ | - | 2021 | 4 | MAMA | 9a9a |
| 2021-NQ28 | NQ | - | 2021 | 4 | MAMA | 9a9a |
| 2021-NQ29 | NQ | - | 2021 | 4 | P1P2 | 9a9r |
| 2021-NQ3 | NQ | - | 2021 | 4 | MAMD | 9a9a |

| | | | | | | |
|---|---|---|---|---|---|---|
| 2021-NQ30 | NQ | - | 2021 | 4 | MAMA | 9a9a |
| 2021-NQ31 | NQ | - | 2021 | 4 | MAMD | 9a9a |
| 2021-NQ32 | NQ | - | 2021 | 4 | MAMD | 9a9a |
| 2021-NQ33 | NQ | - | 2021 | 4 | MAMA | 9a9a |
| 2021-NQ34 | NQ | - | 2021 | 4 | MAMA | 9a9a |
| 2021-NQ35 | NQ | - | 2021 | 4 | MAMA | 9a9a |
| 2021-NQ36 | NQ | - | 2021 | 4 | MAMD | 9a9a |
| 2021-NQ37 | NQ | - | 2021 | 4 | MAMD | 9a9a |
| 2021-NQ38 | NQ | - | 2021 | 4 | MAMA | 9a9a |
| 2021-NQ39 | NQ | - | 2021 | 4 | MAMA | 9a9a |
| 2021-NQ4 | NQ | - | 2021 | 4 | MAMD | 9a9a |
| 2021-NQ40 | NQ | - | 2021 | 4 | MAMD | 9a9a |
| 2021-NQ41 | NQ | - | 2021 | 4 | MAMD | 9a9a |
| 2021-NQ42 | NQ | - | 2021 | 4 | MAMD | 9a9a |
| 2021-NQ43 | NQ | - | 2021 | 4 | MAMA | 9a9a |
| 2021-NQ44 | NQ | - | 2021 | 4 | MAMD | 9a9a |
| 2021-NQ45 | NQ | - | 2021 | 4 | MAMD | 9a9a |
| 2021-NQ46 | NQ | - | 2021 | 4 | MAMA | 9a9a |
| 2021-NQ47 | NQ | - | 2021 | 4 | P2P2 | 9r9r |
| 2021-NQ48 | NQ | - | 2021 | 4 | MAMA | 9a9a |
| 2021-NQ49 | NQ | - | 2021 | 4 | MAMD | 9a9a |
| 2021-NQ5 | NQ | - | 2021 | 4 | MAMA | 9a9a |
| 2021-NQ50 | NQ | - | 2021 | 4 | MAMA | 9a9a |
| 2021-NQ51 | NQ | - | 2021 | 4 | MAMA | 9a9a |
| 2021-NQ52 | NQ | - | 2021 | 4 | MAMA | 9a9a |
| 2021-NQ53 | NQ | - | 2021 | 4 | MAMA | 9a9a |
| 2021-NQ54 | NQ | - | 2021 | 4 | MAMA | 9a9a |
| 2021-NQ55 | NQ | - | 2021 | 4 | MAMD | 9a9a |
| 2021-NQ56 | NQ | - | 2021 | 4 | MAMD | 9a9a |
| 2021-NQ57 | NQ | - | 2021 | 4 | MAMD | 9a9a |
| 2021-NQ58 | NQ | - | 2021 | 4 | MAMD | 9a9a |
| 2021-NQ59 | NQ | - | 2021 | 4 | MAMD | 9a9a |
| 2021-NQ6 | NQ | - | 2021 | 4 | MAMA | 9a9a |
| 2021-NQ60 | NQ | - | 2021 | 4 | MAMA | 9a9a |
| 2021-NQ61 | NQ | - | 2021 | 4 | MAMA | 9a9a |
| 2021-NQ62 | NQ | - | 2021 | 4 | MAMD | 9a9a |
| 2021-NQ63 | NQ | - | 2021 | 4 | MAMA | 9a9a |
| 2021-NQ64 | NQ | - | 2021 | 4 | MAMA | 9a9a |
| 2021-NQ65 | NQ | - | 2021 | 4 | P1P2 | 9a9r |
| 2021-NQ66 | NQ | - | 2021 | 4 | MAMA | 9a9a |
| 2021-NQ67 | NQ | - | 2021 | 4 | MAMA | 9a9a |
| 2021-NQ68 | NQ | - | 2021 | 4 | MAMA | 9a9a |
| 2021-NQ69 | NQ | - | 2021 | 4 | MAMA | 9a9a |
| 2021-NQ7 | NQ | - | 2021 | 4 | MAMD | 9a9a |

| | | | | | | |
|---|---|---|---|---|---|---|
| 2021-NQ70 | NQ | - | 2021 | 4 | MAMA | 9a9a |
| 2021-NQ71 | NQ | - | 2021 | 4 | MAMA | 9a9a |
| 2021-NQ73 | NQ | - | 2021 | 4 | MAMA | 9a9a |
| 2021-NQ74 | NQ | - | 2021 | 4 | MAMD | 9a9a |
| 2021-NQ75 | NQ | - | 2021 | 4 | MAMA | 9a9a |
| 2021-NQ76 | NQ | - | 2021 | 4 | MAMA | 9a9a |
| 2021-NQ77 | NQ | - | 2021 | 4 | MAMA | 9a9a |
| 2021-NQ78 | NQ | - | 2021 | 4 | MAMD | 9a9a |
| 2021-NQ79 | NQ | - | 2021 | 4 | MAMD | 9a9a |
| 2021-NQ8 | NQ | - | 2021 | 4 | MAMA | 9a9a |
| 2021-NQ80 | NQ | - | 2021 | 4 | MAMA | 9a9a |
| 2021-NQ81 | NQ | - | 2021 | 4 | MAMA | 9a9a |
| 2021-NQ82 | NQ | - | 2021 | 4 | MAP2 | 9a9r |
| 2021-NQ9 | NQ | - | 2021 | 4 | MAMD | 9a9a |
| ANO2101-M1 | M | ANO2101 | 2021 | 4 | MA | 9a9a |
| ANO2101-M2 | M | ANO2101 | 2021 | 4 | MA | 9a9a |
| ANO2101-M3 | M | ANO2101 | 2021 | 4 | MA | 9a9a |
| ANO2101-M7 | M | ANO2101 | 2021 | 4 | MA | 9a9a |
| ANO2101-M8 | M | ANO2101 | 2021 | 4 | MA | 9a9a |
| ANO2101-W1 | W | ANO2101 | 2021 | 3 | MAMA | 9a9a |
| ANO2101-W2 | W | ANO2101 | 2021 | 4 | MAMA | 9a9a |
| ANO2101-W3 | W | ANO2101 | 2021 | 4 | MAMA | 9a9a |
| ANO2101-W4 | W | ANO2101 | 2021 | 4 | MAMA | 9a9a |
| ANO2101-W5 | W | ANO2101 | 2021 | 4 | MAMA | 9a9a |
| ANO2102-M1 | M | ANO2102 | 2021 | 4 | MA | 9a9a |
| ANO2102-M2 | M | ANO2102 | 2021 | 4 | MA | 9a9a |
| ANO2102-M3 | M | ANO2102 | 2021 | 4 | MA | 9a9a |
| ANO2102-M4 | M | ANO2102 | 2021 | 4 | MA | 9a9a |
| ANO2102-W2 | W | ANO2102 | 2021 | 4 | MAMA | 9a9a |
| ANO2102-W3 | W | ANO2102 | 2021 | 4 | MAMA | 9a9a |
| ANO2102-W4 | W | ANO2102 | 2021 | 4 | MAMA | 9a9a |
| ANO2102-W5 | W | ANO2102 | 2021 | 4 | MAMA | 9a9a |
| ANO2103-W1 | W | ANO2103 | 2021 | 3 | MAMA | 9a9a |
| ANO2103-W2 | W | ANO2103 | 2021 | 4 | MAMA | 9a9a |
| ANO2103-W3 | W | ANO2103 | 2021 | 4 | MAMA | 9a9a |
| ANO2103-W4 | W | ANO2103 | 2021 | 4 | MAMA | 9a9a |
| ANO2103-W5 | W | ANO2103 | 2021 | 4 | MAMA | 9a9a |
| ANO2104-G1 | G | ANO2104 | 2021 | 4 | MDP2 | 9a9r |
| ANO2104-W1 | W | ANO2104 | 2021 | 3 | MAMA | 9a9a |
| ANO2104-W2 | W | ANO2104 | 2021 | 4 | MAMD | 9a9a |
| ANO2104-W3 | W | ANO2104 | 2021 | 4 | MAMD | 9a9a |
| ANO2104-W4 | W | ANO2104 | 2021 | 4 | MAMD | 9a9a |
| ANO2104-W5 | W | ANO2104 | 2021 | 4 | MAMD | 9a9a |
| ANO2105-W1 | W | ANO2105 | 2021 | 3 | MAMA | 9a9a |

| | | | | | | |
|---|---|---|---|---|---|---|
| ANO2105-W2 | W | ANO2105 | 2021 | 4 | MAMA | 9a9a |
| ANO2105-W3 | W | ANO2105 | 2021 | 4 | MAMD | 9a9a |
| ANO2105-W4 | W | ANO2105 | 2021 | 4 | MAMD | 9a9a |
| ANO2105-W5 | W | ANO2105 | 2021 | 4 | MAMD | 9a9a |
| ANO2106-W1 | W | ANO2106 | 2021 | 3 | MAMA | 9a9a |
| ANO2106-W2 | W | ANO2106 | 2021 | 4 | MAMA | 9a9a |
| ANO2106-W3 | W | ANO2106 | 2021 | 4 | MAMA | 9a9a |
| ANO2106-W4 | W | ANO2106 | 2021 | 4 | MAMA | 9a9a |
| ANO2106-W5 | W | ANO2106 | 2021 | 4 | MAMA | 9a9a |
| ANO2107-W1 | W | ANO2107 | 2021 | 3 | MAMD | 9a9a |
| ANO2107-W2 | W | ANO2107 | 2021 | 4 | MAMD | 9a9a |
| ANO2107-W3 | W | ANO2107 | 2021 | 4 | MAMA | 9a9a |
| ANO2107-W4 | W | ANO2107 | 2021 | 4 | MAMA | 9a9a |
| ANO2107-W5 | W | ANO2107 | 2021 | 4 | MAMD | 9a9a |
| ARN18C4-G2 | G | ARN18C4 | 2018 | 2 | MAMD | 9a9a |
| ARN18C4-G3 | G | ARN18C4 | 2018 | 2 | MAMA | 9a9a |
| ARN18C4-G4 | G | ARN18C4 | 2018 | 2 | MAMA | 9a9a |
| ARN18C4-G5 | G | ARN18C4 | 2018 | 2 | MAMA | 9a9a |
| ARN18C4-G6 | G | ARN18C4 | 2018 | 2 | MAMA | 9a9a |
| ARN18C4-G7 | G | ARN18C4 | 2018 | 2 | MAMD | 9a9a |
| ARN18C4-Q1 | Q | ARN18C4 | 2018 | 2 | MAMD | 9a9a |
| ARN18C4-Q2 | Q | ARN18C4 | 2018 | 2 | MAMD | 9a9a |
| ARN18C4-W1 | W | ARN18C4 | 2018 | 1 | MAMA | 9a9a |
| ARN18C4-W2 | W | ARN18C4 | 2018 | 1 | MAMD | 9a9a |
| ARN18C4-W3 | W | ARN18C4 | 2018 | 1 | MAMD | 9a9a |
| ARN18C4-W4 | W | ARN18C4 | 2018 | 1 | MAMD | 9a9a |
| ARN18C4-W5 | W | ARN18C4 | 2018 | 1 | MAMA | 9a9a |
| ARN18G4-W1 | W | ARN18G4 | 2018 | 2 | MAP1 | 9a9a |
| ARN18G4-W2 | W | ARN18G4 | 2018 | 2 | MAP1 | 9a9a |
| ARN18G4-W3 | W | ARN18G4 | 2018 | 2 | MAP1 | 9a9a |
| ARN18G4-W4 | W | ARN18G4 | 2018 | 2 | MAP1 | 9a9a |
| ARN18G4-W5 | W | ARN18G4 | 2018 | 2 | MAP1 | 9a9a |
| ARN19C1-G2 | G | ARN19C1 | 2019 | 2 | MAMA | 9a9a |
| ARN19C1-G3 | G | ARN19C1 | 2019 | 2 | MAMA | 9a9a |
| ARN19C1-G4 | G | ARN19C1 | 2019 | 2 | MAMD | 9a9a |
| ARN19C1-G5 | G | ARN19C1 | 2019 | 2 | MAMD | 9a9a |
| ARN19C1-G6 | G | ARN19C1 | 2019 | 2 | MAMD | 9a9a |
| ARN19C1-G7 | G | ARN19C1 | 2019 | 2 | MAMD | 9a9a |
| ARN19C1-G8 | G | ARN19C1 | 2019 | 2 | MAMA | 9a9a |
| ARN19C1-G9 | G | ARN19C1 | 2019 | 2 | MAMD | 9a9a |
| ARN19C2-G1 | G | ARN19C2 | 2019 | 2 | MAMD | 9a9a |
| ARN19C2-G2 | G | ARN19C2 | 2019 | 2 | MAP1 | 9a9a |
| ARN19C2-M2 | M | ARN19C2 | 2019 | 2 | MA | 9a9a |
| ARN19C2-M3 | M | ARN19C2 | 2019 | 2 | MA | 9a9a |

| | | | | | | |
|---|---|---|---|---|---|---|
| ARN19C3-W1 | W | ARN19C3 | 2019 | 2 | MAP1 | 9a9a |
| ARN19C3-W2 | W | ARN19C3 | 2019 | 2 | P1P1 | 9a9a |
| ARN19C3-W3 | W | ARN19C3 | 2019 | 2 | MAP1 | 9a9a |
| ARN19C3-W4 | W | ARN19C3 | 2019 | 2 | MAP1 | 9a9a |
| ARN19C3-W5 | W | ARN19C3 | 2019 | 2 | MDP1 | 9a9a |
| ARN19C4-M2 | M | ARN19C4 | 2019 | 2 | MA | 9a9a |
| ARN19G2-W1 | W | ARN19G2 | 2019 | 2 | MAMA | 9a9a |
| ARN19G2-W2 | W | ARN19G2 | 2019 | 2 | MAMA | 9a9a |
| ARN19G2-W3 | W | ARN19G2 | 2019 | 2 | MAMA | 9a9a |
| ARN19G2-W4 | W | ARN19G2 | 2019 | 2 | MAMA | 9a9a |
| ARN19G2-W5 | W | ARN19G2 | 2019 | 2 | MAMA | 9a9a |
| ARN19G3-G1 | G | ARN19G3 | 2019 | 2 | MAMD | 9a9a |
| ARN19G3-G2 | G | ARN19G3 | 2019 | 2 | MAMD | 9a9a |
| ARN19G3-W1 | W | ARN19G3 | 2019 | 2 | MAMD | 9a9a |
| ARN19G3-W2 | W | ARN19G3 | 2019 | 2 | MAMA | 9a9a |
| ARN19G3-W3 | W | ARN19G3 | 2019 | 2 | MAMA | 9a9a |
| ARN19G3-W4 | W | ARN19G3 | 2019 | 2 | MAMA | 9a9a |
| ARN19G3-W5 | W | ARN19G3 | 2019 | 2 | MAMA | 9a9a |
| ARN2001-G1 | G | ARN2001 | 2020 | 2 | MDP2 | 9a9r |
| ARN2001-G10 | G | ARN2001 | 2020 | 4 | MDP2 | 9a9r |
| ARN2001-G2 | G | ARN2001 | 2020 | 2 | MDP2 | 9a9r |
| ARN2001-G3 | G | ARN2001 | 2020 | 2 | MDP2 | 9a9r |
| ARN2001-G4 | G | ARN2001 | 2020 | 2 | MDP2 | 9a9r |
| ARN2001-G5 | G | ARN2001 | 2020 | 2 | MDP2 | 9a9r |
| ARN2001-G6 | G | ARN2001 | 2020 | 2 | MDP2 | 9a9r |
| ARN2001-G7 | G | ARN2001 | 2020 | 2 | MAP2 | 9a9r |
| ARN2001-G8 | G | ARN2001 | 2020 | 2 | MDP2 | 9a9r |
| ARN2001-G9 | G | ARN2001 | 2020 | 4 | MDP2 | 9a9r |
| ARN2001-W1 | W | ARN2001 | 2020 | 2 | MAMA | 9a9a |
| ARN2001-W2 | W | ARN2001 | 2020 | 2 | MAMD | 9a9a |
| ARN2001-W3 | W | ARN2001 | 2020 | 2 | MAMD | 9a9a |
| ARN2001-W4 | W | ARN2001 | 2020 | 2 | MAMD | 9a9a |
| ARN2001-W5 | W | ARN2001 | 2020 | 2 | MAMA | 9a9a |
| ARN2002-W1 | W | ARN2002 | 2020 | 2 | MAP1 | 9a9a |
| ARN2002-W2 | W | ARN2002 | 2020 | 2 | MAP1 | 9a9a |
| ARN2002-W3 | W | ARN2002 | 2020 | 2 | MAP1 | 9a9a |
| ARN2002-W4 | W | ARN2002 | 2020 | 2 | MAP1 | 9a9a |
| ARN2002-W5 | W | ARN2002 | 2020 | 2 | MAP1 | 9a9a |
| ARN2101-G1 | G | ARN2101 | 2021 | 4 | MAP2 | 9a9r |
| ARN2101-G2 | G | ARN2101 | 2021 | 4 | P1P2 | 9a9r |
| ARN2101-G3 | G | ARN2101 | 2021 | 4 | P2P2 | 9a9r |
| ARN2101-G4 | G | ARN2101 | 2021 | 4 | P2P2 | 9a9r |
| ARN2101-G5 | G | ARN2101 | 2021 | 4 | P1P2 | 9a9r |
| ARN2101-G6 | G | ARN2101 | 2021 | 4 | P2P2 | 9a9r |

| | | | | | | |
|---|---|---|---|---|---|---|
| ARN2101-G7 | G | ARN2101 | 2021 | 4 | P2P2 | 9a9r |
| ARN2101-G8 | G | ARN2101 | 2021 | 4 | P2P2 | 9r9r |
| ARN2101-M1 | M | ARN2101 | 2021 | 4 | P2 | 9r9r |
| ARN2101-M2 | M | ARN2101 | 2021 | 4 | P2 | 9r9r |
| ARN2101-M3 | M | ARN2101 | 2021 | 4 | P2 | 9r9r |
| ARN2101-W1g | W | ARN2101 | 2021 | 4 | MAP1 | 9a9a |
| ARN2102-G2 | G | ARN2102 | 2021 | 4 | MAP1 | 9a9r |
| ARN2102-G3 | G | ARN2102 | 2021 | 4 | P1P2 | 9a9r |
| ARN2102-G4 | G | ARN2102 | 2021 | 4 | P2P2 | 9a9r |
| ARN2102-G5 | G | ARN2102 | 2021 | 4 | P1P2 | 9a9r |
| ARN2102-G6 | G | ARN2102 | 2021 | 4 | P2P2 | 9a9r |
| ARN2102-G7 | G | ARN2102 | 2021 | 4 | P2P2 | 9a9r |
| ARN2102-G8 | G | ARN2102 | 2021 | 4 | P1P2 | 9a9r |
| ARN2102-M1 | M | ARN2102 | 2021 | 4 | P2 | 9a9r |
| ARN2102-M2 | M | ARN2102 | 2021 | 4 | P2 | 9r9r |
| ARN2102-M3 | M | ARN2102 | 2021 | 4 | P2 | 9r9r |
| ARN2102-M4 | M | ARN2102 | 2021 | 4 | P2 | 9r9r |
| ARN2102-M5 | M | ARN2102 | 2021 | 4 | P2 | 9r9r |
| ARN2102-W2 | W | ARN2102 | 2021 | 3 | MAP1 | 9a9a |
| ARN2102-W4 | W | ARN2102 | 2021 | 3 | MAP1 | 9a9a |
| ARN2102-W5 | W | ARN2102 | 2021 | 3 | MAP1 | 9a9a |
| ARN2103-W1 | W | ARN2103 | 2021 | 3 | MAMA | 9a9a |
| ARN2103-W2 | W | ARN2103 | 2021 | 3 | MAMA | 9a9a |
| ARN2103-W3 | W | ARN2103 | 2021 | 3 | MAMA | 9a9a |
| ARN2103-W4 | W | ARN2103 | 2021 | 3 | MAMA | 9a9a |
| ARN2103-W5 | W | ARN2103 | 2021 | 3 | MAMA | 9a9a |
| ARN2104-W1 | W | ARN2104 | 2021 | 3 | MAP2 | 9a9r |
| ARN2104-W2 | W | ARN2104 | 2021 | 3 | MDP2 | 9a9r |
| ARN2104-W3 | W | ARN2104 | 2021 | 3 | MDP2 | 9a9r |
| ARNC1-W1 | W | ARNC1 | 2019 | 1 | MAMA | 9a9a |
| ARNC1-W2 | W | ARNC1 | 2019 | 1 | MAMD | 9a9a |
| ARNC1-W3 | W | ARNC1 | 2019 | 1 | MAMA | 9a9a |
| ARNC1-W4 | W | ARNC1 | 2019 | 1 | MAMD | 9a9a |
| ARNC1-W5 | W | ARNC1 | 2019 | 1 | MAMD | 9a9a |
| ARNC2-W1 | W | ARNC2 | 2019 | 1 | MAMA | 9a9a |
| ARNC2-W2 | W | ARNC2 | 2019 | 1 | MAMA | 9a9a |
| ARNC2-W3 | W | ARNC2 | 2019 | 1 | MAMA | 9a9a |
| ARNC2-W4 | W | ARNC2 | 2019 | 1 | MAMA | 9a9a |
| ARNC2-W5 | W | ARNC2 | 2019 | 1 | MAMA | 9a9a |
| ARNC3-W1 | W | ARNC3 | 2019 | 1 | MAMA | 9a9a |
| ARNC3-W2 | W | ARNC3 | 2019 | 1 | MAMD | 9a9a |
| ARNC3-W3 | W | ARNC3 | 2019 | 1 | MAMD | 9a9a |
| ARNC3-W4 | W | ARNC3 | 2019 | 1 | MAMD | 9a9a |
| ARNC3-W5 | W | ARNC3 | 2019 | 1 | MAMD | 9a9a |

| | | | | | | |
|---|---|---|---|---|---|---|
| ARN-G2A | MQ | - | 2019 | 4 | MAMA | 9a9a |
| ARN-G2B-MQ | MQ | - | 2019 | 4 | MAMA | 9a9a |
| ARN-G4.2-MQ | MQ | - | 2019 | 4 | MAP1 | 9a9a |
| ARN-G4.3 | MQ | - | 2019 | 4 | MAP1 | 9a9a |
| DOM19C5-M1 | M | DOM19C5 | 2019 | 2 | MA | 9a9a |
| DOM19C5-MP6 | M | DOM19C5 | 2019 | 2 | MA | 9a9a |
| DOM19C8-M1 | M | DOM19C8 | 2019 | 2 | MA | 9a9a |
| DOM19C9-M1 | M | DOM19C9 | 2019 | 2 | MA | 9a9a |
| DOM19C9-M2 | M | DOM19C9 | 2019 | 2 | MA | 9a9a |
| DOM19C9-M4 | M | DOM19C9 | 2019 | 2 | unk | 9a9a |
| DOM19C9-M5 | M | DOM19C9 | 2019 | 2 | MA | 9a9a |
| DOM19C9-M6 | M | DOM19C9 | 2019 | 2 | MA | 9a9a |
| DOM19C9-M7 | M | DOM19C9 | 2019 | 2 | MA | 9a9a |
| DOM19C9-M8 | M | DOM19C9 | 2019 | 2 | MA | 9a9a |
| DOM2101-W1 | W | DOM2101 | 2021 | 3 | MAMA | 9a9a |
| DOM2101-W2 | W | DOM2101 | 2021 | 4 | MAMA | 9a9a |
| DOM2101-W3 | W | DOM2101 | 2021 | 4 | MAMA | 9a9a |
| DOM2101-W4 | W | DOM2101 | 2021 | 4 | MAMA | 9a9a |
| DOM2101-W5 | W | DOM2101 | 2021 | 4 | MAMA | 9a9a |
| DOM2102-G1 | G | DOM2102 | 2021 | 4 | MAMD | 9a9a |
| DOM2102-G2 | G | DOM2102 | 2021 | 4 | MAMD | 9a9a |
| DOM2102-G3 | G | DOM2102 | 2021 | 4 | MAMD | 9a9a |
| DOM2102-W1 | W | DOM2102 | 2021 | 3 | MAMD | 9a9a |
| DOM2102-W2 | W | DOM2102 | 2021 | 4 | MAMA | 9a9a |
| DOM2102-W3 | W | DOM2102 | 2021 | 4 | MAMD | 9a9a |
| DOM2102-W4 | W | DOM2102 | 2021 | 4 | MAMD | 9a9a |
| DOM2102-W5 | W | DOM2102 | 2021 | 4 | MAMA | 9a9a |
| DOM2103-G1 | G | DOM2103 | 2021 | 4 | MAMA | 9a9a |
| DOM2103-G2 | G | DOM2103 | 2021 | 4 | MAMA | 9a9a |
| DOM2103-G3 | G | DOM2103 | 2021 | 4 | MAMA | 9a9a |
| DOM2103-G4 | G | DOM2103 | 2021 | 4 | MAMA | 9a9a |
| DOM2103-G5 | G | DOM2103 | 2021 | 4 | MAMA | 9a9a |
| DOM2103-G6 | G | DOM2103 | 2021 | 4 | MAMA | 9a9a |
| DOM2103-G7 | G | DOM2103 | 2021 | 4 | MAMA | 9a9a |
| DOM2103-G8 | G | DOM2103 | 2021 | 4 | MAMA | 9a9a |
| DOM2103-W1 | W | DOM2103 | 2021 | 3 | MAMA | 9a9a |
| DOM2103-W2 | W | DOM2103 | 2021 | 4 | MAMA | 9a9a |
| DOM2103-W3 | W | DOM2103 | 2021 | 4 | MAMA | 9a9a |
| DOM2103-W4 | W | DOM2103 | 2021 | 4 | MAMA | 9a9a |
| DOM2103-W5 | W | DOM2103 | 2021 | 4 | MAMA | 9a9a |
| DOM2104-W1 | W | DOM2104 | 2021 | 3 | MAMA | 9a9a |
| DOM2104-W2 | W | DOM2104 | 2021 | 4 | MAMA | 9a9a |
| DOM2104-W3 | W | DOM2104 | 2021 | 4 | MAMA | 9a9a |
| DOM2104-W4 | W | DOM2104 | 2021 | 4 | MAMA | 9a9a |

| | | | | | | |
|---|---|---|---|---|---|---|
| DOM2104-W5 | W | DOM2104 | 2021 | 4 | MAMA | 9a9a |
| DOM2105-G1 | G | DOM2105 | 2021 | 4 | MAMD | 9a9a |
| DOM2105-G2 | G | DOM2105 | 2021 | 4 | MAMD | 9a9a |
| DOM2105-G3 | G | DOM2105 | 2021 | 4 | MAMD | 9a9a |
| DOM2105-G4 | G | DOM2105 | 2021 | 4 | MAMD | 9a9a |
| DOM2105-G5 | G | DOM2105 | 2021 | 4 | MAMD | 9a9a |
| DOM2105-G6 | G | DOM2105 | 2021 | 4 | MAMD | 9a9a |
| DOM2105-G7 | G | DOM2105 | 2021 | 4 | MAMD | 9a9a |
| DOM2105-G8 | G | DOM2105 | 2021 | 4 | MAMA | 9a9a |
| DOM2105-W2 | W | DOM2105 | 2021 | 4 | MAMA | 9a9a |
| DOM2105-W3 | W | DOM2105 | 2021 | 4 | MAMD | 9a9a |
| DOM2105-W4 | W | DOM2105 | 2021 | 4 | MAMD | 9a9a |
| DOM2105-W5 | W | DOM2105 | 2021 | 4 | MAMA | 9a9a |
| DOM2106-G1 | G | DOM2106 | 2021 | 4 | MAMA | 9a9a |
| DOM2106-G2 | G | DOM2106 | 2021 | 4 | MAMA | 9a9a |
| DOM2106-G3 | G | DOM2106 | 2021 | 4 | MAMD | 9a9a |
| DOM2106-W1 | W | DOM2106 | 2021 | 3 | MAMA | 9a9a |
| DOM2106-W2 | W | DOM2106 | 2021 | 4 | MAMA | 9a9a |
| DOM2106-W3 | W | DOM2106 | 2021 | 4 | MAMA | 9a9a |
| DOM2106-W4 | W | DOM2106 | 2021 | 4 | MAMD | 9a9a |
| DOM2106-W5 | W | DOM2106 | 2021 | 4 | MAMD | 9a9a |
| DOM2107-M1 | M | DON2107 | 2021 | 4 | MA | 9a9a |
| DOM2108-M1 | M | DOM2108 | 2021 | 4 | MA | 9a9a |
| DOM2108-M2 | M | DOM2108 | 2021 | 4 | MA | 9a9a |
| DOM2108-W1 | W | DOM2108 | 2021 | 3 | MAMA | 9a9a |
| DOM2108-W2 | W | DOM2108 | 2021 | 4 | MAMA | 9a9a |
| DOM2108-W3 | W | DOM2108 | 2021 | 4 | MAMA | 9a9a |
| DOM2108-W4 | W | DOM2108 | 2021 | 4 | MAMA | 9a9a |
| DOM2108-W5 | W | DOM2108 | 2021 | 4 | MAMA | 9a9a |
| DOMPupa2spupa1 | W | - | | 2 | MAMD | 9a9a |
| DOMPupa2-W3 | W | - | | 2 | MAMA | 9a9a |
| DOMPupa2-W4 | W | - | | 2 | MAMD | 9a9a |
| DOMPupa3spupa1 | W | - | | 2 | MAMA | 9a9a |
| DOMPupa3-W1 | W | - | | 2 | MAMA | 9a9a |
| DOMPupa3-W2 | W | - | | 2 | MAMA | 9a9a |
| DOMPupa3-W3 | W | - | | 2 | MAMA | 9a9a |
| DOMPupa3-W4 | W | - | | 2 | MAMA | 9a9a |
| DOMPupa4spupa1 | W | - | | 2 | MAMD | 9a9a |
| DOMPupa4-W1 | W | - | | 2 | MAMD | 9a9a |
| DOMPupa4-W2 | W | - | | 2 | MAMA | 9a9a |
| DOMPupa4-W3 | W | - | | 2 | MAMA | 9a9a |
| DON18C1-M1 | M | DON19C1 | 2018 | 2 | MA | 9a9a |

| | | | | | | |
|---|---|---|---|---|---|---|
| DON18C1-W1 | W | DON19C1 | 2018 | 1 | MAMA | 9a9a |
| DON18C1-W2 | W | DON19C1 | 2018 | 1 | MAMA | 9a9a |
| DON18C1-W3 | W | DON19C1 | 2018 | 1 | MAMA | 9a9a |
| DON18C1-W4 | W | DON19C1 | 2018 | 1 | MAMA | 9a9a |
| DON18C1-W5 | W | DON19C1 | 2018 | 1 | MAMA | 9a9a |
| DON18C3-G1 | G | DON18C3 | 2018 | 2 | MAMA | 9a9a |
| DON18C3-G2 | G | DON18C3 | 2018 | 2 | MAMD | 9a9a |
| DON18C3-G3 | G | DON18C3 | 2018 | 2 | unk | 9a9a |
| DON18C3-G4 | G | DON18C3 | 2018 | 2 | MAMA | 9a9a |
| DON18C3-W1 | W | DON18C3 | 2018 | 1 | MAMA | 9a9a |
| DON18C3-W2 | W | DON18C3 | 2018 | 1 | MAMA | 9a9a |
| DON18C3-W3 | W | DON18C3 | 2018 | 1 | MAMD | 9a9a |
| DON18C3-W4 | W | DON18C3 | 2018 | 2 | MAMA | 9a9a |
| DON18C3-W5 | W | DON18C3 | 2018 | 1 | MAMA | 9a9a |
| DON19C2-G1 | G | DON19C2 | 2019 | 2 | MAMA | 9a9a |
| DON19C2-G2 | G | DON19C2 | 2019 | 2 | MAMA | 9a9a |
| DON19C2-G3 | G | DON19C2 | 2019 | 2 | MAMA | 9a9a |
| DON19C2-G4 | G | DON19C2 | 2019 | 2 | MAMA | 9a9a |
| DON19C2-G5 | G | DON19C2 | 2019 | 2 | MAMA | 9a9a |
| DON19C2-G6 | G | DON19C2 | 2019 | 2 | MAMA | 9a9a |
| DON19C2-G7 | G | DON19C2 | 2019 | 2 | MAMA | 9a9a |
| DON19C2-W1 | W | DON19C2 | 2019 | 2 | MAMA | 9a9a |
| DON19C2-W2 | W | DON19C2 | 2019 | 2 | MAMA | 9a9a |
| DON19C2-W3 | W | DON19C2 | 2019 | 2 | MAMD | 9a9a |
| DON19C2-W4 | W | DON19C2 | 2019 | 2 | MAMA | 9a9a |
| DON19C2-W5 | W | DON19C2 | 2019 | 2 | MAMD | 9a9a |
| DON19C3-M1 | M | DON19C3 | 2019 | 2 | MA | 9a9a |
| DON19C3-M2 | M | DON19C3 | 2019 | 2 | MA | 9a9a |
| DON19C3-M3 | M | DON19C3 | 2019 | 2 | MA | 9a9a |
| DON19C7-G1 | G | DON19C7 | 2019 | 2 | MAMA | 9a9a |
| DON19M2-W1 | W | DON19M2 | 2019 | 2 | MAMA | 9a9a |
| DON19M2-W2 | W | DON19M2 | 2019 | 2 | MAMA | 9a9a |
| DON19M2-W3 | W | DON19M2 | 2019 | 2 | MAMA | 9a9a |
| DON19M2-W4 | W | DON19M2 | 2019 | 2 | MAMA | 9a9a |
| DON19M2-W5 | W | DON19M2 | 2019 | 2 | MAMA | 9a9a |
| DON2101-G1 | G | DON2101 | 2021 | 4 | MAMA | 9a9a |
| DON2101-G2 | G | DON2101 | 2021 | 4 | MAMD | 9a9a |
| DON2101-G3 | G | DON2101 | 2021 | 4 | MAMA | 9a9a |
| DON2101-G4 | G | DON2101 | 2021 | 4 | MAMA | 9a9a |
| DON2101-G5 | G | DON2101 | 2021 | 4 | MAMD | 9a9a |
| DON2101-G6 | G | DON2101 | 2021 | 4 | MAMA | 9a9a |
| DON2101-G7 | G | DON2101 | 2021 | 4 | MAMD | 9a9a |
| DON2101-G8 | G | DON2101 | 2021 | 4 | MAMD | 9a9a |
| DON2101-M1 | M | DON2102 | 2021 | 4 | MA | 9a9a |

| | | | | | | |
|---|---|---|---|---|---|---|
| DON2101-M1r | M | DON2101 | 2021 | 4 | MD | 9a9a |
| DON2101-M2 | M | DON2102 | 2021 | 4 | MA | 9a9a |
| DON2101-M3 | M | DON2102 | 2021 | 4 | MA | 9a9a |
| DON2101-M4 | M | DON2102 | 2021 | 4 | MA | 9a9a |
| DON2101-W1 | W | DON2101 | 2021 | 3 | MAMA | 9a9a |
| DON2101-W2 | W | DON2101 | 2021 | 3 | MAMA | 9a9a |
| DON2101-W3 | W | DON2101 | 2021 | 3 | MAMD | 9a9a |
| DON2101-W4 | W | DON2101 | 2021 | 3 | MAMD | 9a9a |
| DON2101-W5 | W | DON2101 | 2021 | 3 | MAMD | 9a9a |
| DON2102-G1 | G | DON2102 | 2021 | 4 | MAMA | 9a9a |
| DON2107-W1 | W | DON2107 | 2021 | 3 | MAMA | 9a9a |
| DON2107-W2 | W | DON2107 | 2021 | 3 | MAMA | 9a9a |
| DON2107-W3 | W | DON2107 | 2021 | 3 | MAMA | 9a9a |
| DON2107-W4 | W | DON2107 | 2021 | 3 | MAMA | 9a9a |
| DON2107-W5 | W | DON2107 | 2021 | 3 | MAMA | 9a9a |
| DONC2-W1 | W | DONC2 | 2019 | 1 | MAMA | 9a9a |
| DONC2-W2 | W | DONC2 | 2019 | 1 | MAMA | 9a9a |
| DONC2-W3 | W | DONC2 | 2019 | 1 | MAMA | 9a9a |
| DONC2-W4 | W | DONC2 | 2019 | 1 | MAMA | 9a9a |
| DONC2-W5 | W | DONC2 | 2019 | 1 | MAMA | 9a9a |
| DONC4-W1 | W | DONC4 | 2019 | 1 | MAMA | 9a9a |
| DONC4-W2 | W | DONC4 | 2019 | 1 | MAMA | 9a9a |
| DONC4-W3 | W | DONC4 | 2019 | 1 | MAMA | 9a9a |
| DONC4-W4 | W | DONC4 | 2019 | 1 | MAMA | 9a9a |
| DONC4-W5 | W | DONC4 | 2019 | 1 | MAMA | 9a9a |
| DONC5-W1 | W | DONC5 | 2019 | 1 | MAMA | 9a9a |
| DONC5-W2 | W | DONC5 | 2019 | 1 | MAMA | 9a9a |
| DONC5-W3 | W | DONC5 | 2019 | 1 | MAMA | 9a9a |
| DONC5-W4 | W | DONC5 | 2019 | 1 | MAMA | 9a9a |
| DONC5-W5 | W | DONC5 | 2019 | 1 | MAMA | 9a9a |
| DONNAS-G1-MQ | MQ | - | 2019 | 4 | P2P2 | 9r9r |
| HONE19C1-M1 | M | HONE19C1 | 2019 | 2 | P2 | 9r9r |
| HONE19C1-M2 | M | HONE19C1 | 2019 | 2 | P2 | 9r9r |
| HONE19C1-W1 | W | HONE19C1 | 2019 | 2 | P1P2 | 9a9r |
| HONE19C1-W2 | W | HONE19C1 | 2019 | 2 | MAP2 | 9a9r |
| HONE19C1-W3 | W | HONE19C1 | 2019 | 2 | P1P2 | 9r9r |
| HONE19C1-W4 | W | HONE19C1 | 2019 | 2 | P1P2 | 9a9r |
| HONE19C1-W5 | W | HONE19C1 | 2019 | 2 | P1P2 | 9a9r |
| MAS2101-W2 | W | MAS2101 | 2021 | 4 | MAMA | 9a9a |
| MAS2101-W3 | W | MAS2101 | 2021 | 4 | MAMA | 9a9a |
| MAS2101-W4 | W | MAS2101 | 2021 | 4 | MAMA | 9a9a |
| MAS2101-W5 | W | MAS2101 | 2021 | 4 | MAMA | 9a9a |
| MAS2102-G1 | G | MAS2102 | 2021 | 4 | MAMA | 9a9a |
| MAS2102-G2 | G | MAS2102 | 2021 | 4 | MAMD | 9a9a |

| | | | | | | |
|---|---|---|---|---|---|---|
| MAS2102-G3 | G | MAS2102 | 2021 | 4 | MAMD | 9a9a |
| MAS2102-G4 | G | MAS2102 | 2021 | 4 | MAMD | 9a9a |
| MAS2102-G5 | G | MAS2102 | 2021 | 4 | MAMD | 9a9a |
| MAS2102-G6 | G | MAS2102 | 2021 | 4 | MAMD | 9a9a |
| MAS2102-G7 | G | MAS2102 | 2021 | 4 | MAMA | 9a9a |
| MAS2102-G8 | G | MAS2102 | 2021 | 4 | MAMA | 9a9a |
| MAS2102-W1 | W | MAS2102 | 2021 | 4 | MAMA | 9a9a |
| MAS2102-W2 | W | MAS2102 | 2021 | 4 | MAMD | 9a9a |
| MAS2102-W3 | W | MAS2102 | 2021 | 4 | MAMA | 9a9a |
| MAS2102-W4 | W | MAS2102 | 2021 | 4 | MAMD | 9a9a |
| MAS2102-W5 | W | MAS2102 | 2021 | 4 | MAMD | 9a9a |
| MJO19C1-G1 | G | MJO19C1 | 2019 | 2 | MDP1 | 9a9a |
| MJO19C1-G2 | G | MJO19C1 | 2019 | 2 | P1P1 | 9a9a |
| MJO19C1-M1 | M | MJO19C1 | 2019 | 2 | P1 | 9a9a |
| MJO19C1-W1 | W | MJO19C1 | 2019 | 2 | MAP1 | 9a9a |
| MJO19C1-W2 | W | MJO19C1 | 2019 | 2 | MAP1 | 9a9a |
| MJO19C1-W3 | W | MJO19C1 | 2019 | 2 | MAP1 | 9a9a |
| MJO19C1-W4 | W | MJO19C1 | 2019 | 2 | P1P1 | 9a9a |
| MJO19C1-W5 | W | MJO19C1 | 2019 | 2 | MAP1 | 9a9a |
| MJO19C2-G2 | G | MJO19C2 | 2019 | 2 | MAP1 | 9a9a |
| MJO19C2-M1 | M | MJO19C2 | 2019 | 2 | P1 | 9a9a |
| MJO19C2-M2 | M | MJO19C2 | 2019 | 2 | P1 | 9a9a |
| MJO19C2-M3 | M | MJO19C2 | 2019 | 2 | P1 | 9a9a |
| MJO19C2-W1 | W | MJO19C2 | 2019 | 2 | MDP1 | 9a9a |
| MJO19C2-W2 | W | MJO19C2 | 2019 | 2 | P1P1 | 9a9a |
| MJO19C2-W3 | W | MJO19C2 | 2019 | 2 | MDP1 | 9a9a |
| MJO19C2-W4 | W | MJO19C2 | 2019 | 2 | P1P1 | 9a9a |
| MJO19C2-W5 | W | MJO19C2 | 2019 | 2 | MDP1 | 9a9a |
| MJO19C3-G2 | G | MJO19C3 | 2019 | 2 | MAP1 | 9a9a |
| MJO19C3-G3 | G | MJO19C3 | 2019 | 2 | MDP1 | 9a9a |
| MJO19C3-G4 | G | MJO19C3 | 2019 | 2 | MAP1 | 9a9a |
| MJO19C3-G5 | G | MJO19C3 | 2019 | 2 | MAP1 | 9a9a |
| MJO19C3-M1 | M | MJO19C3 | 2019 | 2 | P1 | 9a9a |
| MJO19C3-M2 | M | MJO19C3 | 2019 | 2 | P1 | 9a9a |
| MJO19C3-M3 | M | MJO19C3 | 2019 | 2 | P1 | 9a9a |
| MJO19C3-M4 | M | MJO19C3 | 2019 | 2 | P1 | 9a9a |
| MJO19C3-M5 | M | MJO19C3 | 2019 | 2 | P1 | 9a9a |
| MJO19C3-M7 | M | MJO19C3 | 2019 | 2 | P1 | 9a9a |
| MJO19C3-W1 | W | MJO19C3 | 2019 | 2 | P1P1 | 9a9a |
| MJO19C3-W2 | W | MJO19C3 | 2019 | 2 | MAP1 | 9a9a |
| MJO19C3-W3 | W | MJO19C3 | 2019 | 2 | MAP1 | 9a9a |
| MJO19C3-W4 | W | MJO19C3 | 2019 | 2 | P1P1 | 9a9a |
| MJO19C3-W5 | W | MJO19C3 | 2019 | 2 | MDP1 | 9a9a |
| MJO19C4-G1 | G | MJO19C4 | 2019 | 2 | MDP1 | 9a9a |

| | | | | | | |
|---|---|---|---|---|---|---|
| MJO19C4-W1 | W | MJO19C4 | 2019 | 2 | MDP1 | 9a9a |
| MJO19C4-W2 | W | MJO19C4 | 2019 | 2 | MDP1 | 9a9a |
| MJO19C4-W3 | W | MJO19C4 | 2019 | 2 | MDP1 | 9a9a |
| MJO19C4-W4 | W | MJO19C4 | 2019 | 2 | MDP1 | 9a9a |
| MJO19C4-W5 | W | MJO19C4 | 2019 | 2 | MDP1 | 9a9a |
| MJO19C5-G1 | G | MJO19C5 | 2019 | 2 | MDP1 | 9a9a |
| MJO19C5-M1 | M | MJO19C5 | 2019 | 2 | P1 | 9a9a |
| MJO19C5-M3 | M | MJO19C5 | 2019 | 2 | P1 | 9a9a |
| MJO19C5-M4 | M | MJO19C5 | 2019 | 2 | P1 | 9a9a |
| MJO19C5-M5 | M | MJO19C5 | 2019 | 2 | P1 | 9a9a |
| MJO19C5-M6 | M | MJO19C5 | 2019 | 2 | P1 | 9a9a |
| MJO19C5-M7 | M | MJO19C5 | 2019 | 2 | P1 | 9a9a |
| MJO19C5-Q1 | Q | MJO19C5 | 2019 | 2 | P1P1 | 9a9a |
| MJO19C5-Q2 | Q | MJO19C5 | 2019 | 2 | P1P1 | 9a9a |
| MJO19C5-Q3 | Q | MJO19C5 | 2019 | 2 | P1P1 | 9a9a |
| MJO19C5-W1 | W | MJO19C5 | 2019 | 2 | MAP1 | 9a9a |
| MJO19C5-W2 | W | MJO19C5 | 2019 | 2 | MAP1 | 9a9a |
| MJO19C5-W3 | W | MJO19C5 | 2019 | 2 | MDP1 | 9a9a |
| MJO19C5-W4 | W | MJO19C5 | 2019 | 2 | MDP1 | 9a9a |
| MJO19C5-W5 | W | MJO19C5 | 2019 | 2 | MDP1 | 9a9a |
| MJO19C6-M1 | M | MJO19C6 | 2019 | 2 | MA | 9a9a |
| MJO19C6-M2 | M | MJO19C6 | 2019 | 2 | MA | 9a9a |
| MJO19C6-M3 | M | MJO19C6 | 2019 | 2 | MA | 9a9a |
| MJO19C6-M4 | M | MJO19C6 | 2019 | 2 | MA | 9a9a |
| MJO19C6-M5 | M | MJO19C6 | 2019 | 2 | MA | 9a9a |
| MJO19C6-W1 | W | MJO19C6 | 2019 | 2 | MAMA | 9a9a |
| MJO19C6-W2 | W | MJO19C6 | 2019 | 2 | MAMA | 9a9a |
| MJO19C6-W3 | W | MJO19C6 | 2019 | 2 | MAMD | 9a9a |
| MJO19C6-W4 | W | MJO19C6 | 2019 | 2 | MAMD | 9a9a |
| MJO19C6-W5 | W | MJO19C6 | 2019 | 2 | MAMD | 9a9a |
| MJO19C7-M1 | M | MJO19C7 | 2019 | 2 | P2 | 9r9r |
| MJO19C7-M2 | M | MJO19C7 | 2019 | 2 | P2 | 9r9r |
| MJO19C7-M3 | M | MJO19C7 | 2019 | 2 | P2 | 9r9r |
| MJO19C7-M4 | M | MJO19C7 | 2019 | 2 | P2 | 9r9r |
| MJO19C7-M5 | M | MJO19C7 | 2019 | 2 | P2 | 9r9r |
| MJO19C7-M6 | M | MJO19C7 | 2019 | 2 | P2 | 9r9r |
| MJO19C7-W1 | W | MJO19C7 | 2019 | 2 | MAP2 | 9a9r |
| MJO19C7-W2 | W | MJO19C7 | 2019 | 2 | MAP2 | 9a9a |
| MJO19C7-W3 | W | MJO19C7 | 2019 | 2 | MAP2 | 9r9r |
| MJO19C7-W4 | W | MJO19C7 | 2019 | 2 | MAP2 | 9a9r |
| MJO19C7-W4r | W | MJO19C7 | 2019 | 2 | MAP2 | 9a9r |
| MJO19C7-W5 | W | MJO19C7 | 2019 | 2 | MAP2 | 9a9a |
| MJO19C7-W5r | W | MJO19C7 | 2019 | 2 | MAP2 | 9a9a |
| MJO19C8-M1 | M | MJO19C8 | 2019 | 2 | MA | 9a9a |

| | | | | | | |
|---|---|---|---|---|---|---|
| MJO19C8-W1 | W | MJO19C8 | 2019 | 2 | MAMA | 9a9a |
| MJO19C8-W2 | W | MJO19C8 | 2019 | 2 | MAMA | 9a9a |
| MJO19C8-W3 | W | MJO19C8 | 2019 | 2 | MAMA | 9a9a |
| MJO19C8-W4 | W | MJO19C8 | 2019 | 2 | MAMA | 9a9a |
| MJO19C8-W5 | W | MJO19C8 | 2019 | 2 | MAMA | 9a9a |
| MJO19C9-G1 | G | MJO19C9 | 2019 | 2 | MDP1 | 9a9a |
| MJO19C9-M1 | M | MJO19C9 | 2019 | 2 | P1 | 9a9a |
| MJO19C9-Q1 | Q | MJO19C9 | 2019 | 2 | P1P1 | 9a9a |
| MJO19C9-W1 | W | MJO19C9 | 2019 | 2 | MAP1 | 9a9a |
| MJO19C9-W2 | W | MJO19C9 | 2019 | 2 | MAP1 | 9a9a |
| MJO19C9-W3 | W | MJO19C9 | 2019 | 2 | MDP1 | 9a9a |
| MJO19C9-W4 | W | MJO19C9 | 2019 | 2 | MDP1 | 9a9a |
| MJO19C9-W5 | W | MJO19C9 | 2019 | 2 | MDP1 | 9a9a |
| MJO19G2-Q1 | Q | MJO19G2 | 2019 | 2 | MAMA | 9a9a |
| MJO19G4-Q1 | Q | MJO19G4 | 2019 | 2 | MAP1 | 9a9a |
| MJO19G5-Q1 | Q | MJO19G5 | 2019 | 2 | MAMA | 9a9a |
| MJO2001-M1 | M | MJO2001 | 2020 | 2 | MA | 9a9a |
| MJO2001-M2 | M | MJO2001 | 2020 | 2 | MA | 9a9a |
| MJO2001-M3 | M | MJO2001 | 2020 | 2 | MA | 9a9a |
| MJO2001-W1 | W | MJO2001 | 2020 | 2 | MAMA | 9a9a |
| MJO2001-W2 | W | MJO2001 | 2020 | 2 | MAMA | 9a9a |
| MJO2001-W5 | W | MJO2001 | 2020 | 2 | MAMA | 9a9a |
| MJO2003-M1 | M | MJO2003 | 2020 | 2 | P1 | 9a9a |
| MJO2003-M2 | M | MJO2003 | 2020 | 2 | MDP1 | 9a9a |
| MJO2003-W1 | W | MJO2003 | 2020 | 2 | P1P1 | 9a9a |
| MJO2003-W2 | W | MJO2003 | 2020 | 2 | P1P1 | 9a9a |
| MJO2003-W3 | W | MJO2003 | 2020 | 2 | P1P1 | 9a9a |
| MJO2003-W4 | W | MJO2003 | 2020 | 2 | MAP1 | 9a9a |
| MJO2003-W5 | W | MJO2003 | 2020 | 2 | MDP1 | 9a9a |
| MJO2004-M2 | M | MJO2004 | 2020 | 2 | MA | 9a9a |
| MJO2004-M3 | M | MJO2004 | 2020 | 2 | MA | 9a9a |
| MJO2004-M4 | M | MJO2004 | 2020 | 2 | MA | 9a9a |
| MJO2004-M5 | M | MJO2004 | 2020 | 2 | MA | 9a9a |
| MJO2004-M6 | M | MJO2004 | 2020 | 2 | MA | 9a9a |
| MJO2004-M7 | M | MJO2004 | 2020 | 2 | MA | 9a9a |
| MJO2004-M8 | M | MJO2004 | 2020 | 2 | MA | 9a9a |
| MJO2004-W1 | W | MJO2004 | 2020 | 2 | MAMA | 9a9a |
| MJO2004-W5 | W | MJO2004 | 2020 | 2 | MAMA | 9a9a |
| MJO2005-M1 | M | MJO2005 | 2020 | 2 | MA | 9a9a |
| MJO2005-M2 | M | MJO2005 | 2020 | 2 | MA | 9a9a |
| MJO2005-M3 | M | MJO2005 | 2020 | 2 | MAMA | 9a9a |
| MJO2005-W2 | W | MJO2005 | 2020 | 2 | MAMA | 9a9a |
| MJO2005-W3 | W | MJO2005 | 2020 | 2 | MAMA | 9a9a |
| MJO2005-W5 | W | MJO2005 | 2020 | 2 | MAMA | 9a9a |

| | | | | | | |
|---|---|---|---|---|---|---|
| MJO2006-M1 | M | MJO2006 | 2020 | 2 | MA | 9a9a |
| MJO2006-M2 | M | MJO2006 | 2020 | 2 | MA | 9a9a |
| MJO2006-M3 | M | MJO2006 | 2020 | 2 | MA | 9a9a |
| MJO2006-M4 | M | MJO2006 | 2020 | 2 | MA | 9a9a |
| MJO2006-M5 | M | MJO2006 | 2020 | 2 | MA | 9a9a |
| MJO2006-M6 | M | MJO2006 | 2020 | 2 | MA | 9a9a |
| MJO2006-M7 | M | MJO2006 | 2020 | 2 | MA | 9a9a |
| MJO2006-M8 | M | MJO2006 | 2020 | 2 | MA | 9a9a |
| MJO2006-W1 | W | MJO2006 | 2020 | 2 | MAMA | 9a9a |
| MJO2006-W2 | W | MJO2006 | 2020 | 2 | MAMA | 9a9a |
| MJO2006-W3 | W | MJO2006 | 2020 | 2 | MAMA | 9a9a |
| MJO2006-W4 | W | MJO2006 | 2020 | 2 | MAMA | 9a9a |
| MJO2006-W5 | W | MJO2006 | 2020 | 2 | MAMA | 9a9a |
| MJO2007-M5 | M | MJO2007 | 2020 | 2 | MA | 9a9a |
| MJO2007-M6 | M | MJO2007 | 2020 | 2 | MAMA | 9a9a |
| MJO2007-M7 | M | MJO2007 | 2020 | 2 | MA | 9a9a |
| MJO2007-M8 | M | MJO2007 | 2020 | 2 | MA | 9a9a |
| MJO2007-W2 | W | MJO2007 | 2020 | 2 | MAMA | 9a9a |
| MJO2007-W5 | W | MJO2007 | 2020 | 2 | MAMA | 9a9a |
| MJO20C7-M1 | M | MJO20C7 | 2020 | 2 | P2 | 9r9r |
| MJO20C7-M2 | M | MJO20C7 | 2020 | 2 | P2 | 9r9r |
| MJO20C7-M3 | M | MJO20C7 | 2020 | 2 | P2 | 9r9r |
| MJO20C7-M4 | M | MJO20C7 | 2020 | 2 | P2 | 9r9r |
| MJO20C7-M5 | M | MJO20C7 | 2020 | 2 | P2 | 9r9r |
| MJO20C7-M6 | M | MJO20C7 | 2020 | 2 | P2 | 9r9r |
| MJO20C7-M7 | M | MJO20C7 | 2020 | 2 | P2 | 9r9r |
| MJO20C7-M8 | M | MJO20C7 | 2020 | 2 | P2 | 9r9r |
| MJOC2-MQ | MQ | - | 2019 | 4 | MDP1 | 9a9a |
| MJOC5.1-1wing | MQ | - | 2019 | 4 | P1P1 | 9a9a |
| MJOC5B-MQ | MQ | - | 2019 | 4 | P1P1 | 9a9a |
| MJO-NQ1 | NQ | - | 2019 | 4 | MAMA | 9a9a |
| MJO-NQ3 | NQ | - | 2019 | 4 | MAMD | 9a9a |
| MJO-NQ8 | NQ | - | 2019 | 4 | unk | unkn |
| QUA2001-W1 | W | QUA2001 | 2020 | 2 | MAP1 | 9a9a |
| QUA2001-W2 | W | QUA2001 | 2020 | 2 | MAP1 | 9a9a |
| QUA2001-W3 | W | QUA2001 | 2020 | 2 | MAP1 | 9a9a |
| QUA2001-W4 | W | QUA2001 | 2020 | 2 | MAP1 | 9a9a |
| QUA2001-W5 | W | QUA2001 | 2020 | 2 | P1P1 | 9a9a |
| QUA2002-G1 | G | QUA2002 | 2020 | 2 | P1P1 | 9a9a |
| QUA2002-W4 | W | QUA2002 | 2020 | 2 | MAP1 | 9a9a |
| QUA2002-W5 | W | QUA2002 | 2020 | 2 | MAP1 | 9a9a |
| QUA2101-M1 | M | QUA2101 | 2021 | 4 | P1 | 9a9a |
| QUA2101-M2 | M | QUA2101 | 2021 | 4 | MA | 9a9a |
| QUA2101-M3 | M | QUA2101 | 2021 | 4 | MA | 9a9a |

| | | | | | | |
|---|---|---|---|---|---|---|
| QUA2101-M4 | M | QUA2101 | 2021 | 4 | MA | 9a9a |
| QUA2101-M5 | M | QUA2101 | 2021 | 4 | MA | 9a9a |
| QUA2101-M6 | M | QUA2101 | 2021 | 4 | MA | 9a9a |
| QUA2101-W1 | W | QUA2101 | 2021 | 3 | MAMA | 9a9a |
| QUA2101-W2 | W | QUA2101 | 2021 | 3 | MAMA | 9a9a |
| QUA2101-W3 | W | QUA2101 | 2021 | 3 | MAMA | 9a9a |
| QUA2101-W4 | W | QUA2101 | 2021 | 3 | MAMA | 9a9a |
| QUA2101-W5 | W | QUA2101 | 2021 | 3 | MAMA | 9a9a |
| QUIC11-W1 | W | QUIC11 | 2014 | 1 | MAP2 | 9a9r |
| QUIC11-W2 | W | QUIC11 | 2014 | 1 | MDP1 | 9a9a |
| QUIC11-W3 | W | QUIC11 | 2014 | 1 | MAP1 | 9a9a |
| QUIC11-W4 | W | QUIC11 | 2014 | 1 | MAP1 | 9a9a |
| QUIC11-W5 | W | QUIC11 | 2014 | 1 | MAP1 | 9a9a |
| QUIC12-W3 | W | QUIC12 | 2014 | 1 | MAP1 | 9a9r |
| QUIC12-W4 | W | QUIC12 | 2014 | 1 | MAP2 | 9a9a |
| QUIC12-W4r | W | QUIC12 | 2014 | 1 | P1P2 | 9a9r |
| QUIC12-W5 | W | QUIC12 | 2014 | 1 | MAP1 | 9a9r |
| QUIC12-W5r | W | QUIC12 | 2014 | 1 | MAP2 | 9a9a |
| QUIC13-W1 | W | QUIC13 | 2014 | 1 | MAP1 | 9a9a |
| QUIC13-W2 | W | QUIC13 | 2014 | 1 | MAP1 | 9a9a |
| QUIC13-W3 | W | QUIC13 | 2014 | 1 | MDP1 | 9a9a |
| QUIC13-W4 | W | QUIC13 | 2014 | 1 | MAP1 | 9a9a |
| QUIC13-W5 | W | QUIC13 | 2014 | 1 | MAP1 | 9a9a |
| QUIC14-W1 | W | QUIC14 | 2014 | 1 | MAMA | 9a9a |
| QUIC14-W2 | W | QUIC14 | 2014 | 1 | MAMA | 9a9a |
| QUIC14-W3 | W | QUIC14 | 2014 | 1 | MAMA | 9a9a |
| QUIC14-W4 | W | QUIC14 | 2014 | 1 | MAMA | 9a9a |
| QUIC14-W5 | W | QUIC14 | 2014 | 1 | MAMA | 9a9a |
| QUIC15-W1 | W | QUIC15 | 2014 | 1 | P1P1 | 9a9a |
| QUIC15-W2 | W | QUIC15 | 2014 | 1 | MAP1 | 9a9a |
| QUIC15-W3 | W | QUIC15 | 2014 | 1 | MAP1 | 9a9a |
| QUIC15-W4 | W | QUIC15 | 2014 | 1 | MAP1 | 9a9a |
| QUIC15-W5 | W | QUIC15 | 2014 | 1 | MAP1 | 9a9a |
| QUIC16-W1 | W | QUIC16 | 2014 | 1 | P1P2 | 9a9r |
| QUIC16-W2 | W | QUIC16 | 2014 | 1 | MAP1 | 9a9a |
| QUIC16-W3 | W | QUIC16 | 2014 | 1 | MAP1 | 9a9a |
| QUIC16-W4 | W | QUIC16 | 2014 | 1 | MAP1 | 9a9a |
| QUIC16-W5 | W | QUIC16 | 2014 | 1 | P1P2 | 9a9r |
| QUIC17-W5 | W | QUIC17 | 2014 | 1 | P1P2 | 9a9r |
| QUIC17-W6 | W | QUIC17 | 2014 | 1 | P1P1 | 9a9a |
| QUIC17-W6r | W | QUIC17 | 2014 | 1 | P1P2 | 9a9r |
| QUIC17-W7 | W | QUIC17 | 2014 | 1 | P1P2 | 9a9r |
| QUIC17-W8 | W | QUIC17 | 2014 | 1 | MAP1 | 9a9a |
| QUIC18-W5 | W | QUIC18 | 2014 | 1 | P1P2 | 9a9r |

| | | | | | | |
|---|---|---|---|---|---|---|
| QUIC18-W6 | W | QUIC18 | 2014 | 1 | MAP1 | 9a9a |
| QUIC18-W7 | W | QUIC18 | 2014 | 1 | MAP1 | 9a9a |
| QUIC18-W8 | W | QUIC18 | 2014 | 1 | MDP1 | 9a9a |
| QUIC19-W5 | W | QUIC19 | 2014 | 1 | MAP1 | 9a9a |
| QUIC19-W6 | W | QUIC19 | 2014 | 1 | MAP1 | 9a9a |
| QUIC19-W7 | W | QUIC19 | 2014 | 1 | MAP1 | 9a9a |
| QUIC19-W8 | W | QUIC19 | 2014 | 1 | MDP1 | 9a9a |
| QUIC20-W5 | W | QUIC20 | 2014 | 1 | MAMA | 9a9a |
| QUIC20-W6 | W | QUIC20 | 2014 | 1 | MAMA | 9a9a |
| QUIC20-W7 | W | QUIC20 | 2014 | 1 | MAMA | 9a9a |
| QUIC20-W8 | W | QUIC20 | 2014 | 1 | MAMA | 9a9a |
| QUIC21-W1 | W | QUIC21 | 2014 | 1 | MAP1 | 9a9a |
| QUIC21-W2 | W | QUIC21 | 2014 | 1 | MAP1 | 9a9a |
| QUIC21-W3 | W | QUIC21 | 2014 | 1 | MAP1 | 9a9a |
| QUIC21-W4 | W | QUIC21 | 2014 | 1 | MAP1 | 9a9a |
| QUIC21-W6 | W | QUIC21 | 2014 | 1 | MAP1 | 9a9a |
| QUIC22-W1 | W | QUIC22 | 2014 | 1 | MAP1 | 9a9a |
| QUIC22-W2 | W | QUIC22 | 2014 | 1 | MAP1 | 9a9a |
| QUIC22-W3 | W | QUIC22 | 2014 | 1 | MAP1 | 9a9a |
| QUIC22-W4 | W | QUIC22 | 2014 | 1 | MAP1 | 9a9a |
| QUIC22-W5 | W | QUIC22 | 2014 | 1 | MAP1 | 9a9a |
| QUIC23-W1 | W | QUIC23 | 2014 | 1 | MAMA | 9a9a |
| QUIC23-W2 | W | QUIC23 | 2014 | 1 | MAMA | 9a9a |
| QUIC23-W3 | W | QUIC23 | 2014 | 1 | MAMA | 9a9a |
| QUIC23-W4 | W | QUIC23 | 2014 | 1 | MAMA | 9a9a |
| QUIC23-W5 | W | QUIC23 | 2014 | 1 | MAMA | 9a9a |
| QUIC26-W1 | W | QUIC26 | 2014 | 1 | MAP1 | 9a9a |
| QUIC26-W2 | W | QUIC26 | 2014 | 1 | MAP1 | 9a9a |
| QUIC26-W3 | W | QUIC26 | 2014 | 1 | MAP1 | 9a9a |
| QUIC26-W4 | W | QUIC26 | 2014 | 1 | MAP1 | 9a9a |
| QUIC26-W5 | W | QUIC26 | 2014 | 1 | MAP1 | 9a9a |
| QUIC27-W1 | W | QUIC27 | 2014 | 1 | MAP1 | 9a9a |
| QUIC27-W2 | W | QUIC27 | 2014 | 1 | MAP1 | 9a9a |
| QUIC27-W3 | W | QUIC27 | 2014 | 1 | MAP1 | 9a9a |
| QUIC27-W4 | W | QUIC27 | 2014 | 1 | MAP1 | 9a9a |
| QUIC27-W5 | W | QUIC27 | 2014 | 1 | MAP1 | 9a9a |
| QUIC28-W1 | W | QUIC28 | 2014 | 1 | MDP1 | 9a9a |
| QUIC28-W2 | W | QUIC28 | 2014 | 1 | MAP1 | 9a9a |
| QUIC28-W3 | W | QUIC28 | 2014 | 1 | MAMA | 9a9a |
| QUIC28-W4 | W | QUIC28 | 2014 | 1 | MAMD | 9a9a |
| QUIC28-W5 | W | QUIC28 | 2014 | 1 | MAMD | 9a9a |
| QUIC28-W6 | W | QUIC28 | 2014 | 1 | MAMA | 9a9a |
| QUIC28-W7 | W | QUIC28 | 2014 | 1 | MAP1 | 9a9a |
| QUIC29-W1 | W | QUIC29 | 2014 | 1 | MAP1 | 9a9a |

| | | | | | | |
|---|---|---|---|---|---|---|
| QUIC29-W2 | W | QUIC29 | 2014 | 1 | MAP1 | 9a9a |
| QUIC29-W3 | W | QUIC29 | 2014 | 1 | MAP1 | 9a9a |
| QUIC29-W4 | W | QUIC29 | 2014 | 1 | MAP1 | 9a9a |
| QUIC29-W5 | W | QUIC29 | 2014 | 1 | MAP1 | 9a9a |
| QUIC30-W1 | W | QUIC30 | 2014 | 1 | MAP1 | 9a9a |
| QUIC30-W2 | W | QUIC30 | 2014 | 1 | MAP1 | 9a9a |
| QUIC30-W3 | W | QUIC30 | 2014 | 1 | P1P1 | 9a9a |
| QUIC30-W4 | W | QUIC30 | 2014 | 1 | MAP1 | 9a9a |
| QUIC30-W5 | W | QUIC30 | 2014 | 1 | MAP1 | 9a9a |
| QUIC32-W1 | W | QUIC32 | 2014 | 1 | MAMA | 9a9a |
| QUIC32-W2 | W | QUIC32 | 2014 | 1 | MAMD | 9a9a |
| QUIC32-W4 | W | QUIC32 | 2014 | 1 | MAMD | 9a9a |
| QUIC32-W5 | W | QUIC32 | 2014 | 1 | MAMA | 9a9a |
| QUIN1401-W1 | W | QUIN1401 | 2014 | 2 | MAP1 | 9a9a |
| QUIN1401-W2 | W | QUIN1401 | 2014 | 2 | MAP1 | 9a9a |
| QUIN1401-W3 | W | QUIN1401 | 2014 | 2 | MDP1 | 9a9a |
| QUIN1401-W4 | W | QUIN1401 | 2014 | 2 | MAP1 | 9a9a |
| QUIN1401-W5 | W | QUIN1401 | 2014 | 2 | MAP1 | 9a9a |
| QUIN1402-W1 | W | QUIN1402 | 2014 | 2 | P1P2 | 9a9r |
| QUIN1402-W2 | W | QUIN1402 | 2014 | 2 | MAP1 | 9a9a |
| QUIN1403-W3 | W | QUIN1403 | 2014 | 2 | MAP1 | 9a9a |
| QUIN1403-W4 | W | QUIN1403 | 2014 | 2 | P1P2 | 9a9r |
| QUIN1403-W5 | W | QUIN1403 | 2014 | 2 | P1P2 | 9a9r |
| QUIN1404-W1 | W | QUIN1404 | 2014 | 2 | P1P2 | 9a9r |
| QUIN1404-W2 | W | QUIN1404 | 2014 | 2 | MAP1 | 9a9a |
| QUIN1404-W4 | W | QUIN1404 | 2014 | 2 | P1P2 | 9a9r |
| QUIN1404-W5 | W | QUIN1404 | 2014 | 2 | P1P2 | 9a9r |
| QUIN1405-W1 | W | QUIN1405 | 2014 | 2 | MAP1 | 9a9a |
| QUIN1405-W2 | W | QUIN1405 | 2014 | 2 | P1P2 | 9a9r |
| QUIN1405-W3 | W | QUIN1405 | 2014 | 2 | MAP1 | 9a9a |
| QUIN1405-W4 | W | QUIN1405 | 2014 | 2 | MAP1 | 9a9a |
| QUIN1406-W1 | W | QUIN1406 | 2014 | 2 | MAP1 | 9a9a |
| QUIN1406-W2 | W | QUIN1406 | 2014 | 2 | MAP1 | 9a9r |
| QUIN1406-W3 | W | QUIN1406 | 2014 | 2 | MAP1 | 9a9a |
| QUIN1406-W4 | W | QUIN1406 | 2014 | 2 | P2P2 | 9r9r |
| QUIN1406-W5 | W | QUIN1406 | 2014 | 2 | P1P2 | 9a9r |
| QUIN1407-W1 | W | QUIN1407 | 2014 | 2 | MAP1 | 9a9a |
| QUIN1407-W2 | W | QUIN1407 | 2014 | 2 | P1P2 | 9a9r |
| QUIN1407-W3 | W | QUIN1407 | 2014 | 2 | MAP1 | 9a9a |
| QUIN1407-W4 | W | QUIN1407 | 2014 | 2 | MAP1 | 9a9a |
| QUIN1407-W5 | W | QUIN1407 | 2014 | 2 | P1P2 | 9a9r |
| QUIN1408-W1 | W | QUIN1408 | 2014 | 2 | MAP1 | 9a9a |
| QUIN1408-W2 | W | QUIN1408 | 2014 | 2 | MAP1 | 9a9a |
| QUIN1408-W3 | W | QUIN1408 | 2014 | 2 | P1P1 | 9a9a |

| | | | | | | |
|---|---|---|---|---|---|---|
| QUIN1408-W4 | W | QUIN1408 | 2014 | 2 | MAP1 | 9a9a |
| QUIN1408-W5 | W | QUIN1408 | 2014 | 2 | MAP1 | 9a9a |
| QUIN1409-W1 | W | QUIN1409 | 2014 | 2 | MAP1 | 9a9a |
| QUIN1409-W2 | W | QUIN1409 | 2014 | 2 | MAP1 | 9a9a |
| QUIN1409-W3 | W | QUIN1409 | 2014 | 2 | MAP1 | 9a9a |
| QUIN1410-W1 | W | QUIN1410 | 2014 | 2 | P1P2 | 9a9r |
| QUIN1410-W3 | W | QUIN1410 | 2014 | 2 | MDP1 | 9a9a |
| QUIN1410-W4 | W | QUIN1410 | 2014 | 2 | P1P1 | 9a9a |
| QUIN18C13-G1 | G | QUIN18C13 | 2018 | 2 | MDP1 | 9a9a |
| QUIN18C13-G2 | G | QUIN18C13 | 2018 | 2 | MAP1 | 9a9a |
| QUIN18C13-G3 | G | QUIN18C13 | 2018 | 2 | P1P1 | 9a9a |
| QUIN18C13-G4 | G | QUIN18C13 | 2018 | 2 | MAP1 | 9a9a |
| QUIN18C13-W1 | W | QUIN18C13 | 2018 | 2 | MDP1 | 9a9a |
| QUIN18C13-W2 | W | QUIN18C13 | 2018 | 2 | MAP1 | 9a9a |
| QUIN18C13-W3 | W | QUIN18C13 | 2018 | 2 | MAP1 | 9a9a |
| QUIN18C13-W4 | W | QUIN18C13 | 2018 | 2 | MAP1 | 9a9a |
| QUIN18C13-W5 | W | QUIN18C13 | 2018 | 2 | MAP1 | 9a9a |
| QUIN18C14-M1 | M | QUIN18C14 | 2018 | 2 | P2 | 9r9r |
| QUIN18C14-M2 | M | QUIN18C14 | 2018 | 2 | P2 | 9r9r |
| QUIN18C14-M3 | M | QUIN18C14 | 2018 | 2 | P2 | 9r9r |
| QUIN18C15-M1 | M | QUIN18C15 | 2018 | 2 | MA | 9a9a |
| QUIN18C15-M2 | M | QUIN18C15 | 2018 | 2 | MA | 9a9a |
| QUIN18C15-M3 | M | QUIN18C15 | 2018 | 2 | MA | 9a9a |
| QUIN18C15-W1 | W | QUIN18C15 | 2018 | 2 | P2P2 | 9a9r |
| QUIN18C15-W2 | W | QUIN18C15 | 2018 | 2 | P1P2 | 9a9r |
| QUIN18C15-W3 | W | QUIN18C15 | 2018 | 2 | P2P2 | 9a9r |
| QUIN18C15-W4 | W | QUIN18C15 | 2018 | 2 | P1P2 | 9a9r |
| QUIN18C15-W5 | W | QUIN18C15 | 2018 | 2 | P2P2 | 9a9r |
| QUIN19C1-M1 | M | QUIN19C1 | 2019 | 2 | MA | 9a9a |
| QUIN19C1-M2 | M | QUIN19C1 | 2019 | 2 | MA | 9a9a |
| QUIN19C2-M1 | M | QUIN19C2 | 2019 | 2 | P2 | 9r9r |
| QUIN19C2-M2 | M | QUIN19C2 | 2019 | 2 | P2 | 9r9r |
| QUIN19C2-M3 | M | QUIN19C2 | 2019 | 2 | P2 | 9r9r |
| QUIN19C2-M4 | M | QUIN19C2 | 2019 | 2 | P2 | 9r9r |
| QUIN19C2-M5 | M | QUIN19C2 | 2019 | 2 | P2 | 9r9r |
| QUIN19C2-M6 | M | QUIN19C2 | 2019 | 2 | P2 | 9r9r |
| QUIN19C2-M7 | M | QUIN19C2 | 2019 | 2 | P2 | 9r9r |
| QUIN19C2-M8 | M | QUIN19C2 | 2019 | 2 | P2 | 9r9r |
| QUIN19C2-W1 | W | QUIN19C2 | 2019 | 2 | P2P2 | 9a9r |
| QUIN19C2-W2 | W | QUIN19C2 | 2019 | 2 | P1P2 | 9a9r |
| QUIN19C2-W3 | W | QUIN19C2 | 2019 | 2 | P1P1 | 9a9r |
| QUIN19C2-W3r | W | QUIN19C2 | 2019 | 2 | P1P1 | 9a9r |
| QUIN19C2-W4 | W | QUIN19C2 | 2019 | 2 | MAP1 | 9a9a |
| QUIN19C2-W5 | W | QUIN19C2 | 2019 | 2 | MAP1 | 9a9r |

| | | | | | | |
|---|---|---|---|---|---|---|
| QUIN19C2-W5r | W | QUIN19C2 | 2019 | 2 | P1P2 | 9a9a |
| QUIN19C3-G1 | G | QUIN19C3 | 2019 | 2 | P1P2 | 9a9r |
| QUIN19C3-G2 | G | QUIN19C3 | 2019 | 2 | P1P2 | 9a9r |
| QUIN19C3-G5 | G | QUIN19C3 | 2019 | 2 | P2P2 | 9r9r |
| QUIN19C3-G6 | G | QUIN19C3 | 2019 | 2 | P1P2 | 9a9r |
| QUIN19C3-G7 | G | QUIN19C3 | 2019 | 2 | P1P2 | 9a9r |
| QUIN19C3-Q1 | Q | QUIN19C3 | 2019 | 2 | P1P2 | 9a9r |
| QUIN19C3-Q2 | Q | QUIN19C3 | 2019 | 2 | P1P2 | 9a9r |
| QUIN19C3-Q3 | Q | QUIN19C3 | 2019 | 2 | P1P2 | 9a9r |
| QUIN19C3-W1 | W | QUIN19C3 | 2019 | 2 | MDP1 | 9a9a |
| QUIN19C3-W2 | W | QUIN19C3 | 2019 | 2 | MAP1 | 9a9a |
| QUIN19C3-W3 | W | QUIN19C3 | 2019 | 2 | MAP1 | 9a9a |
| QUIN19C4-G1 | G | QUIN19C4 | 2019 | 2 | P2P2 | 9r9r |
| QUIN19C4-M2 | M | QUIN19C4 | 2019 | 2 | P2 | 9r9r |
| QUIN19C4-M3 | M | QUIN19C4 | 2019 | 2 | P2 | 9r9r |
| QUIN19C4-M4 | M | QUIN19C4 | 2019 | 2 | P2 | 9r9r |
| QUIN19C4-M5 | M | QUIN19C4 | 2019 | 2 | P2 | 9r9r |
| QUIN19C4-M6 | M | QUIN19C4 | 2019 | 2 | P2 | 9r9r |
| QUIN19C4-M7 | M | QUIN19C4 | 2019 | 2 | P2 | 9r9r |
| QUIN19C4-M8 | M | QUIN19C4 | 2019 | 2 | P2 | 9r9r |
| QUIN19C4-W1 | W | QUIN19C4 | 2019 | 2 | P1P2 | 9a9r |
| QUIN19C4-W2 | W | QUIN19C4 | 2019 | 2 | P1P2 | 9a9r |
| QUIN19C4-W3 | W | QUIN19C4 | 2019 | 2 | MAP2 | 9a9r |
| QUIN19C4-W4 | W | QUIN19C4 | 2019 | 2 | P2P2 | 9a9r |
| QUIN19C4-W5 | W | QUIN19C4 | 2019 | 2 | P1P2 | 9r9r |
| QUIN19C5-W1 | W | QUIN19C5 | 2019 | 2 | P2P2 | 9a9r |
| QUIN19C5-W2 | W | QUIN19C5 | 2019 | 2 | MAP2 | 9a9r |
| QUIN19C5-W3 | W | QUIN19C5 | 2019 | 2 | P2P2 | 9r9r |
| QUIN19C5-W4 | W | QUIN19C5 | 2019 | 2 | P1P2 | 9a9r |
| QUIN19C5-W5 | W | QUIN19C5 | 2019 | 2 | P2P2 | 9a9r |
| QUIN19C6-M1 | M | QUIN19C6 | 2019 | 2 | P2 | 9r9r |
| QUIN19C6-M2 | M | QUIN19C6 | 2019 | 2 | P2 | 9r9r |
| QUIN19C6-M3 | M | QUIN19C6 | 2019 | 2 | P2 | 9r9r |
| QUIN19C6-M5 | M | QUIN19C6 | 2019 | 2 | P2 | 9r9r |
| QUIN19C6-M7 | M | QUIN19C6 | 2019 | 2 | P2 | 9r9r |
| QUIN19C6-M8 | M | QUIN19C6 | 2019 | 2 | P2 | 9r9r |
| QUIN19C7-W1 | W | QUIN19C7 | 2019 | 2 | P2P2 | 9a9r |
| QUIN19C7-W2 | W | QUIN19C7 | 2019 | 2 | P1P2 | 9a9r |
| QUIN19C7-W3 | W | QUIN19C7 | 2019 | 2 | P2P2 | 9a9r |
| QUIN19C7-W4 | W | QUIN19C7 | 2019 | 2 | P1P2 | 9a9r |
| QUIN19C7-W5 | W | QUIN19C7 | 2019 | 2 | MAP2 | 9a9a |
| QUIN19M1-W1 | W | QUIN19M1 | 2019 | 2 | MAMA | 9a9a |
| QUIN19M1-W2 | W | QUIN19M1 | 2019 | 2 | MAMA | 9a9a |
| QUIN19M1-W3 | W | QUIN19M1 | 2019 | 2 | MAMA | 9a9a |

| | | | | | | |
|---|---|---|---|---|---|---|
| QUIN19M1-W4 | W | QUIN19M1 | 2019 | 2 | MAMA | 9a9a |
| QUIN19M1-W5 | W | QUIN19M1 | 2019 | 2 | MAMA | 9a9a |
| QUIN2001-G1 | G | QUIN2001 | 2020 | 2 | MAMD | 9a9a |
| QUIN2001-W2 | W | QUIN2001 | 2020 | 2 | MAMD | 9a9a |
| QUIN2001-W3 | W | QUIN2001 | 2020 | 2 | MAMD | 9a9a |
| QUIN2001-W4 | W | QUIN2001 | 2020 | 2 | MAMA | 9a9a |
| QUIN2001-W5 | W | QUIN2001 | 2020 | 2 | MAMD | 9a9a |
| QUIN2002-G1 | G | QUIN2002 | 2020 | 2 | MAMA | 9a9a |
| QUIN2002-G2 | G | QUIN2002 | 2020 | 2 | MAMA | 9a9a |
| QUIN2002-M1 | M | QUIN2002 | 2020 | 2 | MA | 9a9a |
| QUIN2002-M2 | M | QUIN2002 | 2020 | 2 | MA | 9a9a |
| QUIN2002-M3 | M | QUIN2002 | 2020 | 2 | MAMA | 9a9a |
| QUIN2002-M4 | M | QUIN2002 | 2020 | 2 | MA | 9a9a |
| QUIN2002-M5 | M | QUIN2002 | 2020 | 2 | MA | 9a9a |
| QUIN2002-M6 | M | QUIN2002 | 2020 | 2 | MA | 9a9a |
| QUIN2002-M7 | M | QUIN2002 | 2020 | 2 | MA | 9a9a |
| QUIN2002-W5 | W | QUIN2002 | 2020 | 2 | MAMA | 9a9a |
| QUIN2003-M1 | M | QUIN2003 | 2020 | 2 | P2 | 9r9r |
| QUIN2003-M2 | M | QUIN2003 | 2020 | 2 | P2 | 9r9r |
| QUIN2003-M3 | M | QUIN2003 | 2020 | 2 | P2 | 9r9r |
| QUIN2003-M4 | M | QUIN2003 | 2020 | 2 | P2 | 9r9r |
| QUIN2003-M5 | M | QUIN2003 | 2020 | 2 | P2 | 9r9r |
| QUIN2003-M6 | M | QUIN2003 | 2020 | 2 | P2 | 9r9r |
| QUIN2003-M7 | M | QUIN2003 | 2020 | 2 | P2 | 9r9r |
| QUIN2003-M8 | M | QUIN2003 | 2020 | 2 | P2 | 9r9r |
| QUIN2003-W1 | W | QUIN2003 | 2020 | 2 | P1P2 | 9r9r |
| QUIN2003-W2 | W | QUIN2003 | 2020 | 2 | P1P2 | 9a9r |
| QUIN2004-M1 | M | QUIN2004 | 2020 | 2 | P2 | 9r9r |
| QUIN2004-M2 | M | QUIN2004 | 2020 | 2 | P2 | 9r9r |
| QUIN2004-M3 | M | QUIN2004 | 2020 | 2 | P2 | 9r9r |
| QUIN2004-M4 | M | QUIN2004 | 2020 | 2 | P2 | 9r9r |
| QUIN2004-M5 | M | QUIN2004 | 2020 | 2 | P2 | 9r9r |
| QUIN2004-M6 | M | QUIN2004 | 2020 | 2 | P2 | 9r9r |
| QUIN2004-M7 | M | QUIN2004 | 2020 | 2 | P2 | 9r9r |
| QUIN2004-W1 | W | QUIN2004 | 2020 | 2 | P2P2 | 9a9r |
| QUIN2004-W2 | W | QUIN2004 | 2020 | 2 | P2P2 | 9a9r |
| QUIN2004-W3 | W | QUIN2004 | 2020 | 2 | P2P2 | 9r9r |
| QUIN2004-W5 | W | QUIN2004 | 2020 | 2 | P2P2 | 9a9r |
| QUIN2005-M1 | M | QUIN2005 | 2020 | 2 | P2 | 9r9r |
| QUIN2005-W1 | W | QUIN2005 | 2020 | 2 | P1P2 | 9a9r |
| QUIN2005-W2 | W | QUIN2005 | 2020 | 2 | P2P2 | 9a9r |
| QUIN2005-W3 | W | QUIN2005 | 2020 | 2 | P1P2 | 9a9r |
| QUIN2005-W4 | W | QUIN2005 | 2020 | 2 | MAP2 | 9a9r |
| QUIN2006-M1 | M | QUIN2006 | 2020 | 2 | MA | 9a9a |

| | | | | | | |
|---|---|---|---|---|---|---|
| QUIN2006-M2 | M | QUIN2006 | 2020 | 2 | MA | 9a9a |
| QUIN2006-M3 | M | QUIN2006 | 2020 | 2 | MA | 9a9a |
| QUIN2006-M4 | M | QUIN2006 | 2020 | 2 | MA | 9a9a |
| QUIN2006-M5 | M | QUIN2006 | 2020 | 2 | MA | 9a9a |
| QUIN2006-M6 | M | QUIN2006 | 2020 | 2 | MA | 9a9a |
| QUIN2006-M7 | M | QUIN2006 | 2020 | 2 | MA | 9a9a |
| QUIN2006-M8 | M | QUIN2006 | 2020 | 2 | MA | 9a9a |
| QUIN2006-W1 | W | QUIN2006 | 2020 | 2 | MAMA | 9a9a |
| QUIN2006-W2 | W | QUIN2006 | 2020 | 2 | MAMA | 9a9a |
| QUIN2006-W3 | W | QUIN2006 | 2020 | 2 | MAMA | 9a9a |
| QUIN2006-W4 | W | QUIN2006 | 2020 | 2 | MAMA | 9a9a |
| QUIN2006-W5 | W | QUIN2006 | 2020 | 2 | MAMA | 9a9a |
| QUIN2007-G1 | G | QUIN2007 | 2020 | 2 | P2P2 | 9r9r |
| QUIN2007-M1 | M | QUIN2007 | 2020 | 2 | P2 | 9r9r |
| QUIN2007-M2 | M | QUIN2007 | 2020 | 2 | P2 | 9r9r |
| QUIN2007-M3 | M | QUIN2007 | 2020 | 2 | P2 | 9r9r |
| QUIN2007-M4 | M | QUIN2007 | 2020 | 2 | P2 | 9r9r |
| QUIN2007-M5 | M | QUIN2007 | 2020 | 2 | P2 | 9r9r |
| QUIN2007-M6 | M | QUIN2007 | 2020 | 2 | P2 | 9r9r |
| QUIN2007-M7 | M | QUIN2007 | 2020 | 2 | P2 | 9r9r |
| QUIN2007-M9 | M | QUIN2007 | 2020 | 2 | MA | 9a9a |
| QUIN2007-W1 | W | QUIN2007 | 2020 | 2 | P1P2 | unkn |
| QUIN2007-W2 | W | QUIN2007 | 2020 | 2 | P2P2 | 9a9r |
| QUIN2007-W3 | W | QUIN2007 | 2020 | 2 | P1P2 | 9a9r |
| QUIN2007-W4 | W | QUIN2007 | 2020 | 2 | MAP2 | unkn |
| QUIN2007-W5 | W | QUIN2007 | 2020 | 2 | P1P2 | 9a9r |
| QUIN2008-M1 | M | QUIN2008 | 2020 | 2 | P2 | 9r9r |
| QUIN2008-M2 | M | QUIN2008 | 2020 | 2 | P2 | 9r9r |
| QUIN2008-M3 | M | QUIN2008 | 2020 | 2 | P2 | 9r9r |
| QUIN2008-M4 | M | QUIN2008 | 2020 | 2 | P2 | 9r9r |
| QUIN2008-M5 | M | QUIN2008 | 2020 | 2 | P2 | 9r9r |
| QUIN2008-M6 | M | QUIN2008 | 2020 | 2 | P2 | 9r9r |
| QUIN2008-M7 | M | QUIN2008 | 2020 | 2 | P2 | 9r9r |
| QUIN2008-M8 | M | QUIN2008 | 2020 | 2 | P2 | 9r9r |
| QUIN2008-W1 | W | QUIN2008 | 2020 | 2 | P2P2 | 9r9r |
| QUIN2008-W2 | W | QUIN2008 | 2020 | 2 | P1P2 | 9a9r |
| QUIN2008-W3 | W | QUIN2008 | 2020 | 2 | P2P2 | 9r9r |
| QUIN2008-W4 | W | QUIN2008 | 2020 | 2 | MAP2 | 9a9r |
| QUIN2008-W5 | W | QUIN2008 | 2020 | 2 | MAP1 | 9a9a |
| QUIN2101-G1 | G | QUIN2101 | 2021 | 4 | P2P2 | 9r9r |
| QUIN2101-G2 | G | QUIN2101 | 2021 | 4 | P2P2 | 9r9r |
| QUIN2101-M1 | M | QUIN2101 | 2021 | 4 | P2 | 9r9r |
| QUIN2101-M2 | M | QUIN2101 | 2021 | 4 | P2 | 9r9r |
| QUIN2101-M3 | M | QUIN2101 | 2021 | 4 | P2 | 9r9r |

| | | | | | | |
|---|---|---|---|---|---|---|
| QUIN2101-M4 | M | QUIN2101 | 2021 | 4 | P2 | 9r9r |
| QUIN2101-M5 | M | QUIN2101 | 2021 | 4 | P2 | 9r9r |
| QUIN2101-M6 | M | QUIN2101 | 2021 | 4 | P2 | 9r9r |
| QUIN2101-M7 | M | QUIN2101 | 2021 | 4 | P2 | 9r9r |
| QUIN2101-M8 | M | QUIN2101 | 2021 | 4 | P2 | 9r9r |
| QUIN2101-W1 | W | QUIN2101 | 2021 | 3 | P2P2 | 9a9r |
| QUIN2101-W2 | W | QUIN2101 | 2021 | 3 | P2P2 | 9r9r |
| QUIN2101-W3 | W | QUIN2101 | 2021 | 3 | P2P2 | 9a9r |
| QUIN2101-W4 | W | QUIN2101 | 2021 | 3 | P1P2 | 9a9r |
| QUIN2101-W5 | W | QUIN2101 | 2021 | 3 | MAP2 | 9a9r |
| QUIN2102-M1 | M | QUIN2102 | 2021 | 4 | P2 | 9r9r |
| QUIN2102-M2 | M | QUIN2102 | 2021 | 4 | P2 | 9r9r |
| QUIN2102-M3 | M | QUIN2102 | 2021 | 4 | P2 | 9r9r |
| QUIN2102-M4 | M | QUIN2102 | 2021 | 4 | P2 | 9r9r |
| QUIN2102-W1 | W | QUIN2102 | 2021 | 3 | P2P2 | 9r9r |
| QUIN2102-W2 | W | QUIN2102 | 2021 | 3 | MAP2 | 9a9a |
| QUIN2102-W3 | W | QUIN2102 | 2021 | 3 | MAP2 | 9a9a |
| QUIN2102-W4 | W | QUIN2102 | 2021 | 3 | P1P2 | 9a9r |
| QUIN2102-W5 | W | QUIN2102 | 2021 | 3 | MAP2 | 9a9r |
| QUIN2103-M1 | M | QUIN2103 | 2021 | 4 | MA | 9a9a |
| QUIN2103-M3 | M | QUIN2103 | 2021 | 4 | MA | 9a9a |
| QUIN2103-M4 | M | QUIN2103 | 2021 | 4 | unk | 9a9a |
| QUIN2103-M5 | M | QUIN2103 | 2021 | 4 | MA | 9a9a |
| QUIN2103-W1 | W | QUIN2103 | 2021 | 3 | MAMA | 9a9a |
| QUIN2103-W2 | W | QUIN2103 | 2021 | 3 | MAMA | 9a9a |
| QUIN2103-W3 | W | QUIN2103 | 2021 | 3 | MAMA | 9a9a |
| QUIN2103-W4 | W | QUIN2103 | 2021 | 3 | MAMA | 9a9a |
| QUIN2103-W5 | W | QUIN2103 | 2021 | 3 | MAMA | 9a9a |
| QUIN2104-M2 | M | QUIN2104 | 2021 | 4 | MA | 9a9a |
| QUIN2104-M3 | M | QUIN2104 | 2021 | 4 | MA | 9a9a |
| QUIN2104-M4 | M | QUIN2104 | 2021 | 4 | MA | 9a9a |
| QUIN2104-W1 | W | QUIN2104 | 2021 | 3 | MAMA | 9a9a |
| QUIN2104-W2 | W | QUIN2104 | 2021 | 3 | MAMA | 9a9a |
| QUIN2104-W3 | W | QUIN2104 | 2021 | 3 | MAMA | 9a9a |
| QUIN2104-W4 | W | QUIN2104 | 2021 | 3 | MAMA | 9a9a |
| QUIN2104-W5 | W | QUIN2104 | 2021 | 3 | MAMA | 9a9a |
| QUIN2105-M1 | M | QUIN2105 | 2021 | 4 | P1 | 9a9a |
| QUIN2105-M2 | M | QUIN2105 | 2021 | 4 | P1 | 9a9a |
| QUIN2105-W1 | W | QUIN2105 | 2021 | 3 | MAP1 | 9a9a |
| QUIN2105-W2 | W | QUIN2105 | 2021 | 3 | MAP1 | 9a9a |
| QUIN2105-W3 | W | QUIN2105 | 2021 | 3 | MAP1 | 9a9a |
| QUIN2105-W4 | W | QUIN2105 | 2021 | 3 | P1P1 | 9a9a |
| QUIN2105-W5 | W | QUIN2105 | 2021 | 3 | MAP1 | 9a9a |
| QUIN2106-M2 | M | QUIN2106 | 2021 | 4 | P1 | 9a9a |

| | | | | | | |
|---|---|---|---|---|---|---|
| QUIN2106-M3 | M | QUIN2106 | 2021 | 4 | P1 | 9a9a |
| QUIN2106-M4 | M | QUIN2106 | 2021 | 4 | P1 | 9a9a |
| QUIN2106-M5 | M | QUIN2106 | 2021 | 4 | P1 | 9a9a |
| QUIN2106-M6 | M | QUIN2106 | 2021 | 4 | P1 | 9a9a |
| QUIN2106-W1 | W | QUIN2106 | 2021 | 3 | P1P1 | 9a9a |
| QUIN2106-W2 | W | QUIN2106 | 2021 | 3 | MAP1 | 9a9a |
| QUIN2106-W3 | W | QUIN2106 | 2021 | 4 | MAP1 | 9a9a |
| QUIN2106-W3r | W | QUIN2106 | 2021 | 3 | MAP1 | 9a9a |
| QUIN2106-W4 | W | QUIN2106 | 2021 | 3 | MAP1 | 9a9a |
| QUIN2106-W5 | W | QUIN2106 | 2021 | 3 | MAP1 | 9a9a |
| QUINI2001-G1 | G | QUINI2001 | 2020 | 2 | MAMD | 9a9a |
| QUINI2001-G2 | G | QUINI2001 | 2020 | 2 | MAMD | 9a9a |
| QUINI2001-G3 | G | QUINI2001 | 2020 | 2 | MAMA | 9a9a |
| QUINI2001-G4 | G | QUINI2001 | 2020 | 2 | MAMA | 9a9a |
| QUINI2001-G5 | G | QUINI2001 | 2020 | 2 | MAMD | 9a9a |
| QUINI2001-G6 | G | QUINI2001 | 2020 | 2 | MAMD | 9a9a |
| QUINI2001-G7 | G | QUINI2001 | 2020 | 2 | MAMA | 9a9a |
| QUINI2001-G8 | G | QUINI2001 | 2020 | 2 | MAMD | 9a9a |
| QUINI2001-W1 | W | QUINI2001 | 2020 | 2 | MAMA | 9a9a |
| QUINI2001-W2 | W | QUINI2001 | 2020 | 2 | MAMD | 9a9a |
| QUINI2001-W4 | W | QUINI2001 | 2020 | 2 | MAMD | 9a9a |
| QUINI2001-W5 | W | QUINI2001 | 2020 | 2 | MAMD | 9a9a |
| QUINI2002-M1 | M | QUINI2002 | 2020 | 2 | MA | 9a9a |
| QUINI2002-M2 | M | QUINI2002 | 2020 | 2 | MA | 9a9a |
| QUINI2002-M3 | M | QUINI2002 | 2020 | 2 | MA | 9a9a |
| QUINI2002-M4 | M | QUINI2002 | 2020 | 2 | MA | 9a9a |
| QUINI2002-W1 | W | QUINI2002 | 2020 | 2 | MAMD | 9a9a |
| QUINI2002-W2 | W | QUINI2002 | 2020 | 2 | MAMA | 9a9a |
| QUINI2002-W3 | W | QUINI2002 | 2020 | 2 | MAMA | 9a9a |
| QUINI2002-W4 | W | QUINI2002 | 2020 | 2 | MAMD | 9a9a |
| QUINI2002-W5 | W | QUINI2002 | 2020 | 2 | MAMA | 9a9a |
| QUINI2003-M1 | M | QUINI2003 | 2020 | 2 | MA | 9a9a |
| QUINI2003-M3 | M | QUINI2003 | 2020 | 2 | MA | 9a9a |
| QUINI2003-M4 | M | QUINI2003 | 2020 | 2 | MA | 9a9a |
| QUINI2003-W1 | W | QUINI2003 | 2020 | 2 | MAMD | 9a9a |
| QUINI2003-W2 | W | QUINI2003 | 2020 | 2 | MAMD | 9a9a |
| QUINI2003-W3 | W | QUINI2003 | 2020 | 2 | MAMD | 9a9a |
| QUINI2003-W4 | W | QUINI2003 | 2020 | 2 | MAMD | 9a9a |
| QUINI2003-W5 | W | QUINI2003 | 2020 | 2 | MAMA | 9a9a |
| QUINI2004-M1 | M | QUINI2004 | 2020 | 2 | MA | 9a9a |
| QUINI2004-M2 | M | QUINI2004 | 2020 | 2 | MA | 9a9a |
| QUINI2004-M3 | M | QUINI2004 | 2020 | 2 | MA | 9a9a |
| QUINI2004-M4 | M | QUINI2004 | 2020 | 2 | P2 | 9r9r |
| QUINI2004-M5 | M | QUINI2004 | 2020 | 2 | MA | 9a9a |

| | | | | | | |
|---|---|---|---|---|---|---|
| QUINI2004-M6 | M | QUINI2004 | 2020 | 2 | MA | 9a9a |
| QUINI2004-M7 | M | QUINI2004 | 2020 | 2 | MA | 9a9a |
| QUINI2004-W1 | W | QUINI2004 | 2020 | 2 | MAMD | 9a9a |
| QUINI2004-W2 | W | QUINI2004 | 2020 | 2 | MAMD | 9a9a |
| QUINI2004-W3 | W | QUINI2004 | 2020 | 2 | MAMD | 9a9a |
| QUINI2004-W4 | W | QUINI2004 | 2020 | 2 | MAMD | 9a9a |
| QUINI2004-W5 | W | QUINI2004 | 2020 | 2 | MAMD | 9a9a |
| QUINI2005-M2 | M | QUINI2005 | 2020 | 2 | P1 | 9a9a |
| QUINI2005-W1 | W | QUINI2005 | 2020 | 2 | MAP1 | 9a9a |
| QUINI2005-W2 | W | QUINI2005 | 2020 | 2 | MAP1 | 9a9a |
| QUINI2005-W3 | W | QUINI2005 | 2020 | 2 | MAP1 | 9a9a |
| QUINI2005-W4 | W | QUINI2005 | 2020 | 2 | MAP1 | 9a9a |
| QUINI2005-W5 | W | QUINI2005 | 2020 | 2 | MAP1 | 9a9a |
| QUINI2006-G1 | G | QUINI2006 | 2020 | 2 | P1P1 | 9a9a |
| QUINI2006-G10 | G | QUINI2006 | 2020 | 4 | P1P1 | 9a9a |
| QUINI2006-G2 | G | QUINI2006 | 2020 | 2 | MAP1 | 9a9a |
| QUINI2006-G3 | G | QUINI2006 | 2020 | 2 | P1P1 | 9a9a |
| QUINI2006-G4 | G | QUINI2006 | 2020 | 2 | P1P1 | 9a9a |
| QUINI2006-G5 | G | QUINI2006 | 2020 | 2 | P1P1 | 9a9a |
| QUINI2006-G6 | G | QUINI2006 | 2020 | 2 | P1P1 | 9a9a |
| QUINI2006-G7 | G | QUINI2006 | 2020 | 2 | P1P1 | 9a9a |
| QUINI2006-G8 | G | QUINI2006 | 2020 | 2 | P1P1 | 9a9a |
| QUINI2006-G9 | G | QUINI2006 | 2020 | 4 | P1P1 | 9a9a |
| QUINI2006-M1 | M | QUINI2006 | 2020 | 2 | P1 | 9a9a |
| QUINI2006-M2 | M | QUINI2006 | 2020 | 2 | P1 | 9a9a |
| QUINI2006-M3 | M | QUINI2006 | 2020 | 2 | P1 | 9a9a |
| QUINI2006-M4 | M | QUINI2006 | 2020 | 2 | P1 | 9a9a |
| QUINI2006-M5 | M | QUINI2006 | 2020 | 2 | P1 | 9a9a |
| QUINI2006-M6 | M | QUINI2006 | 2020 | 2 | P1 | 9a9a |
| QUINI2006-M7 | M | QUINI2006 | 2020 | 2 | P1 | 9a9a |
| QUINI2006-M8 | M | QUINI2006 | 2020 | 2 | P1 | 9a9a |
| QUINI2006-W1 | W | QUINI2006 | 2020 | 2 | P1P1 | 9a9a |
| QUINI2006-W2 | W | QUINI2006 | 2020 | 2 | MAP1 | 9a9a |
| QUINI2006-W3 | W | QUINI2006 | 2020 | 2 | MAP1 | 9a9a |
| QUINI2006-W4 | W | QUINI2006 | 2020 | 2 | MAP1 | 9a9a |
| QUINI2006-W5 | W | QUINI2006 | 2020 | 2 | MAP1 | 9a9a |
| QUINI2007-G1 | G | QUINI2007 | 2020 | 2 | P1P1 | 9a9a |
| QUINI2007-G2 | G | QUINI2007 | 2020 | 2 | P1P1 | 9a9a |
| QUINI2007-G3 | G | QUINI2007 | 2020 | 2 | P1P1 | 9a9a |
| QUINI2007-G4 | G | QUINI2007 | 2020 | 2 | MAP1 | 9a9a |
| QUINI2007-M1 | M | QUINI2007 | 2020 | 2 | P1 | 9a9a |
| QUINI2007-M2 | M | QUINI2007 | 2020 | 2 | P1 | 9a9a |
| QUINI2007-M3 | M | QUINI2007 | 2020 | 2 | P1 | 9a9a |
| QUINI2007-M4 | M | QUINI2007 | 2020 | 2 | P1 | 9a9a |

| | | | | | | |
|---|---|---|---|---|---|---|
| QUINI2007-M5 | M | QUINI2007 | 2020 | 2 | P1 | 9a9a |
| QUINI2007-M6 | M | QUINI2007 | 2020 | 2 | P1 | 9a9a |
| QUINI2007-M7 | M | QUINI2007 | 2020 | 2 | P1 | 9a9a |
| QUINI2007-M8 | M | QUINI2007 | 2020 | 2 | P1 | 9a9a |
| QUINI2007-W1 | W | QUINI2007 | 2020 | 2 | MAP1 | 9a9a |
| QUINI2007-W2 | W | QUINI2007 | 2020 | 2 | MAP1 | 9a9a |
| QUINI2007-W3 | W | QUINI2007 | 2020 | 2 | MAP1 | 9a9a |
| QUINI2007-W4 | W | QUINI2007 | 2020 | 2 | MAP1 | 9a9a |
| QUINI2007-W5 | W | QUINI2007 | 2020 | 2 | MAP1 | 9a9a |
| QUINI2008-G1 | G | QUINI2008 | 2020 | 2 | P1P1 | 9a9a |
| QUINI2008-G2 | G | QUINI2008 | 2020 | 2 | P1P1 | 9a9a |
| QUINI2008-G3 | G | QUINI2008 | 2020 | 2 | MAP1 | 9a9a |
| QUINI2008-G4 | G | QUINI2008 | 2020 | 2 | P1P1 | 9a9a |
| QUINI2008-G5 | G | QUINI2008 | 2020 | 2 | P1P1 | 9a9a |
| QUINI2008-W1 | W | QUINI2008 | 2020 | 2 | MAP1 | 9a9a |
| QUINI2008-W2 | W | QUINI2008 | 2020 | 2 | MAP1 | 9a9a |
| QUINI2008-W3 | W | QUINI2008 | 2020 | 2 | MAP1 | 9a9a |
| QUINI2008-W4 | W | QUINI2008 | 2020 | 2 | MAP1 | 9a9a |
| QUINI2008-W5 | W | QUINI2008 | 2020 | 2 | P1P1 | 9a9a |
| QUINI2009-G1 | G | QUINI2009 | 2020 | 2 | P1P1 | 9a9a |
| QUINI2009-M1 | M | QUINI2009 | 2020 | 2 | P1 | 9a9a |
| QUINI2009-M2 | M | QUINI2009 | 2020 | 2 | P1 | 9a9a |
| QUINI2009-M3 | M | QUINI2009 | 2020 | 2 | P1 | 9a9a |
| QUINI2009-W1 | W | QUINI2009 | 2020 | 2 | MAP1 | 9a9a |
| QUINI2009-W2 | W | QUINI2009 | 2020 | 2 | MAP1 | 9a9a |
| QUINI2009-W3 | W | QUINI2009 | 2020 | 2 | MAP1 | 9a9a |
| QUINI2009-W4 | W | QUINI2009 | 2020 | 2 | MAP1 | 9a9a |
| QUINI2009-W5 | W | QUINI2009 | 2020 | 2 | MAP1 | 9a9a |
| QUINI2010-W1 | W | QUINI2010 | 2020 | 2 | MAP1 | 9a9a |
| QUINI2010-W2 | W | QUINI2010 | 2020 | 2 | MAP1 | 9a9a |
| QUINI2010-W3 | W | QUINI2010 | 2020 | 2 | MAP1 | 9a9a |
| QUINI2010-W4 | W | QUINI2010 | 2020 | 2 | MAP1 | 9a9a |
| QUINI2010-W5 | W | QUINI2010 | 2020 | 2 | MAP1 | 9a9a |
| QUINI2011-G1 | G | QUINI2011 | 2020 | 2 | P1P1 | 9a9a |
| QUINI2011-M1 | M | QUINI2011 | 2020 | 2 | P1 | 9a9a |
| QUINI2011-W1 | W | QUINI2011 | 2020 | 2 | MAP1 | 9a9a |
| QUINI2012-G1 | G | QUINI2012 | 2020 | 2 | MAP2 | 9a9r |
| QUINI2012-G2 | G | QUINI2012 | 2020 | 2 | P1P2 | 9a9r |
| QUINI2012-G3 | G | QUINI2012 | 2020 | 2 | P2P2 | 9r9r |
| QUINI2012-M2 | M | QUINI2012 | 2020 | 2 | P2 | 9r9r |
| QUINI2012-W1 | W | QUINI2012 | 2020 | 2 | MAP1 | 9a9a |
| QUINI2012-W3 | W | QUINI2012 | 2020 | 2 | P1P1 | 9a9a |
| QUINI2012-W5 | W | QUINI2012 | 2020 | 2 | MAP1 | 9a9a |
| QUINI2101-M1 | M | QUINI2101 | 2021 | 4 | P1 | 9a9a |

| | | | | | | |
|---|---|---|---|---|---|---|
| QUINI2101-M2 | M | QUINI2101 | 2021 | 4 | P1 | 9a9a |
| QUINI2101-M3 | M | QUINI2101 | 2021 | 4 | P1 | 9a9a |
| QUINI2101-M4 | M | QUINI2101 | 2021 | 4 | P1 | 9a9a |
| QUINI2101-M5 | M | QUINI2101 | 2021 | 4 | MA | 9a9a |
| QUINI2101-M6 | M | QUINI2101 | 2021 | 4 | P1 | 9a9a |
| QUINI2101-M7 | M | QUINI2101 | 2021 | 4 | P1 | 9a9a |
| QUINI2101-M8 | M | QUINI2101 | 2021 | 4 | P1 | 9a9a |
| QUINI2101-W1 | W | QUINI2101 | 2021 | 3 | MAP1 | 9a9a |
| QUINI2101-W2 | W | QUINI2101 | 2021 | 3 | MAP1 | 9a9a |
| QUINI2101-W3 | W | QUINI2101 | 2021 | 3 | MAP1 | 9a9a |
| QUINI2101-W4 | W | QUINI2101 | 2021 | 3 | MAP1 | 9a9a |
| QUINI2101-W5 | W | QUINI2101 | 2021 | 3 | MAP1 | 9a9a |
| QUINI2102-W1 | W | QUINI2102 | 2021 | 3 | MAP2 | 9a9a |
| QUINI2102-W2 | W | QUINI2102 | 2021 | 3 | P1P2 | 9a9r |
| QUINI2102-W3 | W | QUINI2102 | 2021 | 3 | P1P2 | 9a9r |
| QUINI2102-W4 | W | QUINI2102 | 2021 | 3 | MAP2 | 9a9r |
| QUINI2102-W5 | W | QUINI2102 | 2021 | 3 | MAP2 | 9a9r |
| QUINI2103-M1 | M | QUINI2103 | 2021 | 4 | MA | 9a9a |
| QUINI2103-M2 | M | QUINI2103 | 2021 | 4 | MA | 9a9a |
| QUINI2103-M3 | M | QUINI2103 | 2021 | 4 | MA | 9a9a |
| QUINI2103-M4 | M | QUINI2103 | 2021 | 4 | MA | 9a9a |
| QUINI2103-M5 | M | QUINI2103 | 2021 | 4 | MA | 9a9a |
| QUINI2103-M6 | M | QUINI2103 | 2021 | 4 | MA | 9a9a |
| QUINI2103-M7 | M | QUINI2103 | 2021 | 4 | MA | 9a9a |
| QUINI2103-W1 | W | QUINI2103 | 2021 | 3 | MAMA | 9a9a |
| QUINI2103-W2 | W | QUINI2103 | 2021 | 3 | MAMA | 9a9a |
| QUINI2103-W3 | W | QUINI2103 | 2021 | 3 | MAMA | 9a9a |
| QUINI2103-W4 | W | QUINI2103 | 2021 | 3 | MAMA | 9a9a |
| QUINI2103-W5 | W | QUINI2103 | 2021 | 3 | MAMA | 9a9a |
| QUINI2104-G1 | G | QUINI2104 | 2021 | 4 | MAMA | 9a9a |
| QUINI2104-W1 | W | QUINI2104 | 2021 | 3 | MAMA | 9a9a |
| QUINI2104-W2 | W | QUINI2104 | 2021 | 3 | MAMA | 9a9a |
| QUINI2104-W3 | W | QUINI2104 | 2021 | 3 | MAMA | 9a9a |
| QUINI2104-W4 | W | QUINI2104 | 2021 | 3 | MAMA | 9a9a |
| QUINI2104-W5 | W | QUINI2104 | 2021 | 3 | MAMA | 9a9a |
| QUINI2105-M1 | M | QUINI2105 | 2021 | 4 | P1 | 9a9a |
| QUINI2105-M3 | M | QUINI2105 | 2021 | 4 | P1 | 9a9a |
| QUINI2105-M4 | M | QUINI2105 | 2021 | 4 | MA | 9a9a |
| QUINI2105-M5 | M | QUINI2105 | 2021 | 4 | MA | 9a9a |
| QUINI2105-W1 | W | QUINI2105 | 2021 | 3 | MAP1 | 9a9a |
| QUINI2105-W2 | W | QUINI2105 | 2021 | 3 | MAP1 | 9a9a |
| QUINI2105-W3 | W | QUINI2105 | 2021 | 3 | MAP1 | 9a9a |
| QUINI2105-W4 | W | QUINI2105 | 2021 | 3 | MAP1 | 9a9a |
| QUINI2105-W5 | W | QUINI2105 | 2021 | 3 | MAP1 | 9a9a |

| | | | | | | |
|---|---|---|---|---|---|---|
| QUINI2106-M1 | M | QUINI2106 | 2021 | 4 | MA | 9a9a |
| QUINI2106-M2 | M | QUINI2106 | 2021 | 4 | MD | 9a9a |
| QUINI2106-M3 | M | QUINI2106 | 2021 | 4 | MA | 9a9a |
| QUINI2106-M4 | M | QUINI2106 | 2021 | 4 | MA | 9a9a |
| QUINI2106-M5 | M | QUINI2106 | 2021 | 4 | MD | 9a9a |
| QUINI2106-M6 | M | QUINI2106 | 2021 | 4 | MD | 9a9a |
| QUINI2106-W1 | W | QUINI2106 | 2021 | 3 | MAMD | 9a9a |
| QUINI2107-G1 | G | QUINI2107 | 2021 | 4 | MAMA | 9a9a |
| QUINI2108-G1 | G | QUINI2108 | 2021 | 4 | MAMD | 9a9a |
| QUINI2108-G2 | G | QUINI2108 | 2021 | 4 | MAMD | 9a9a |
| QUINI2108-G3 | G | QUINI2108 | 2021 | 4 | MAMD | 9a9a |
| QUINI2108-G4 | G | QUINI2108 | 2021 | 4 | MAMA | 9a9a |
| QUINI2108-W1 | W | QUINI2108 | 2021 | 3 | MAMA | 9a9a |
| QUINI2108-W2 | W | QUINI2108 | 2021 | 3 | MAMD | 9a9a |
| QUINI2108-W3 | W | QUINI2108 | 2021 | 3 | MAMD | 9a9a |
| QUINI2108-W4 | W | QUINI2108 | 2021 | 3 | MAMD | 9a9a |
| QUINI2108-W5 | W | QUINI2108 | 2021 | 3 | MAMD | 9a9a |
| QUINI2109-M1 | M | QUINI2109 | 2021 | 4 | MA | 9a9a |
| QUINI2109-M3 | M | QUINI2109 | 2021 | 4 | MA | 9a9a |
| QUINI2109-M5 | M | QUINI2109 | 2021 | 4 | MA | 9a9a |
| QUINI2109-W1 | W | QUINI2109 | 2021 | 3 | MAMA | 9a9a |
| QUINI2109-W2 | W | QUINI2109 | 2021 | 3 | MAMA | 9a9a |
| QUINI2109-W4 | W | QUINI2109 | 2021 | 3 | MAMA | 9a9a |
| QUINI2109-W5 | W | QUINI2109 | 2021 | 3 | MAMA | 9a9a |
| QUINI2110-M1 | M | QUINI2110 | 2021 | 4 | MA | 9a9a |
| QUINI2110-M2 | M | QUINI2110 | 2021 | 4 | MA | 9a9a |
| QUINI2110-M3 | M | QUINI2110 | 2021 | 4 | MA | 9a9a |
| QUINI2110-M4 | M | QUINI2110 | 2021 | 4 | MA | 9a9a |
| QUINI2110-M5 | M | QUINI2110 | 2021 | 4 | MA | 9a9a |
| QUINI2110-M6 | M | QUINI2110 | 2021 | 4 | MD | 9a9a |
| QUINI2110-M7 | M | QUINI2110 | 2021 | 4 | MD | 9a9a |
| QUINI2110-W1 | W | QUINI2110 | 2021 | 3 | MAMD | 9a9a |
| QUINI2110-W2 | W | QUINI2110 | 2021 | 3 | MAMD | 9a9a |
| QUINI2110-W3 | W | QUINI2110 | 2021 | 3 | MAMD | 9a9a |
| QUINI2110-W4 | W | QUINI2110 | 2021 | 3 | MAMA | 9a9a |
| QUINI2110-W5 | W | QUINI2110 | 2021 | 3 | MAMD | 9a9a |
| QUINI2111-M1 | M | QUINI2111 | 2021 | 4 | MA | 9a9a |
| QUINI2111-M2 | M | QUINI2111 | 2021 | 4 | MA | 9a9a |
| QUINI2111-M4 | M | QUINI2111 | 2021 | 4 | MA | 9a9a |
| QUINI2111-M5 | M | QUINI2111 | 2021 | 4 | MA | 9a9a |
| QUINI2111-M6 | M | QUINI2111 | 2021 | 4 | MA | 9a9a |
| QUINI2111-M7 | M | QUINI2111 | 2021 | 4 | MA | 9a9a |
| QUINI2111-M8 | M | QUINI2111 | 2021 | 4 | MA | 9a9a |
| QUINI2111-W1 | W | QUINI2111 | 2021 | 3 | MAMA | 9a9a |

| | | | | | | |
|---|---|---|---|---|---|---|
| QUINI2111-W2 | W | QUINI2111 | 2021 | 3 | MAMA | 9a9a |
| QUINI2111-W3 | W | QUINI2111 | 2021 | 3 | MAMA | 9a9a |
| QUINI2111-W4 | W | QUINI2111 | 2021 | 3 | MAMA | 9a9a |
| QUINI2111-W5 | W | QUINI2111 | 2021 | 3 | MAMA | 9a9a |
| QUINI2112-M1 | M | QUINI2112 | 2021 | 4 | MA | 9a9a |
| QUINI2112-M2 | M | QUINI2112 | 2021 | 4 | MA | 9a9a |
| QUINI2112-M3 | M | QUINI2112 | 2021 | 4 | MA | 9a9a |
| QUINI2112-M4 | M | QUINI2112 | 2021 | 4 | MA | 9a9a |
| QUINI2112-M5 | M | QUINI2112 | 2021 | 4 | MA | 9a9a |
| QUINI2112-M6 | M | QUINI2112 | 2021 | 4 | MA | 9a9a |
| QUINI2112-M7 | M | QUINI2112 | 2021 | 4 | MA | 9a9a |
| QUINI2112-M8 | M | QUINI2112 | 2021 | 4 | MA | 9a9a |
| QUINI2112-W1 | W | QUINI2112 | 2021 | 3 | MAMA | 9a9a |
| QUINI2112-W2 | W | QUINI2112 | 2021 | 3 | MAMA | 9a9a |
| QUINI2112-W3 | W | QUINI2112 | 2021 | 3 | MAMA | 9a9a |
| QUINI2112-W4 | W | QUINI2112 | 2021 | 3 | MAMA | 9a9a |
| QUINI2112-W5 | W | QUINI2112 | 2021 | 3 | MAMA | 9a9a |
| QUINI2113-M1 | M | QUINI2113 | 2021 | 4 | MA | 9a9a |
| QUINI2113-M2 | M | QUINI2113 | 2021 | 4 | MA | 9a9a |
| QUINI2113-M3 | M | QUINI2113 | 2021 | 4 | MA | 9a9a |
| QUINI2113-M4 | M | QUINI2113 | 2021 | 4 | MD | 9a9a |
| QUINI2113-M5 | M | QUINI2113 | 2021 | 4 | MA | 9a9a |
| QUINI2113-M6 | M | QUINI2113 | 2021 | 4 | MA | 9a9a |
| QUINI2113-M7 | M | QUINI2113 | 2021 | 4 | MA | 9a9a |
| QUINI2113-M8 | M | QUINI2113 | 2021 | 4 | MA | 9a9a |
| QUINI2113-W1 | W | QUINI2113 | 2021 | 3 | MAMD | 9a9a |
| QUINI2113-W2 | W | QUINI2113 | 2021 | 3 | MAMA | 9a9a |
| QUINI2113-W3 | W | QUINI2113 | 2021 | 3 | MAMD | 9a9a |
| QUINI2113-W4 | W | QUINI2113 | 2021 | 3 | MAMA | 9a9a |
| QUINI2113-W5 | W | QUINI2113 | 2021 | 3 | MAMD | 9a9a |
| QUINI2114-W1 | W | QUINI2114 | 2021 | 3 | MAMA | 9a9a |
| QUINI2114-W2 | W | QUINI2114 | 2021 | 3 | MAMA | 9a9a |
| QUINI2114-W3 | W | QUINI2114 | 2021 | 3 | MAMA | 9a9a |
| QUINI2114-W4 | W | QUINI2114 | 2021 | 3 | MAMD | 9a9a |
| QUINI2114-W5 | W | QUINI2114 | 2021 | 3 | MAMD | 9a9a |
| QUINI2115-W1 | W | QUINI2115 | 2021 | 3 | MAMD | 9a9a |
| QUINI2115-W2 | W | QUINI2115 | 2021 | 3 | MAMD | 9a9a |
| QUINI2115-W3 | W | QUINI2115 | 2021 | 3 | MAMA | 9a9a |
| QUINI2115-W4 | W | QUINI2115 | 2021 | 3 | MAMA | 9a9a |
| QUINI2115-W5 | W | QUINI2115 | 2021 | 3 | MAMA | 9a9a |
| QUINI2116-M1 | M | QUINI2116 | 2021 | 4 | MA | 9a9a |
| QUINI2116-M2 | M | QUINI2116 | 2021 | 4 | MA | 9a9a |
| QUINI2116-M3 | M | QUINI2116 | 2021 | 4 | MA | 9a9a |
| QUINI2116-M4 | M | QUINI2116 | 2021 | 4 | MA | 9a9a |

| | | | | | | |
|---|---|---|---|---|---|---|
| QUINI2116-M5 | M | QUINI2116 | 2021 | 4 | MA | 9a9a |
| QUINI2116-M6 | M | QUINI2116 | 2021 | 4 | MA | 9a9a |
| QUINI2116-W1 | W | QUINI2116 | 2021 | 3 | MAMD | 9a9a |
| QUINI2116-W2 | W | QUINI2116 | 2021 | 3 | MAMA | 9a9a |
| QUINI2116-W3 | W | QUINI2116 | 2021 | 3 | MAMA | 9a9a |
| QUINI2116-W4 | W | QUINI2116 | 2021 | 3 | MAMD | 9a9a |
| QUINI2116-W5 | W | QUINI2116 | 2021 | 3 | MAMD | 9a9a |
| QUINI2117-M1 | M | QUINI2117 | 2021 | 4 | MA | 9a9a |
| QUINI2117-M2 | M | QUINI2117 | 2021 | 4 | MA | 9a9a |
| QUINI2117-M3 | M | QUINI2117 | 2021 | 4 | MA | 9a9a |
| QUINI2117-M4 | M | QUINI2117 | 2021 | 4 | MA | 9a9a |
| QUINI2117-M5 | M | QUINI2117 | 2021 | 4 | MA | 9a9a |
| QUINI2117-W1 | W | QUINI2117 | 2021 | 3 | MAMA | 9a9a |
| QUINI2117-W2 | W | QUINI2117 | 2021 | 3 | MAMA | 9a9a |
| QUINI2117-W4 | W | QUINI2117 | 2021 | 3 | MAMD | 9a9a |
| QUINI2117-W5 | W | QUINI2117 | 2021 | 3 | MAMA | 9a9a |
| QUINM220-G1 | G | QUINM220 | 2020 | 2 | MAP2 | 9a9r |
| QUINM220-G2 | G | QUINM220 | 2020 | 2 | MAP2 | 9a9r |
| QUINM220-M1 | M | QUINM220 | 2020 | 2 | P2 | 9r9r |
| QUINM220-M2 | M | QUINM220 | 2020 | 2 | P2 | 9r9r |
| QUINM220-M3 | M | QUINM220 | 2020 | 2 | P2 | 9r9r |
| QUINM220-M4 | M | QUINM220 | 2020 | 2 | P2 | unkn |
| QUINM220-M5 | M | QUINM220 | 2020 | 2 | P2 | 9r9r |
| QUINM220-M6 | M | QUINM220 | 2020 | 2 | P2 | 9r9r |
| QUINM220-M7 | M | QUINM220 | 2020 | 2 | P2 | 9r9r |
| QUINM220-M8 | M | QUINM220 | 2020 | 2 | P2 | 9r9r |
| QUIN-MQ3 | MQ | QUIN | 2019 | 4 | P1P2 | 9a9r |
| QUIN-MQ4 | MQ | QUIN | 2019 | 4 | P1P2 | 9a9r |
| RUM2101-M1 | M | RUM2101 | 2021 | 4 | MA | 9a9a |
| RUM2101-M2 | M | RUM2101 | 2021 | 4 | MA | 9a9a |
| RUM2101-M3 | M | RUM2101 | 2021 | 4 | MA | 9a9a |
| RUM2101-M4 | M | RUM2101 | 2021 | 4 | MA | 9a9a |
| RUM2101-M5 | M | RUM2101 | 2021 | 4 | MA | 9a9a |
| RUM2101-M6 | M | RUM2101 | 2021 | 4 | MA | 9a9a |
| RUM2101-M7 | M | RUM2101 | 2021 | 4 | MA | 9a9a |
| RUM2101-M8 | M | RUM2101 | 2021 | 3 | MA | 9a9a |
| RUM2101-W1 | W | RUM2101 | 2021 | 4 | MAMA | 9a9a |
| RUM2101-W2 | W | RUM2101 | 2021 | 4 | MAMA | 9a9a |
| RUM2101-W3 | W | RUM2101 | 2021 | 4 | MAMA | 9a9a |
| RUM2101-W4 | W | RUM2101 | 2021 | 4 | MAMA | 9a9a |
| RUM2101-W5 | W | RUM2101 | 2021 | 3 | MAMA | 9a9a |
| RUM2102-W1 | W | RUM2102 | 2021 | 4 | MAMA | 9a9a |
| RUM2102-W2 | W | RUM2102 | 2021 | 4 | MAMA | 9a9a |
| RUM2102-W3 | W | ? | 2021 | 4 | MAMA | 9a9a |

| | | | | | | |
|---|---|---|---|---|---|---|
| RUM2102-W3 | W | RUM2102 | 2021 | 4 | MAMA | 9a9a |
| RUM2102-W4 | W | RUM2102 | 2021 | 4 | MAMA | 9a9a |
| RUM2102-W5 | W | RUM2102 | 2021 | 4 | MAMA | 9a9a |
| RUM2103-M1 | M | RUM2103 | 2021 | 4 | MA | 9a9a |
| RUM2103-M2 | M | RUM2103 | 2021 | 4 | MA | 9a9a |
| RUM2103-M3 | M | RUM2103 | 2021 | 4 | MAMA | 9a9a |
| RUM2103-M4 | M | RUM2103 | 2021 | 4 | MA | 9a9a |
| RUM2103-M5 | M | RUM2103 | 2021 | 4 | MA | 9a9a |
| RUM2103-M6 | M | RUM2103 | 2021 | 4 | MA | 9a9a |
| RUM2103-W2 | W | RUM2103 | 2021 | 4 | MAMA | 9a9a |
| RUM2103-W3 | W | RUM2103 | 2021 | 4 | MAMA | 9a9a |
| RUM2103-W4 | W | RUM2103 | 2021 | 4 | MAMA | 9a9a |
| RUM2103-W5 | W | RUM2103 | 2021 | 4 | MAMA | 9a9a |
| RUM2104-G1 | G | RUM2104 | 2021 | 4 | MAMD | 9a9a |
| RUM2104-G2 | G | RUM2104 | 2021 | 4 | MAMA | 9a9a |
| RUM2104-G3 | G | RUM2104 | 2021 | 4 | MAMA | 9a9a |
| RUM2104-G4 | G | RUM2104 | 2021 | 4 | MAMD | 9a9a |
| RUM2104-G5 | G | RUM2104 | 2021 | 4 | MAMA | 9a9a |
| RUM2104-G6 | G | RUM2104 | 2021 | 4 | MAMA | 9a9a |
| RUM2104-G7 | G | RUM2104 | 2021 | 4 | MAMA | 9a9a |
| RUM2104-G8 | G | RUM2104 | 2021 | 3 | MAMA | 9a9a |
| RUM2104-W1 | W | RUM2104 | 2021 | 4 | MAMA | 9a9a |
| RUM2104-W2 | W | RUM2104 | 2021 | 4 | MAMD | 9a9a |
| RUM2104-W3 | W | RUM2104 | 2021 | 4 | MAMD | 9a9a |
| RUM2104-W4 | W | RUM2104 | 2021 | 4 | MAMD | 9a9a |
| RUM2104-W5 | W | RUM2104 | 2021 | 4 | MAMA | 9a9a |
| RUM2105-G1 | G | RUM2105 | 2021 | 4 | MAMD | 9a9a |
| RUM2105-G2 | G | RUM2105 | 2021 | 4 | MAMA | 9a9a |
| RUM2105-G3 | G | RUM2105 | 2021 | 4 | MAMD | 9a9a |
| RUM2105-G4 | G | RUM2105 | 2021 | 4 | MAMA | 9a9a |
| RUM2105-G5 | G | RUM2105 | 2021 | 4 | MAMD | 9a9a |
| RUM2105-G6 | G | RUM2105 | 2021 | 4 | MAMA | 9a9a |
| RUM2105-G7 | G | RUM2105 | 2021 | 4 | MAMD | 9a9a |
| RUM2105-G8 | G | RUM2105 | 2021 | 3 | MAMD | 9a9a |
| RUM2105-W1 | W | RUM2105 | 2021 | 4 | MAMA | 9a9a |
| RUM2105-W2 | W | RUM2105 | 2021 | 4 | MAMA | 9a9a |
| RUM2105-W4 | W | RUM2105 | 2021 | 4 | MAMA | 9a9a |
| RUM2105-W5 | W | RUM2105 | 2021 | 4 | MAMD | 9a9a |
| RUM2106-G1 | G | RUM2106 | 2021 | 4 | MAMA | 9a9a |
| RUM2106-G2 | G | RUM2106 | 2021 | 4 | MAMA | 9a9a |
| RUM2106-G3 | G | RUM2106 | 2021 | 3 | MAMA | 9a9a |
| RUM2106-W1 | W | RUM2106 | 2021 | 4 | MAMA | 9a9a |
| RUM2106-W2 | W | RUM2106 | 2021 | 4 | MAMA | 9a9a |
| RUM2106-W3 | W | RUM2106 | 2021 | 4 | MAMA | 9a9a |

| | | | | | | |
|---|---|---|---|---|---|---|
| RUM2106-W4 | W | RUM2106 | 2021 | 4 | MAMA | 9a9a |
| RUM2106-W5 | W | RUM2106 | 2021 | 2 | MAMA | 9a9a |
| SV19G2-W1 | W | SV19G2 | 2019 | 2 | MAMD | 9a9a |
| SV19G2-W2 | W | SV19G2 | 2019 | 2 | MAMD | 9a9a |
| SV19G2-W3 | W | SV19G2 | 2019 | 2 | MAMD | 9a9a |
| SV19G2-W4 | W | SV19G2 | 2019 | 2 | MAMD | 9a9a |
| SV19G2-W5 | W | SV19G2 | 2019 | 2 | MAMA | 9a9a |
| SV2001-M1 | M | SV2001 | 2020 | 2 | MA | 9a9a |
| SV2001-M2 | M | SV2001 | 2020 | 2 | MA | 9a9a |
| SV2001-M3 | M | SV2001 | 2020 | 2 | MA | 9a9a |
| SV2001-M4 | M | SV2001 | 2020 | 2 | MA | 9a9a |
| SV2001-M5 | M | SV2001 | 2020 | 2 | MA | 9a9a |
| SV2001-M6 | M | SV2001 | 2020 | 2 | MA | 9a9a |
| SV2001-M7 | M | SV2001 | 2020 | 2 | MA | 9a9a |
| SV2001-M8 | M | SV2001 | 2020 | 2 | MA | 9a9a |
| SV2001-W1 | W | SV2001 | 2020 | 2 | MAMA | 9a9a |
| SV2001-W2 | W | SV2001 | 2020 | 2 | MAMA | 9a9a |
| SV2001-W3 | W | SV2001 | 2020 | 2 | MAMA | 9a9a |
| SV2001-W4 | W | SV2001 | 2020 | 2 | MAMA | 9a9a |
| SV2001-W5 | W | SV2001 | 2020 | 2 | MAMA | 9a9a |
| SV2002-G1 | G | SV2002 | 2020 | 2 | MAP1 | 9a9a |
| SV2002-G2 | G | SV2002 | 2020 | 2 | MAP1 | 9a9a |
| SV2002-G3 | G | SV2002 | 2020 | 2 | MAP1 | 9a9a |
| SV2002-G4 | G | SV2002 | 2020 | 2 | P1P1 | 9a9a |
| SV2002-G5 | G | SV2002 | 2020 | 2 | MAP1 | 9a9a |
| SV2002-G6 | G | SV2002 | 2020 | 2 | MAP1 | 9a9a |
| SV2002-G7 | G | SV2002 | 2020 | 2 | MAP1 | 9a9a |
| SV2002-G8 | G | SV2002 | 2020 | 2 | MAP1 | 9a9a |
| SV2002-M1 | M | SV2002 | 2020 | 2 | P1 | 9a9a |
| SV2002-M12 | M | SV2002 | 2020 | 2 | P1 | 9a9a |
| SV2002-M2 | M | SV2002 | 2020 | 2 | P1 | 9a9a |
| SV2002-M3 | M | SV2002 | 2020 | 2 | P1 | 9a9a |
| SV2002-M4 | M | SV2002 | 2020 | 2 | P1 | 9a9a |
| SV2002-M5 | M | SV2002 | 2020 | 2 | P1 | 9a9a |
| SV2002-M6 | M | SV2002 | 2020 | 2 | P1 | 9a9a |
| SV2002-M7 | M | SV2002 | 2020 | 2 | P1 | 9a9a |
| SV2002-W5 | W | SV2002 | 2020 | 2 | MAP1 | 9a9a |
| SV2003-G1 | G | SV2003 | 2020 | 2 | P1P1 | 9a9a |
| SV2003-M1 | M | SV2003 | 2020 | 2 | P1 | 9a9a |
| SV2003-M2 | M | SV2003 | 2020 | 2 | P1 | 9a9a |
| SV2003-W1 | W | SV2003 | 2020 | 2 | P1P1 | 9a9a |
| SV2003-W2 | W | SV2003 | 2020 | 2 | MDP1 | 9a9a |
| SV2003-W3 | W | SV2003 | 2020 | 2 | MDP1 | 9a9a |
| SV2003-W4 | W | SV2003 | 2020 | 2 | MDP1 | 9a9a |

| | | | | | | |
|---|---|---|---|---|---|---|
| SV2003-W5 | W | SV2003 | 2020 | 4 | MAP1 | 9a9a |
| SV2101-G1 | G | SV2101 | 2021 | 4 | P1P2 | 9a9r |
| SV2101-M1 | M | SV2101 | 2021 | 4 | P2 | 9r9r |
| SV2101-M2 | M | SV2101 | 2021 | 4 | P2 | 9r9r |
| SV2101-M3 | M | SV2101 | 2021 | 4 | P2 | 9r9r |
| SV2101-M4 | M | SV2101 | 2021 | 3 | P2 | 9r9r |
| SV2101-W1 | W | SV2101 | 2021 | 3 | MAP1 | 9a9a |
| SV2101-W2 | W | SV2101 | 2021 | 3 | MAP2 | 9a9r |
| SV2101-W3 | W | SV2101 | 2021 | 3 | P1P2 | 9a9r |
| SV2101-W4 | W | SV2101 | 2021 | 3 | P1P2 | 9a9r |
| SV2101-W5 | W | SV2101 | 2021 | 4 | P1P2 | 9a9a |
| SV2102-M1 | M | SV2102 | 2021 | 4 | P2 | 9r9r |
| SV2102-M2 | M | SV2102 | 2021 | 4 | P2 | 9r9r |
| SV2102-M3 | M | SV2102 | 2021 | 4 | P2 | 9r9r |
| SV2102-M4 | M | SV2102 | 2021 | 4 | P2 | 9r9r |
| SV2102-M5 | M | SV2102 | 2021 | 4 | P2 | 9r9r |
| SV2102-M6 | M | SV2102 | 2021 | 4 | P2 | 9r9r |
| SV2102-M7 | M | SV2102 | 2021 | 4 | P2 | 9r9r |
| SV2102-M8 | M | SV2102 | 2021 | 4 | P2 | 9r9r |
| SV2102-W1g | W | SV2102 | 2021 | 4 | MAP1 | 9a9a |
| SV2102-W2g | W | SV2102 | 2021 | 4 | P1P2 | 9a9r |
| SV2102-W3g | W | SV2102 | 2021 | 3 | P1P2 | 9a9a |
| SV2102-W4 | W | SV2102 | 2021 | 4 | P1P2 | 9a9r |
| SV2102-W4g | W | SV2102 | 2021 | 4 | P1P2 | 9a9a |
| SV2102-W5g | W | SV2102 | 2021 | 4 | P1P2 | 9a9r |
| SV2103-M1 | M | SV2103 | 2021 | 4 | P1 | 9a9a |
| SV2103-M3 | M | SV2103 | 2021 | 4 | P1 | 9a9a |
| SV2103-M4 | M | SV2103 | 2021 | 4 | P1 | 9a9a |
| SV2103-M5 | M | SV2103 | 2021 | 4 | P1 | 9a9a |
| SV2103-M6 | M | SV2103 | 2021 | 4 | P1 | 9a9a |
| SV2103-M7 | M | SV2103 | 2021 | 3 | P1 | 9a9a |
| SV2103-W1 | W | SV2103 | 2021 | 3 | MAP1 | 9a9a |
| SV2103-W2 | W | SV2103 | 2021 | 3 | MAP1 | 9a9a |
| SV2103-W3 | W | SV2103 | 2021 | 3 | MAP1 | 9a9a |
| SV2103-W5 | W | SV2103 | 2021 | 3 | MAP1 | 9a9a |
| SV2104-M1 | M | SV2104 | 2021 | 3 | P1 | 9a9a |
| SV2104-M3 | M | SV2104 | 2021 | 3 | P1 | 9a9a |
| SV2104-M4 | M | SV2104 | 2021 | 3 | P1 | 9a9a |
| SV2104-M5 | M | SV2104 | 2021 | 3 | P1 | 9a9a |
| SV2104-M6 | M | SV2104 | 2021 | 3 | P1 | 9a9a |
| SV2104-W1 | W | SV2104 | 2021 | 3 | MAP1 | 9a9a |
| SV2104-W2 | W | SV2104 | 2021 | 3 | MAP1 | 9a9a |
| SV2104-W3 | W | SV2104 | 2021 | 3 | MAP1 | 9a9a |
| SV2104-W4 | W | SV2104 | 2021 | 3 | MAP1 | 9a9a |

| | | | | | | |
|---|---|---|---|---|---|---|
| SV2104-W5 | W | SV2104 | 2021 | 4 | MAP1 | 9a9a |
| SV2105-M1 | M | SV2105 | 2021 | 4 | P1 | 9a9a |
| SV2105-M3 | M | SV2105 | 2021 | 4 | P1 | 9a9a |
| SV2105-M5 | M | SV2105 | 2021 | 4 | P1 | 9a9a |
| SV2105-W1 | W | SV2105 | 2021 | 4 | MAP1 | 9a9a |
| SV2105-W2 | W | SV2105 | 2021 | 4 | MAP1 | 9a9a |
| SV2105-W3 | W | SV2105 | 2021 | 4 | MAP1 | 9a9a |
| SV2105-W4 | W | SV2105 | 2021 | 4 | MAP1 | 9a9a |
| SV2105-W5 | W | SV2105 | 2021 | 4 | MAP1 | 9a9a |
| SV-Gyne1 | NQ | - | 2019 | 4 | MAMA | 9a9a |
| SV-Gyne2 | NQ | - | 2019 | 4 | MAMD | 9a9a |
| VAR2101-W2 | W | VAR2101 | 2021 | 4 | MAMA | 9a9a |
| VAR2101-W3 | W | VAR2101 | 2021 | 4 | MAMA | 9a9a |
| VAR2101-W4 | W | VAR2101 | 2021 | 4 | MAMA | 9a9a |
| VAR2101-W5 | W | VAR2101 | 2021 | 4 | MAMA | 9a9a |
